# Age-dependent downregulation of endothelial lamin A/C impairs mechanotransduction and promotes cardiovascular disease

**DOI:** 10.64898/2026.11.12.764634

**Authors:** Iñigo Ruiz-Polo de Lara, Anna Bar, Pilar Gonzalo, Alberto del Monte-Monge, María González-Amor, Ana Barettino, Sara Pascual, Carlos Torroja, Judith Alonso, María J. Andrés-Manzano, Janusz Pyka, Gema de Rivas Hidalgo, Zeinep Berkimbayeva, Manel Taurón, Stephane Heymans, Cristina Rodríguez, José Martínez-González, Beatriz Dorado, Jaime Millán, Stefan Chlopicki, Ignacio Benedicto, Vicente Andrés

## Abstract

**Aims:** Aging is a major risk factor for cardiovascular disease, but the underlying mechanisms remain incompletely understood. Here, we investigated age-associated changes in endothelial lamin A/C expression and the contribution of endothelial lamin A/C loss to cardiovascular dysfunction and disease.

**Methods and Results:** Aging was associated with reduced endothelial lamin A/C expression in human coronary arteries and mouse aorta and heart. We generated an inducible endothelial-specific *Lmna* knockout mouse model on an atheroprone *Ldlr*-null background (*Lmna*KO*^iEC^*), which exhibited a 14.6% reduction in median survival. Vascular and cardiac phenotypes were evaluated using vascular reactivity assays, magnetic resonance imaging, blood pressure measurements, echocardiography, histology, nitric oxide assays, and biochemical analyses. *Lmna*KO*^iEC^* mice exhibited increased atherosclerotic burden, impaired endothelium-dependent vasodilation, vascular hyperpermeability, arterial stiffening, and hypertension. They also displayed increased cardiac mass, elevated NT-proBNP levels, incipient myocardial fibrosis, and signs of increased cardiac filling pressures despite preserved systolic function, resembling early stages of heart failure with preserved ejection fraction (HFpEF). Cardiac proteomics revealed inflammatory, profibrotic, and metabolic remodeling, with enrichment of molecular signatures associated with human HFpEF. Mechanistically, aortic single-cell transcriptomics revealed endothelial activation and suppression of the flow-responsive KLF2–NOS3 pathway. Consistently, endothelial lamin A/C deficiency reduced aortic nitric oxide production *ex vivo,* while siRNA-mediated *LMNA* knockdown impaired shear stress-induced *KLF2* and *NOS3* expression in human aortic endothelial cells.

**Conclusion:** These findings identify age-associated endothelial lamin A/C downregulation as a previously unrecognized contributor to cardiovascular aging and disease, and suggest preservation of endothelial nuclear mechanotransduction as a potential therapeutic strategy.

**Translational Relevance:** Our findings identify age-associated endothelial lamin A/C downregulation as a previously unrecognized contributor to cardiovascular aging, linking impaired nuclear mechanotransduction to age-related endothelial dysfunction and cardiovascular pathologies, including atherosclerosis, hypertension, and cardiac alterations resembling early HFpEF. These results highlight endothelial lamin A/C and downstream flow-responsive KLF2-NOS3 signaling as potential targets for therapeutic strategies aimed at preserving endothelial function and mitigating age-related cardiovascular disease.

**Graphical Abstract:** 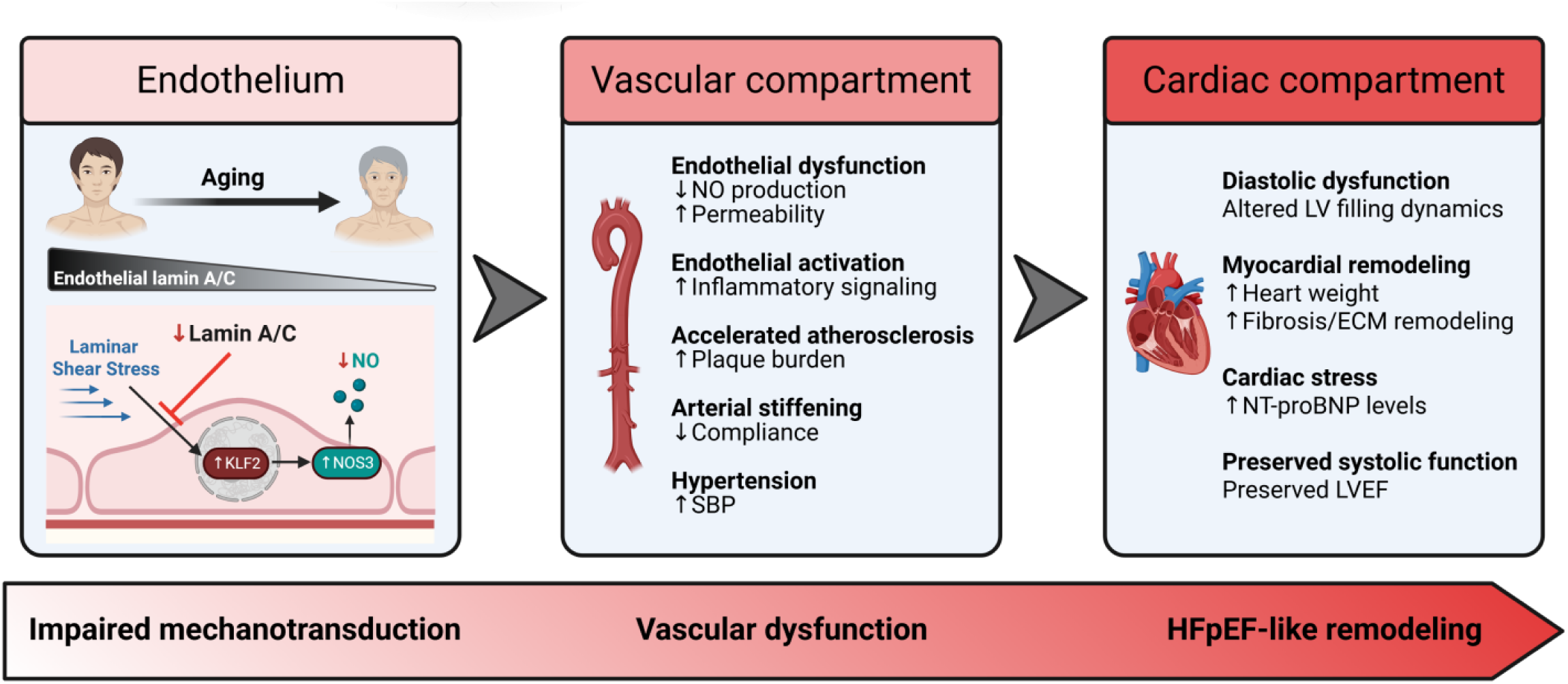

## Introduction

Aging is the strongest risk factor for cardiovascular disease (CVD), which remains the leading cause of mortality worldwide^1,2^. As life expectancy continues to increase, the burden of age-associated CVD is rising rapidly. Although this association is partly explained by cumulative exposure to traditional cardiovascular risk factors, clinical and epidemiological evidence indicates that aging itself independently promotes cardiovascular dysfunction and disease development^1,3–5^. These observations suggest the existence of age-dependent mechanisms that contribute directly to cardiovascular pathology. Identifying such mechanisms is therefore essential for developing therapeutic strategies to mitigate age-associated cardiovascular decline.

Endothelial cells (ECs) play a central role in maintaining vascular homeostasis by regulating vascular tone, permeability, inflammation, and thrombosis^6–8^. During aging, this homeostatic balance progressively deteriorates, leading to endothelial dysfunction characterized by reduced nitric oxide (NO) bioavailability, increased oxidative stress, and chronic low-grade inflammation^6–8^. Endothelial dysfunction is increasingly recognized not only as an early hallmark of cardiovascular aging but also as a key pathogenic mechanism linking aging to vascular and cardiac remodeling. Through impaired vascular homeostasis, endothelial dysfunction contributes to several age-related cardiovascular alterations, including arterial stiffening, hypertension, atherosclerosis, and heart failure with preserved ejection fraction (HFpEF)^3,9–13^.

A key determinant of endothelial homeostasis is the ability of ECs to sense and adapt to hemodynamic forces, particularly laminar shear stress (LSS)^14^. The conversion of these mechanical stimuli into adaptive transcriptional responses is mediated by specialized mechanotransduction pathways that couple extracellular forces to changes in gene expression. While mechanotransduction has historically been studied primarily through membrane-associated mechanosensors, increasing evidence indicates that the nucleus itself functions as a mechanosensitive organelle that integrates physical stimuli into adaptive transcriptional responses^15,16^.

This role of the nucleus in mechanotransduction directly implicates the nuclear lamina, a filamentous network underlying the nuclear inner membrane that provides structural support and regulates nuclear mechanics. Major components of the nuclear lamina are lamin A and lamin C (collectively termed lamin A/C), the ubiquitously expressed A-type lamin isoforms encoded by the *LMNA* gene^17^. Through interactions with the linker of nucleoskeleton and cytoskeleton (LINC) complex and chromatin, A-type lamins couple the cytoskeleton to the genome and play a central role in nuclear mechanotransduction^16^. Beyond their structural role, A-type lamins regulate chromatin organization, epigenetic states, gene expression programs, and cellular responses to stress, positioning them as key regulators of cellular adaptation to environmental cues^18^. Moreover, alterations in lamin A/C function underlie a wide spectrum of human diseases, collectively termed laminopathies, including premature aging syndromes and severe cardiovascular disorders, highlighting their fundamental importance in tissue homeostasis and cardiovascular function^19–21^. Accumulating evidence indicates that endothelial responses to mechanical forces become altered with aging, impairing adaptation to physiological flow patterns and contributing to age-related CVD^22^. While age-associated changes such as arterial stiffening and altered blood flow profiles modify vascular mechanics^22,23^, whether aging also induces cell-intrinsic defects in endothelial mechanotransduction remains poorly understood.

Here, we demonstrate that lamin A/C expression declines in ECs during physiological aging in both human and mouse tissues. To investigate the functional and pathological consequences of endothelial lamin A/C deficiency, we developed an inducible EC-specific *Lmna* knockout model (*Lmna*KO^iEC^). We hypothesized that age-associated downregulation of endothelial lamin A/C impairs nuclear mechanotransduction, disrupting shear stress-responsive signaling and ultimately promoting endothelial dysfunction and age-related CVD. To test this hypothesis, we combined comprehensive vascular and cardiac phenotyping with multi-omics analyses of cardiac and aortic tissues, and mechanobiological *in vitro* studies using human ECs exposed to LSS.

## Results

### Endothelial lamin A/C expression declines with aging in mouse and human tissues

To determine whether lamin A/C expression is altered during aging, we carried out flow cytometry experiments in aortas and hearts from young and aged wild-type mice **(Supplemental Figures S1 and S2)**. In the aorta, lamin A/C expression was significantly reduced with age in ECs (CD31⁺CD45⁻) but not in non-EC populations (CD31⁻) **(Figure 1A)**. In the heart, aging was associated with decreased lamin A/C expression in ECs (CD31⁺CD45⁻), leukocytes (CD31⁻CD45⁺), and non-EC/non-leukocyte populations (CD31⁻CD45⁻) **(Figure 1B)**. These findings demonstrate age-associated lamin A/C downregulation in mouse aortic and cardiac ECs.

**Figure 1.**
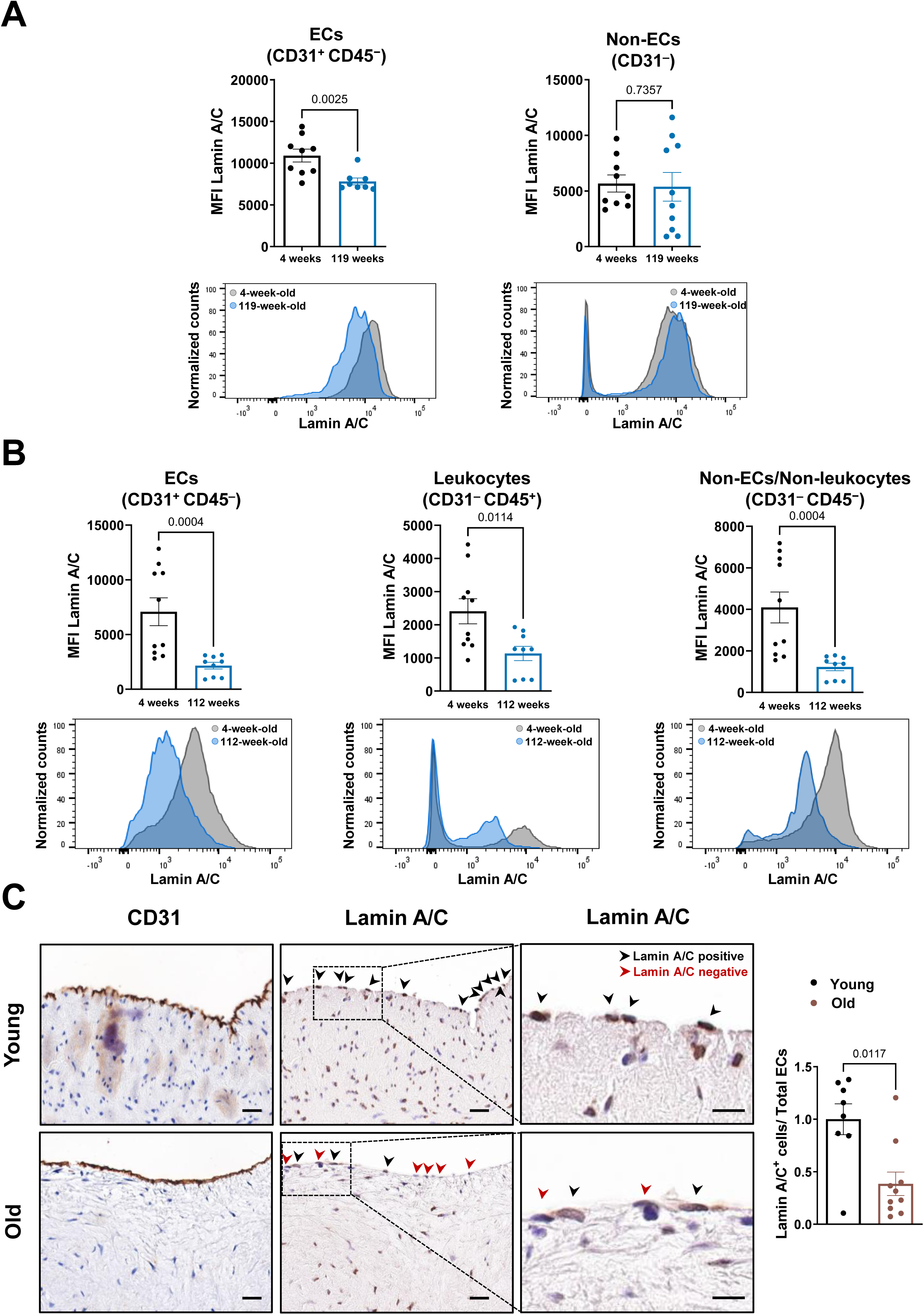
Endothelial lamin A/C expression declines during aging in mouse and human tissues. **(A)** Flow cytometry analysis of lamin A/C expression in endothelial cells (ECs; CD31⁺CD45⁻; young, n=9; aged, n=8) and non-ECs (CD31⁻; young, n=9; aged, n=10) isolated from aortas of young (4-week-old) and aged (119-week-old) wild-type mice. **(B)** Flow cytometry analysis of lamin A/C expression in ECs (CD31⁺CD45⁻), leukocytes (CD31⁻CD45⁺), and non-endothelial/non-leukocyte cells (CD31⁻CD45⁻) isolated from hearts of young (4-week-old) and aged (112-week-old) wild-type mice (young, n=10; aged, n=9). **(C)** Representative immunohistochemical staining for CD31 and lamin A/C in coronary arteries from young (23.2 ± 2.4 years; n=8) and aged (65.70 ± 0.98 years; n=10) individuals, counterstained with hematoxylin. Boxed regions are shown at higher magnification. Black arrowheads indicate lamin A/C-positive ECs while red arrowheads indicate lamin A/C-negative ECs. Scale bars, 20 μm (low magnification) and 10 μm (high magnification). Quantification shows the ratio of lamin A/C-positive EC nuclei relative to the total number of EC nuclei. Statistical analyses were performed using an unpaired t-test or Mann-Whitney U test in A and B, and Mann-Whitney U test in C. MFI: mean fluorescence intensity.

To assess the translational relevance of these findings, we next analyzed endothelial lamin A/C expression by immunohistochemistry, together with the endothelial marker CD31, in human coronary arteries from young (23.2 ± 2.4 years) and aged (65.70 ± 0.98 years) patients with end-stage dilated cardiomyopathy who underwent heart transplantation (see Methods). Aged coronary arteries exhibited a significant reduction in the proportion of lamin A/C⁺ ECs compared with young controls, indicating that endothelial lamin A/C expression declines with aging in human arteries **(Figure 1C)**.

### Generation and characterization of an inducible endothelial-specific lamin A/C-deficient mouse model

To investigate the functional consequences of endothelial lamin A/C loss across multiple cardiovascular phenotypes, including atherosclerosis, we generated an inducible endothelial-specific *Lmna* knockout model on an LDL receptor-deficient background (*Ldlr*⁻^/^⁻). *Lmna*^flox/flox^ mice^24^ were crossed with *Ldlr*⁻^/^⁻ mice harboring the tamoxifen-inducible *Cdh5Cre^ERT2^*transgene^25^ to generate *Lmna*^flox/flox^ *Ldlr*⁻^/^⁻ *Cdh5Cre^ERT2^*⁺^/^⁻ (hereafter referred to as *Lmna*KO*^iEC^*) and their Cre-negative *Lmna*^flox/flox^ *Ldlr*⁻^/^⁻ *Cdh5Cre^ERT2^*⁻^/^⁻ littermate controls. The efficiency and specificity of *Lmna* deletion in ECs were assessed at 16 weeks of age, 12 weeks after tamoxifen administration **(Figure 2A)**. RT-qPCR assays of sorted cells confirmed reduced *Lmna* expression in liver ECs (CD31⁺CD45⁻) from *LmnaKO^iEC^* mice compared with controls, whereas no significant changes were detected in the non-endothelial fraction (CD31⁻) **(Figure 2B)**. Consistently, immunofluorescence analysis of thoracic aorta cross-sections revealed a marked reduction in lamin A/C in CD31⁺ luminal ECs, while expression was preserved in non-endothelial cells **(Figure 2C)**. To quantify the extent of endothelial lamin A/C protein downregulation, we performed *en face* whole-mount immunofluorescence analysis of thoracic aortas for lamin A/C and the endothelial nuclear marker ETS-related gene (ERG) **(Figure 2D)**. This analysis confirmed a robust reduction in the percentage of lamin A/C⁺ endothelial nuclei in *Lmna*KO*^iEC^*aortas compared with controls **(Figure 2E)**. *Lmna*KO*^iEC^* mice were born at the expected Mendelian ratios and displayed normal fertility and viability. However, long-term longitudinal monitoring revealed lower body weight in *Lmna*KO*^iEC^* mice **(Figure 2F)** and a 14.6% reduction in median survival **(Figure 2G)**.

**Figure 2.**
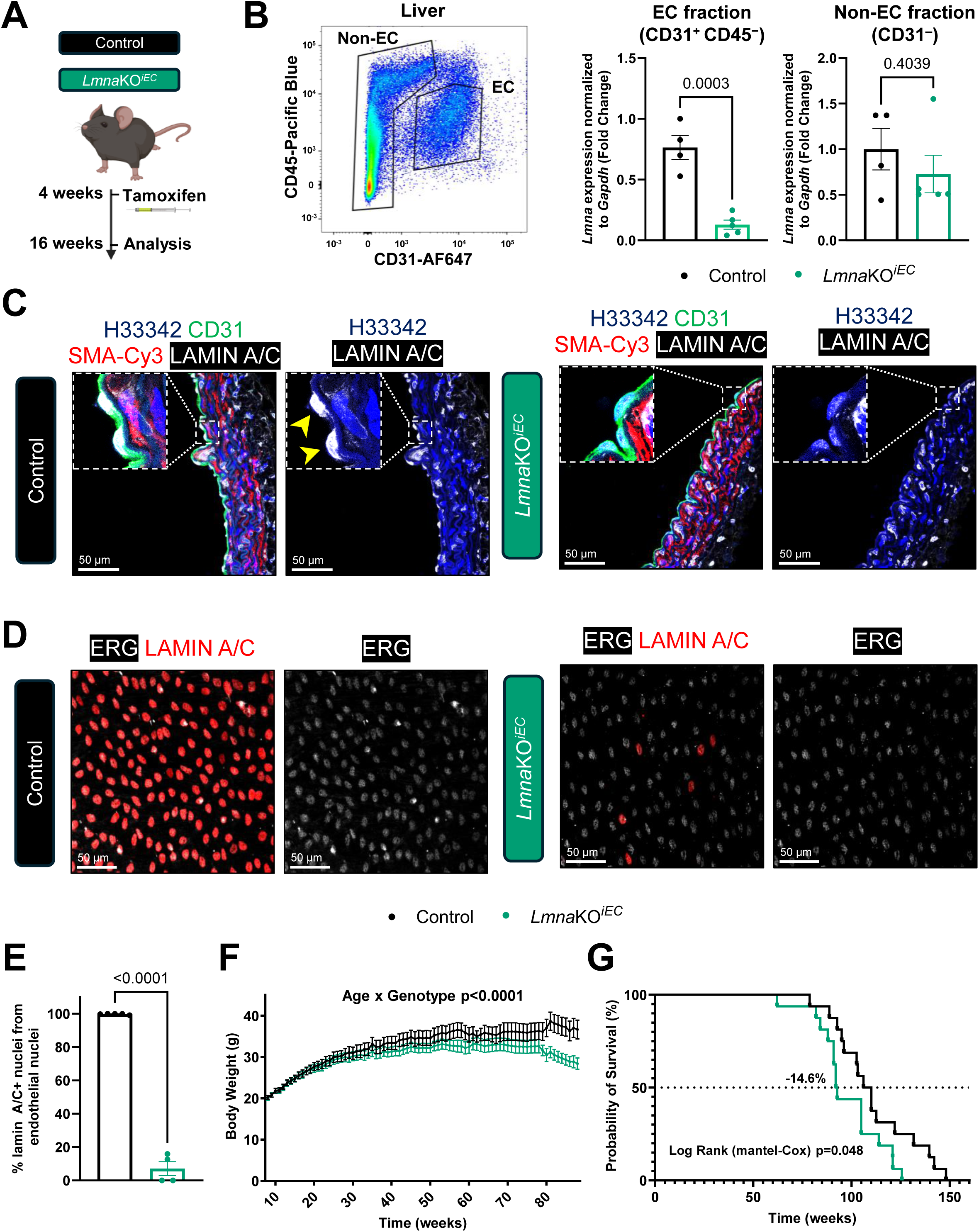
Efficient and specific endothelial lamin A/C deletion reduces body weight and survival in *Lmna*KO*^iEC^*mice. **(A)** Experimental design. *Lmna*KO*^iEC^* and control mice received tamoxifen at 4 weeks of age, and lamin A/C expression was analyzed at 16 weeks of age. **(B)** Liver endothelial cells (ECs; CD31⁺CD45⁻) and non-ECs (CD31⁻) were isolated by fluorescence-activated cell sorting, and *Lmna* expression was quantified by RT-qPCR (control, n=4; *Lmna*KO*^iEC^*, n=5). **(C)** Representative immunofluorescence images of transverse thoracic aorta sections showing lamin A/C (white), smooth muscle actin (SMA, red), ECs (CD31, green), and nuclei (Hoechst 33342, blue). **(D)** Representative *en face* immunofluorescence images of thoracic aortas showing lamin A/C (red) and ETS-related gene (ERG)-positive nuclei (white). **(E)** Quantification of lamin A/C-positive endothelial nuclei (ERG⁺) relative to the total number of endothelial nuclei (control, n=5; *Lmna*KO*^iEC^*, n=4). **(F)** Body weight curves of mice fed normal chow monitored weekly from 8 weeks of age (control, n=16; *Lmna*KO*^iEC^*, n=15). **(G)** Kaplan-Meier survival analysis of control and *Lmna*KO*^iEC^* mice. Median survival was 108.1 weeks in control mice (n=16) and 92.3 weeks in *Lmna*KO*^iEC^* mice (n=15). Statistical analyses were performed using an unpaired t-test in B and E, a mixed-effects two-way ANOVA with Greenhouse-Geisser correction in F, and the Mantel-Cox log-rank test in G.

### Endothelial lamin A/C deficiency promotes atherosclerosis and impairs endothelial function

To determine whether endothelial lamin A/C deficiency increases susceptibility to atherosclerosis, *Lmna*KO*^iEC^* and control mice were fed high fat-diet (HFD) for 12 weeks starting at 24 weeks of age. Despite comparable circulating lipid levels between genotypes **(Supplemental Figure S3),** *Lmna*KO*^iEC^* mice exhibited increased atherosclerotic burden, evidenced by a trend toward increased Oil Red-positive area in the aortas **(Figure 3A)** and significantly increased plaque area in the aortic root **(Figure 3B)**.

**Figure 3.**
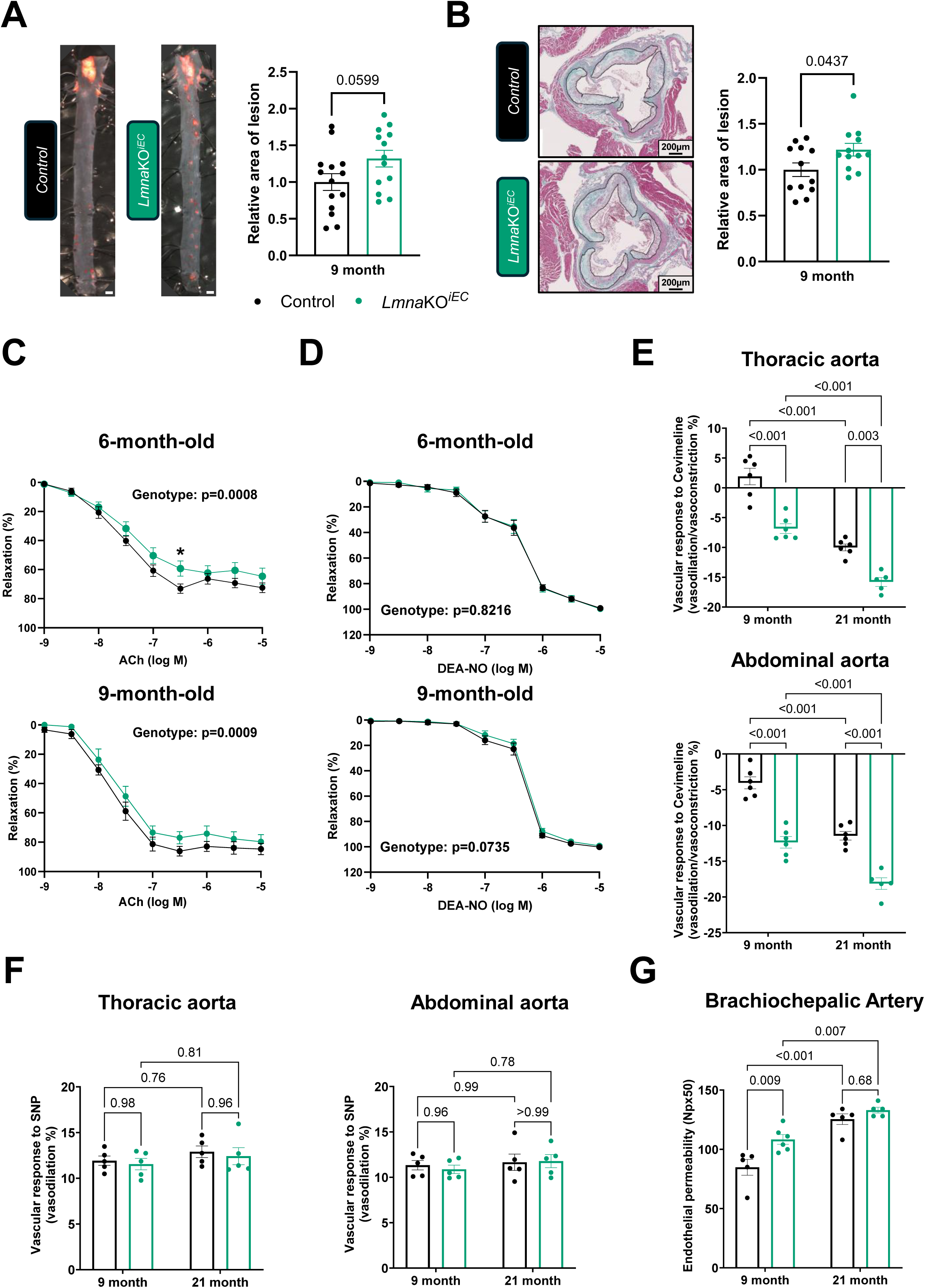
Endothelial lamin A/C deficiency promotes atherosclerosis, impairs nitric oxide-dependent vasodilation, and increases endothelial permeability. Mice were fed high-fat diet (HFD) for 12 weeks (A-D) or maintained on standard chow (E-G). **(A)** Representative Oil Red O-stained aortas and quantification of total plaque area expressed relative to the corresponding control group (control, n=14; *Lmna*KO*^iEC^*, n=13). Scale bars, 1 mm. **(B)** Representative Masson’s trichrome-stained transverse sections of the aortic root and quantification of plaque area expressed relative to the corresponding control group (n=12 per genotype). Atherosclerotic lesions are outlined in black. **(C)** Endothelium-dependent vasodilation in response to increasing concentrations of acetylcholine (ACh) measured by wire myography in thoracic aortic rings from control and *Lmna*KO*^iEC^*mice (6-month-old cohort: n=20; 9-month-old cohort: n=9). **(D)** Endothelium-independent relaxation in response to the nitric oxide (NO) donor DEA-NO (6-month-old cohort: n=20 per genotype; 9-month-old cohort: control, n=11; *Lmna*KO*^iEC^*, n=9). **(E)** MRI-based assessment of vascular responses to cevimeline in the thoracic and abdominal aorta. Responses are expressed as the percentage of vasodilation (positive values) or vasoconstriction (negative values) relative to baseline (9-month-old cohort: n=6 per genotype; 21-month-old cohort: control, n=6; *Lmna*KO*^iEC^*, n=5). (F) MRI-based assessment of vascular responses to sodium nitroprusside (SNP) in the thoracic and abdominal aorta. Responses are expressed as the percentage of vasodilation relative to baseline (9-month-old cohort: n=5; 21-month-old cohort: n=5). **(G)** MRI-based quantification of endothelial permeability in the brachiocephalic artery following administration of gadolinium-loaded liposomes. Endothelial permeability is expressed as the number of pixels within the region of interest showing a ≥50% increase in signal intensity (9-month-old cohort: control, n=5; *Lmna*KO*^iEC^*, n=6; 21-month-old cohort: n=5 per genotype). Statistical analyses were performed using an unpaired t-test in A and B, and two-way ANOVA in C-G. Asterisks in C indicate *P*<0.05 in the multiple-comparison analysis at each dose.

Given the established role of endothelial dysfunction in atherogenesis, we next assessed whether lamin A/C loss impairs endothelial function. *Ex vivo* wire myography of thoracic aortic rings from HFD-fed *Lmna*KO*^iEC^* mice revealed significantly reduced endothelium-dependent relaxation in response to acetylcholine at both 6 and 9 months of age **(Figure 3C)**. Notably, this impairment was comparable to that observed in aged wild-type mice, indicating that endothelial lamin A/C deficiency recapitulates an aging-like vascular phenotype **(Supplemental Figure S4A)**. In contrast, vasorelaxation induced by the NO donor diethylamine NONOate (DEA-NO) and contractile responses to phenylephrine were preserved in *Lmna*KO*^iEC^* mice **(Figure 3D and Supplemental Figure S4B)**, indicating intact vascular smooth muscle cell (VSMC) responsiveness and supporting an endothelial origin of the impaired vasoreactivity following EC-specific *Lmna* ablation.

To determine whether endothelial dysfunction also occurs *in vivo* and independently of HFD, we assessed aortic vasoreactivity by magnetic resonance imaging (MRI) in mice maintained on normal chow. At 9 months of age, *Lmna*KO*^iEC^* mice exhibited a switch from vasodilation to vasoconstriction in response to the muscarinic agonist cevimeline in the thoracic aorta, as well as enhanced vasoconstriction in the abdominal aorta, indicating impaired endothelium-dependent vasodilation **(Figure 3E)**. These alterations were further exacerbated in *Lmna*KO*^iEC^* mice at 21 months of age and were also observed, to a lesser extent, in aged control mice, suggesting accelerated age-dependent progression of endothelial dysfunction caused by endothelial lamin A/C deficiency **(Figure 3E)**. In contrast, vasodilatory responses to the endothelium-independent NO donor sodium nitroprusside were preserved across genotypes, further supporting intact VSMC responsiveness in *Lmna*KO*^iEC^* mice **(Figure 3F)**. Moreover, *in vivo* assessment of vascular permeability using gadolinium-labeled liposomes and MRI revealed increased endothelial permeability in the brachiocephalic artery of 9-month-old *Lmna*KO*^iEC^*mice compared with age-matched controls **(Figure 3G)**. Vascular permeability increased with aging in both groups, with no significant differences between genotypes at 21 months, indicating an accelerated increase in age-related endothelial permeability in *LmnaKO^iEC^*mice, with age-matched control mice reaching comparable levels by 21 months **(Figure 3G)**.

### Endothelial lamin A/C deficiency promotes features of cardiovascular aging

To further investigate the cardiovascular consequences of endothelial lamin A/C deficiency, we assessed vascular stiffness, blood pressure, and cardiac function in *Lmna*KO*^iEC^* and control mice. Pulse wave velocity (PWV), measured by Doppler imaging in mice maintained on standard diet, was significantly increased in the thoracic and abdominal aorta of 21-month-old *Lmna*KO*^iEC^* mice compared with age-matched controls, whereas no between-genotype differences were observed at 9 months of age **(Figure 4A)**. Although PWV increased with age in both genotypes, the greater increase observed in *Lmna*KO*^iEC^* mice indicates that endothelial lamin A/C deficiency accelerates age-related vascular stiffening **(Figure 4A)**. Moreover, *Lmna*KO*^iEC^* mice maintained on standard diet showed a trend toward increased systolic blood pressure (SBP) compared with controls, which reached statistical significance after 12 weeks of HFD **(Figure 4B)**.

**Figure 4.**
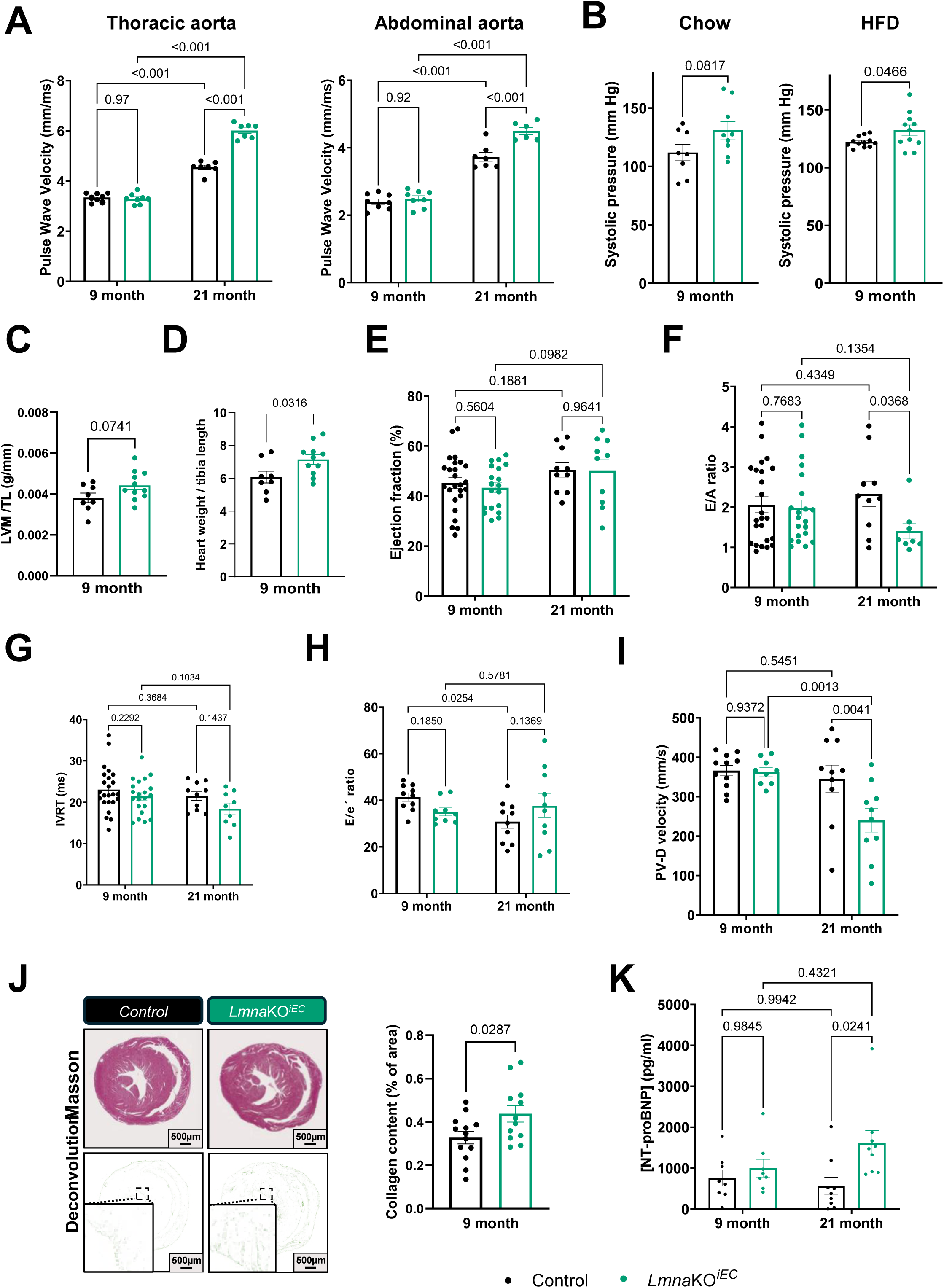
Endothelial lamin A/C deficiency increases vascular stiffness and promotes hypertension and diastolic dysfunction with preserved ejection fraction. Mice were fed standard chow in A and B or high-fat diet (HFD) for 12 weeks (9-month-old cohort) or 6 weeks (21-month-old cohort) in B-K. **(A)** Doppler-based measurement of pulse wave velocity in the thoracic and abdominal aorta (9-month-old cohort: n=8; 21-month-old cohort: n=7). **(B)** Systolic blood pressure (SBP) in 9-month-old mice maintained on standard chow (control, n=8; *Lmna*KO*^iEC^*, n=9) or HFD (control, n=12; *Lmna*KO*^iEC^*, n=11). **(C)** MRI-derived quantification of left ventricle mass, normalized to tibia length (control, n=8; *Lmna*KO*^iEC^*, n=11). **(D)** Heart weight normalized to tibia length (control, n=8; *Lmna*KO*^iEC^*, n=11). **(E-I)** Echocardiographic assessment of cardiac function in HFD-fed 9- and 21-month-old mice. Parameters analyzed included left ventricle ejection fraction (LVEF) (E), ratio of early-to-late transmitral flow velocity (E/A) (F), isovolumetric relaxation time (IVRT) (G), index of left ventricular filling pressure (E/e′) (H), and peak pulmonary venous diastolic flow velocity (PV-D) (I) (9-month-old cohort: control, n=25 (E), 24 (F), 24 (G), 10 (H), 11 (I); *Lmna*KO*^iEC^*, n=19 (E), 21 (F), 20 (G), 9 (H), 9 (I); 21-month-old cohort: control, n=10 (E-I); *Lmna*KO*^iEC^*, n=10 (E, H, I), 8 (F), 9 (G). **(J)** Representative Masson’s trichrome-stained heart cross-sections and color deconvolution used for collagen quantification in 9-month-old mice (control, n=13; *Lmna*KO*^iEC^*, n=12). **(K)** Serum N-terminal pro-B-type natriuretic peptide (NT-proBNP) concentrations measured by ELISA in 9- and 21-month-old mice (n=8 per genotype). Statistical analyses were performed using two-way ANOVA in A, D-H, and J, and an unpaired t-test in B, C, and I.

Given the increased vascular stiffness and systolic blood pressure observed in *Lmna*KO*^iEC^* mice, we next investigated whether endothelial lamin A/C deficiency was associated with cardiac structural and functional alterations under HFD-induced cardiometabolic stress. We performed transthoracic echocardiography in 9- and 21-month-old mice after 12 or 6 weeks of HFD, respectively. Cardiac morphology was largely preserved between genotypes at both ages, although *Lmna*KO*^iEC^* mice exhibited trends toward increased left ventricle (LV) posterior wall thickness, LV mass, and LV volume at 9 months of age **(Supplemental Figure S5)**. Cardiac cine-MRI analysis also revealed trends toward increased LV mass **(Figure 4C)** and enlarged LV end-diastolic and end-systolic volumes in *Lmna*KO*^iEC^* mice **(Supplemental Figure S6)**. Consistent with these observations, postmortem analysis demonstrated increased heart weight in *Lmna*KO*^iEC^* mice, suggestive of early cardiac remodeling **(Figure 4D)**.

Systolic function remained preserved in *Lmna*KO*^iEC^*mice, as evidenced by unchanged ejection fraction **(Figure 4E)** and cardiac output **(Supplemental Figure S7A)**. In contrast, *Lmna*KO*^iEC^*mice exhibited progressive alterations in diastolic function. Early transmitral filling velocity (E wave) was significantly increased at both 9 and 21 months of age, whereas atrial contraction velocity (A wave) increased predominantly at 21 months, resulting in a significantly reduced E/A ratio in 21-month-old *Lmna*KO*^iEC^* mice **(Figure 4F and Supplemental Figure S7A)**. Isovolumic relaxation time (IVRT) showed a nonsignificant trend toward shortening in *Lmna*KO*^iEC^* mice, particularly at 21 months of age **(Figure 4G)**, while early diastolic mitral annular velocity (e′) was increased rather than reduced **(Supplemental Figure S7B)**. The E/e′ ratio did not differ significantly between genotypes at either age, although it tended to increase in 21-month-old *Lmna*KO*^iEC^* mice relative to age-matched controls **(Figure 4H)**. The E/IVRT ratio was significantly elevated in *Lmna*KO*^iEC^* mice at both ages, suggesting increased ventricular filling pressures **(Supplemental Figure S7A)**. Furthermore, left atrial dimensions remained comparable between genotypes **(Supplemental Figure S7B)** and pulmonary venous flow analysis revealed a significant reduction in the diastolic component (PV-D) in 21-month-old *Lmna*KO*^iEC^*mice, whereas the systolic component (PV-S) remained unchanged **(Figure 4I and Supplemental Figure S7B)**. Together, these findings suggest early alterations in LV diastolic filling, particularly in aged *Lmna*KO*^iEC^* mice, with possible increases in filling pressures but without consistent evidence of impaired active myocardial relaxation.

To determine whether these functional abnormalities were accompanied by fibrotic remodeling, we analyzed collagen deposition in heart sections by Masson’s trichrome staining. Hearts from 9-month-old *Lmna*KO*^iEC^*mice fed HFD exhibited a mild but significant increase in collagen deposition compared with controls, indicative of incipient myocardial fibrosis **(Figure 4J)**. Circulating levels of NT-proBNP, a clinical biomarker of cardiac stress^26^, were unchanged at 9 months of age but significantly increased in 21-month-old *Lmna*KO*^iEC^* mice **(Figure 4K)**. Together with the observed alterations in LV diastolic filling, these findings support early cardiac remodeling and stress associated with endothelial lamin A/C deficiency, potentially involving increased passive myocardial stiffness and filling pressures.

### Cardiac proteomic remodeling in *Lmna*KO*^iEC^* mice shares molecular features with human HFpEF

To gain mechanistic insight into the cardiac phenotype associated with endothelial lamin A/C deficiency, we performed mass spectrometry-based proteomic analysis of whole hearts from 9-month-old *Lmna*KO*^iEC^* and control mice after 12 weeks of HFD. Differential expression analysis identified 132 upregulated and 96 downregulated proteins in *Lmna*KO*^iEC^* hearts **(Figure 5A)**. Pathway enrichment analysis revealed molecular signatures associated with endothelial dysfunction, chronic inflammation, and fibrotic remodeling, key features of HFpEF pathogenesis **(Figure 5B)**. *Lmna*KO*^iEC^*hearts exhibited marked alterations in regulators of vascular homeostasis, including reduced expression of NOS3 and AQP1 alongside increased levels of ANGPTL2 and DDAH1 **(Figure 5C)**. These alterations were accompanied by a robust inflammatory and stress-related signature, characterized by increased levels of inflammatory and apoptotic mediators, including STAT1, CASP1, CASP8, PTX3, IFI205, and LGALS3BP **(Figure 5C)**. Upstream regulator analysis predicted activation of inflammatory networks involving TNF, IFNG, IFNAR, IL1B, and the NLRP3 inflammasome, suggesting convergence of innate immune and interferon-driven pathways **(Figure 5D)**.

**Figure 5.**
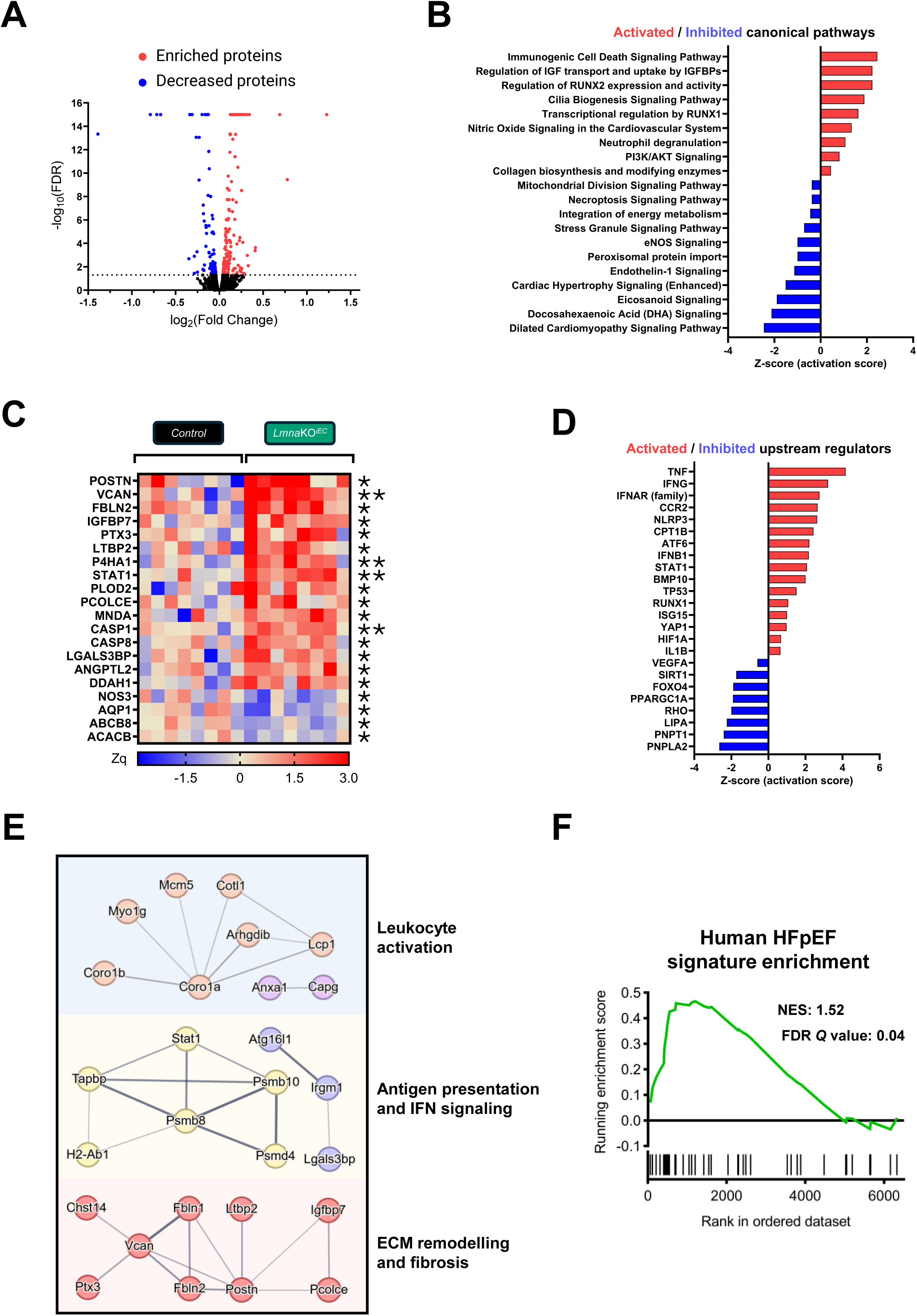
Cardiac proteomic profiling reveals molecular signatures consistent with HFpEF in *Lmna*KO*^iEC^*mice. Whole hearts from 9-month-old control and *Lmna*KO*^iEC^* mice fed a high-fat diet (HFD) for 12 weeks were analyzed by tandem mass tag-based quantitative proteomics (n=8). **(A)** Volcano plot showing fold changes and statistical significance of protein abundance differences between *Lmna*KO*^iEC^* and control hearts. The dotted line indicates FDR=0.05. For visualization purposes, all -log_10_(FDR) values >15 are displayed as Y=15. **(B)** Ingenuity pathway analysis activation *z*-scores of selected canonical pathways significantly altered after Benjamini-Hochberg correction (*P*<0.05). Activated and inhibited pathways are shown in red and blue, respectively. **(C)** Heatmap showing the relative abundance of representative proteins illustrating the major molecular alterations in *Lmna*KO*^iEC^*hearts. **(D)** Ingenuity pathway analysis activation *z*-scores of selected upstream regulators significantly altered after Benjamini-Hochberg correction (*P*<0.05). Activated and inhibited regulators are shown in red and in blue, respectively. **(E)** STRING protein-protein interaction network generated from significantly upregulated proteins in *Lmna*KO*^iEC^* hearts. Major functional clusters associated with leukocyte activation, antigen presentation/interferon signaling, and extracellular matrix (ECM) remodeling/fibrosis are highlighted. **(F)** Gene set enrichment analysis showing enrichment of a previously published human HFpEF proteomic signature^28^ in the cardiac proteome of *Lmna*KO*^iEC^* mice. FDR: false discovery rate. IGF: insulin-like growth factor. IGFBP: insulin-like growth factor binding protein. IFN: interferon. HFpEF: heart failure with preserved ejection fraction. *, *P*<0.05; **, *P*<0.01.

Pathways associated with extracellular matrix (ECM) organization and collagen biosynthesis were also significantly enriched in *Lmna*KO*^iEC^*hearts **(Figure 5B)**. This profibrotic signature was characterized by upregulation of structural and remodeling proteins involved in matrix expansion and tissue stiffening, including POSTN, VCAN, PLOD2, LTBP2, PCOLCE, P4HA1, and FBLN2 **(Figure 5C)**. Moreover, canonical pathway analysis predicted activation of RUNX1/2 transcriptional programs and insulin-like growth factor-binding protein (IGFBP) signaling, consistent with enhanced fibroblast activation **(Figure 5B).** Notably, expression of IGFBP7, a clinical biomarker associated with endothelial senescence and HFpEF remodeling^27^, was significantly elevated **(Figure 5C)**. Conversely, multiple metabolic and mitochondrial pathways were suppressed in *Lmna*KO*^iEC^*hearts **(Figure 5B)**, accompanied by reduced levels of proteins involved in mitochondrial homeostasis and fatty acid metabolism, including ABCB8 and ACACB **(Figure 5C)**. Upstream regulator analysis predicted inhibition of metabolic and stress-resilience programs involving SIRT1, FOXO4, and PPARGC1A, alongside altered PNPLA2 and LIPA signaling **(Figure 5D)**. Protein-protein interaction analysis using STRING further supported the coordinated nature of these molecular alterations. Functional clustering of significantly upregulated proteins identified three major interconnected modules associated with leukocyte activation, antigen presentation/interferon signaling, and ECM remodeling/fibrosis, supporting the presence of integrated inflammatory and profibrotic networks in *Lmna*KO*^iEC^*hearts **(Figure 5E)**.

To assess the translational relevance of these findings, we compared the *Lmna*KO*^iEC^* cardiac proteomic profile with a published proteomic dataset from hearts of control individuals and patients with HFpEF^28^. Gene set enrichment analysis revealed significant enrichment of the human HFpEF proteomic signature in *Lmna*KO*^iEC^* hearts, demonstrating substantial molecular overlap between cardiac remodeling induced by endothelial lamin A/C deficiency and human HFpEF **(Figure 5F)**. Altogether, these data indicate that endothelial lamin A/C deficiency is associated with coordinated inflammatory, profibrotic, metabolic, and vascular remodeling in the heart that closely resembles the molecular signature of human HFpEF.

### Aortic single-cell transcriptomic analysis reveals cellular and molecular alterations after endothelial lamin A/C depletion

To investigate the molecular mechanisms underlying the vascular phenotype induced by endothelial lamin A/C deficiency, we performed single-cell RNA sequencing (scRNA-seq) of aortas from 9-month-old *Lmna*KO*^iEC^*and control mice after 12 weeks of HFD **(Figure 6A)**. Following quality control and doublet removal, 15,756 cells were retained for downstream analysis, with a median of 3,426 detected genes per cell **(Supplemental Figure S8A)**. Unsupervised clustering identified 11 distinct cell populations (C0-C10) visualized by Uniform Manifold Approximation and Projection (UMAP) **(Figure 6B)**. Cell distributions were highly consistent across biological replicates, indicating good reproducibility and minimal batch effects **(Supplemental Figure S8B-F)**. Cluster identities were assigned based on the expression of established cell type-specific markers **(Figure 6C)**. Fibroblasts, characterized by *Pdgfra* and ECM-related genes including *Dcn* and *Fbln1*, represented the largest cell population (41.22%), followed by immune cells expressing the pan-hematopoietic marker *Ptprc* (CD45) (26.67%), VSMCs expressing canonical contractile markers including *Myh11*, *Acta2*, *Tagln*, and *Cnn1* (15.46%), and ECs expressing *Pecam1* (CD31), *Vwf*, and *Cdh5* (8.92%) **(Figure 6C)**.

**Figure 6.**
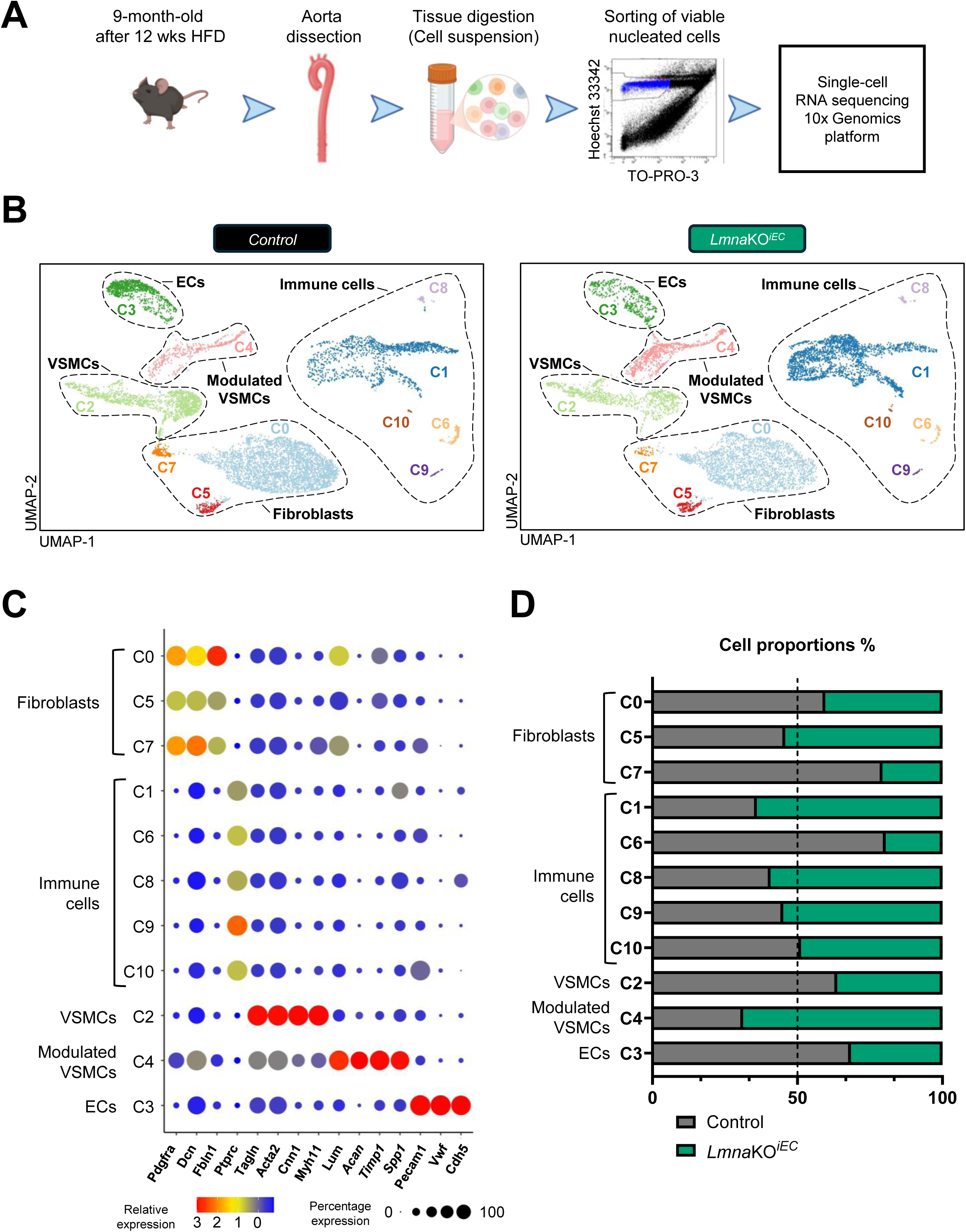
Single-cell RNA sequencing (scRNA-seq) analysis of aortic cells from *Lmna*KO*^iEC^* and control mice. Aortas from 9-month-old control and *Lmna*KO*^iEC^* mice fed high-fat diet (HFD) for 12 weeks were analyzed by scRNA-seq. **(A)** Experimental design. **(B)** Uniform manifold approximation and projection (UMAP) visualization of scRNA-seq data. Major aortic cell populations are indicated. **(C)** Dot plot showing the relative expression and the percentage of cells expressing canonical markers across the identified cell clusters. **(D)** Relative abundance of each cell cluster according to genotype. The dashed line indicates 50% proportion (i.e., an equal proportion of sequenced cells from each genotype within each cluster). EC: endothelial cell. VSMC: vascular smooth muscle cell.

In addition to these canonical vascular populations, we identified a distinct non-canonical cell cluster representing 7.73% of total cells, which we classified as modulated VSMCs. This population showed reduced expression of canonical VSMC markers and high expression of *Lum*, *Acan*, *Timp1*, and *Spp1*, genes associated with ECM remodeling and vascular pathology in atherosclerosis **(Figure 6C)**. Notably, modulated VSMCs were more abundant in *Lmna*KO*^iEC^* aortas, and differences in the relative abundance of several other cell populations were also observed between genotypes, consistent with extensive cellular remodeling following endothelial lamin A/C depletion **(Figure 6D)**. Given the endothelial focus of this study, we next performed a detailed analysis of reclustered ECs. The heterogeneity and transcriptional remodeling of fibroblast, VSMC, and immune cell populations are analyzed in the **Supplemental Results (Supplemental Figures S11-S22)** and **Supplemental Discussion.**

### Endothelial lamin A/C deficiency promotes endothelial activation and suppresses KLF2/4-NOS3 signaling

Unsupervised subclustering of *Pecam1* (CD31)-enriched aortic endothelial populations (cluster C3 in Figure 6B) revealed three transcriptionally distinct clusters (EC0-EC2), present in both control and *Lmna*KO^iEC^ mice **(Figure 7A)**. To further characterize each subcluster, we identified specifically enriched genes **(Figure 7B)** and assessed differences in their relative abundance **(Figure 7C)** between genotypes. EC0, the most abundant cluster, was enriched in genes associated with ECM organization, angiogenesis, and vascular homeostasis (e.g., *Spon1*, *Col4a3*, and *Calcrl*) (**Figure 7B**). EC1 was characterized by expression of genes linked to endothelial regulation and vascular homeostasis (e.g., *Sema3g*, *Sparcl1*, and *Adgrl4*) **(Figure 7B)**. In contrast, EC2 was enriched in genes associated with endothelial activation, inflammation, and ECM remodeling (e.g., *Vcam1*, *Icam1*, *Sele*, *Serpine1*, *Fn1*) **(Figure 7B)**. Notably, the relative abundance of activated EC2 was increased in *Lmna*KO*^iEC^*aortas, whereas EC0 and EC1 populations were reduced (**Figure 7C**). EC2 was also enriched in *Aqp1* **(Figure 7B)**, a marker highly expressed in ECs localized to the aortic arch^29^, a region exposed to disturbed and oscillatory blood flow. These findings suggest that endothelial lamin A/C deficiency promotes the acquisition of an activated endothelial phenotype associated with disturbed flow.

**Figure 7.**
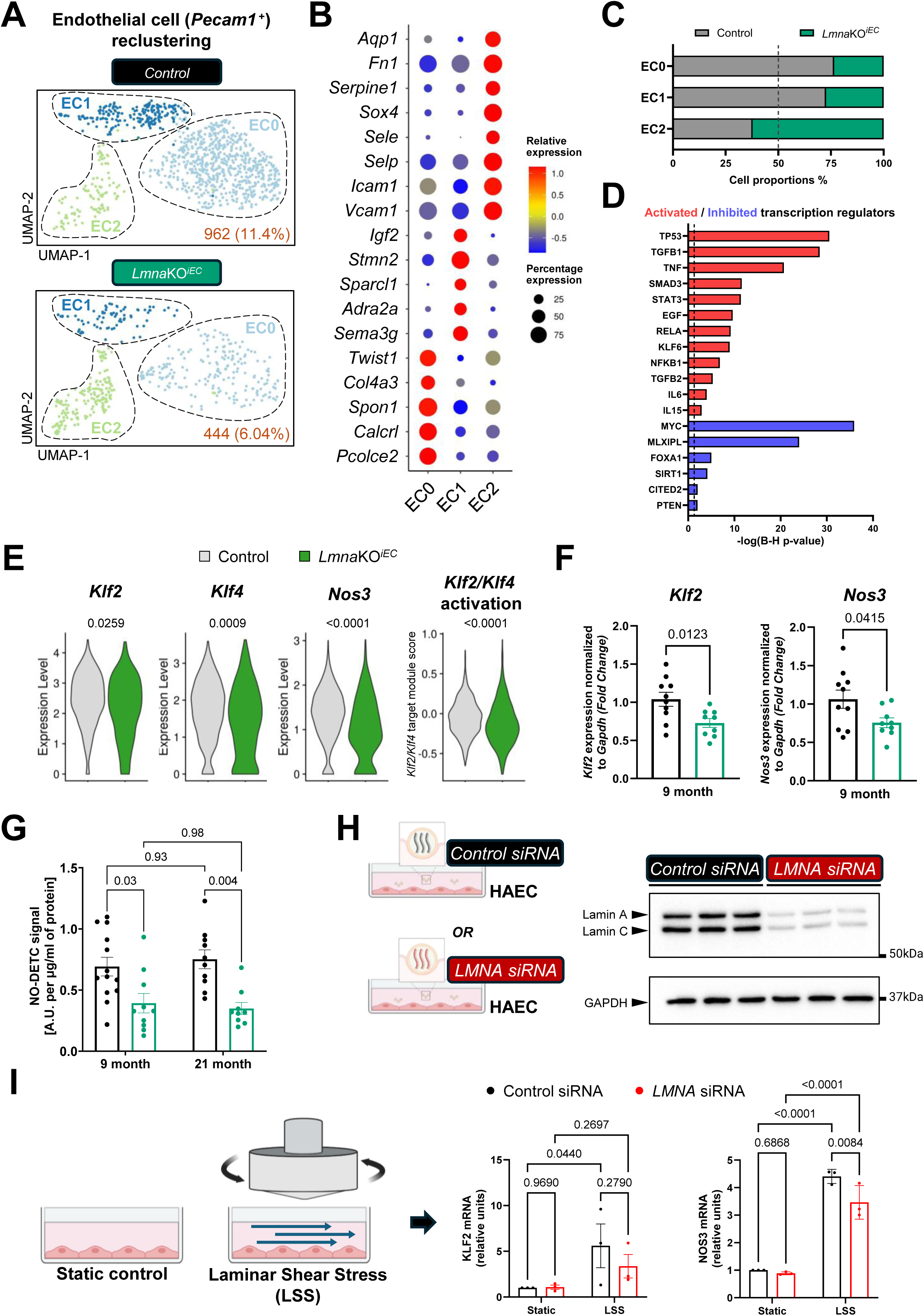
Endothelial lamin A/C deficiency drives inflammatory endothelial activation, reduces nitric oxide production, and impairs shear stress-induced *KLF2* and *NOS3* expression. (A) Uniform manifold approximation and projection (UMAP) visualization of endothelial cell (EC) reclustering in control and *Lmna*KO*^iEC^*mice. Absolute cell number and percentage of sequenced cells are indicated for each genotype (bottom right corner). **(B)** Dot plot showing the average expression and the percentage of cells expressing selected genes across different EC subclusters. **(C)** Cell proportions of each subcluster by genotype. The dashed line indicates 50% proportion (i.e., same number of sequenced cells in both genotypes for each cluster). **(D)** Selected upstream regulators predicted by Ingenuity pathway analysis to be activated (*z*-score>2; marked in red) or inhibited (*z*-score<-2; marked in blue) in *Lmna*KO*^iEC^* ECs. **(E)** Violin plots showing expression levels of key components of the laminar shear stress (LSS)/nitric oxide (NO) axis, as well as the predicted activation score of KLF2 and KLF4 in *Lmna*KO*^iEC^* and control ECs based on the expression of known downstream targets in *Lmna*KO*^iEC^* ECs. **(F)** RT-qPCR analysis of *Nos3* and *Klf2* expression in the aortic arch of 9-month-old control and *Lmna*KO*^iEC^* mice after 12 weeks of high-fat diet (HFD) (control, n=10; *Lmna*KO*^iEC^*, n=9). **(G)** Electron paramagnetic resonance *ex vivo* analysis of NO production by thoracic aortas in 9-month-old (control, n=13; *Lmna*KO*^iEC^*, n=10) and 21-month-old mice (control, n=10; *Lmna*KO*^iEC^*, n=9). NO levels are expressed as the intensity of NO-DETC signal and normalized to the total protein content of each sample. **(H)** Experimental approach to downregulate lamin A/C expression in human aortic endothelial cells (HAECs) and Western blot analysis of lamin A/C and GAPDH in HAECs transfected with control or *LMNA* siRNAs (n=3). Molecular weight markers are indicated on the right. **(I)** Strategy to induce laminar shear stress (LSS) using a viscometer in HAECs and RT-qPCR analysis of *NOS3 and KLF2* expression under static and LSS conditions in control and *LMNA* siRNA-treated cells (n=3). Statistical analyses were performed by Wilcoxon rank-sum test in E, unpaired t-test in F and two-way ANOVA in G and I. NO: nitric oxide. NO-DETC: nitric oxide-diethyldithiocarbamate complex. siRNA: small interfering RNA.

Ingenuity pathway analysis of the global endothelial population predicted activation of inflammatory, ECM remodeling, and mechanotransduction-related pathways in *Lmna*KO*^iEC^* aorta, consistent with an activated endothelial phenotype **(Supplemental Figure S9A)**. It also predicted activation of pro-inflammatory cytokines and transcriptional regulators, including TNF, IL6, IL15, and NF-κB1, as well as growth factors associated with inflammation and endothelial-to-mesenchymal transition, including TGFB1, TGFB2, and EGF **(Figure 7D)**. Conversely, homeostatic regulators involved in vascular quiescence and responses to shear stress, including SIRT1, PTEN, and FOXA1, were predicted to be inhibited **(Figure 7D)**. Consistent with these predictions, *Lmna*KO*^iEC^* ECs exhibited upregulation of inflammatory genes and reduced expression of genes associated with endothelial quiescence, including *Tek* (Tie2), *Cav1*, *Nrp1*, *Icam2*, *Ptgs1* (COX-1), and *Pecam1* **(Supplemental Figure S9B and S9C)**.

*Lmna*KO*^iEC^* ECs exhibited significant downregulation of *Klf2* and *Klf4*, LSS-responsive transcription factors in ECs, together with reduced expression of their downstream target *Nos3*, a critical mediator of endothelial NO production **(Figure 7E)**. To further assess the effects of endothelial lamin A/C depletion on KLF2/4-dependent transcription, we generated a composite score based on genes previously reported to be induced by KLF2/4^30^. This score was reduced in *Lmna*KO*^iEC^*ECs, indicating broad suppression of the KLF2/4-dependent vasoprotective transcriptional program **(Figure 7E)**. As expected, *Klf2* and *Nos3* were highly enriched in ECs compared with other aortic cell populations **(Supplemental Figure S10A and S10B)**, and their reduced expression in *Lmna*KO*^iEC^* mice was validated by RT-qPCR analysis of aortic arches from 9-month-old mice challenged with HFD **(Figure 7F)**. Together, these findings indicate that lamin A/C depletion in ECs impairs endothelial mechanotransduction-related transcriptional programs.

Given the critical role of NOS3 in endothelial NO production, we next determined whether reduced *Nos3* expression in *Lmna*KO*^iEC^* mice was accompanied by decreased vascular NO production. *Ex vivo* electron paramagnetic resonance analysis of thoracic aortas from 9- and 21-month-old mice revealed reduced NO production in *Lmna*KO*^iEC^* aortas at both ages (**Figure 7G**), providing a functional correlate of the impaired endothelium-dependent vasodilation observed in these mice (**Figures 3C and 3E**).

### Lamin A/C deficiency impairs the endothelial transcriptional response to laminar shear stress

To determine whether endothelial lamin A/C deficiency directly impairs shear stress-mediated upregulation of *NOS3*, we exposed cultured human aortic ECs (HAECs) to LSS after transient *LMNA* knockdown with siRNAs, which efficiently reduced lamin A/C protein levels **(Figure 7H)**. Under static conditions, lamin A/C knockdown did not affect *KLF2* or *NOS3* expression, as determined by RT-qPCR **(Figure 7I)**. Exposure to LSS significantly increased *KLF2* expression in control HAECs, whereas this increase did not reach statistical significance in *LMNA*-silenced cells. Although LSS induced *NOS3* expression in both groups, *NOS3* levels were significantly lower in *LMNA*-silenced HAECs than in control cells under LSS conditions **(Figure 7I)**. These findings demonstrate that lamin A/C deficiency attenuates the endothelial transcriptional response to laminar flow by impairing LSS-induced *KLF2* and *NOS3* expression.

## Discussion

The mechanisms linking aging to CVD remain incompletely understood. Here, we demonstrate that lamin A/C expression decreases in ECs during physiological aging in both mouse and human tissues, and that inducible endothelial-specific *Lmna* deletion in *Lmna*KO^iEC^ mice reduces median survival by 14.6% and is sufficient to promote multiple hallmarks of cardiovascular aging, including endothelial dysfunction, arterial stiffening, vascular permeability, hypertension, accelerated atherosclerosis, and HFpEF-like cardiac remodeling. Mechanistically, endothelial lamin A/C deficiency impairs flow-responsive signaling, resulting in suppression of the KLF2-NOS3 axis and reduced endothelial NO production. Together, these findings identify endothelial lamin A/C downregulation as a previously unrecognized mechanism contributing to cardiovascular aging and associated CVD.

Previous studies have reported age-associated downregulation of lamin A/C expression in multiple cell types, including cardiomyocytes, osteoblasts, and hematopoietic stem cells^31–34^. We recently showed that lamin A/C also declines with aging in mouse and human circulating leukocytes and demonstrated that this alteration contributes causally to atherosclerosis^35^. In addition, a comprehensive proteomic analysis across a 50-year human lifespan suggested that LMNA protein abundance decreases in the aging aorta^36^. However, whether lamin A/C declines during physiological aging in ECs and whether this alteration contributes functionally to CVD remained unknown. Our findings extend these observations by identifying the endothelium as another vascular compartment in which age-associated lamin A/C decline promotes cardiovascular aging. More broadly, they support the emerging concept that alterations of the nuclear lamina are active drivers, rather than merely consequences, of cardiovascular aging.

Endothelial dysfunction was one of the earliest and most consistent phenotypes observed in *Lmna*KO*^iEC^* mice. Although endothelial function declined progressively with aging in control animals, impairment of endothelium-dependent vasodilation was significantly accelerated by endothelial lamin A/C depletion. Preserved responses to exogenous NO donors demonstrated that impaired vasorelaxation resulted from endothelial dysfunction rather than altered VSMC responsiveness. Loss of endothelial homeostasis in *Lmna*KO*^iEC^* mice was further supported by reduced NOS3-dependent NO production and increased vascular permeability, both well-established features of endothelial dysfunction^6,11^. *Lmna*KO*^iEC^* mice also developed accelerated arterial stiffness and hypertension, two major manifestations of vascular aging that are closely linked to impaired NO availability and endothelial dysfunction^7,11^. Together, these findings indicate that endothelial lamin A/C deficiency promotes vascular aging by reducing NO-dependent vascular protection and impairing the ability of ECs to maintain vascular homeostasis.

Consistent with the central role of endothelial dysfunction in the initiation and progression of atherosclerosis^9,11^, endothelial-specific lamin A/C deficiency promoted atherosclerotic lesion development in fat-fed *Lmna*KO*^iEC^* mice. The reduction in NO bioavailability provides a plausible mechanistic link between endothelial lamin A/C deficiency and enhanced plaque formation. Consistent with this interpretation, lamin A/C-deficient aortic ECs exhibited upregulation of genes associated with endothelial activation and inflammation, including *Sox4*, which encodes a transcription factor previously linked to disturbed flow and atheroprone endothelial phenotypes^37^. Although the primary focus of this study was the endothelial compartment, scRNA-seq analysis revealed broader remodeling throughout the vascular wall of *Lmna*KO*^iEC^* mice, including increased abundance of activated macrophage subsets, foam cells, and modulated VSMCs (see **Supplemental Results and Supplemental Discussion**). Together, these findings support a model in which endothelial lamin A/C deficiency initiates vascular inflammation that propagates across multiple vascular cell populations, thereby creating a microenvironment that favors atherogenesis.

Endothelial lamin A/C deficiency was also associated with cardiac remodeling and alterations in LV diastolic filling, particularly in aged mice. *Lmna*KO^iEC^ mice exhibited reduced E/A ratio and pulmonary venous diastolic velocity, together with increased myocardial collagen deposition and elevated circulating NT-proBNP levels, while systolic function remained preserved. These findings suggest early diastolic abnormalities and cardiac wall stress, resembling several key features of human HFpEF^12,38^. However, the increased e′ velocity and trend toward shorter IVRT do not provide direct evidence of impaired active myocardial relaxation. Importantly, IVRT is influenced by both myocardial relaxation and left atrial pressure: impaired LV relaxation tends to prolong IVRT, whereas elevated left atrial pressure can shorten it, potentially exerting opposing effects on IVRT that mask impaired myocardial relaxation. Therefore, direct hemodynamic measurements would be required in future studies to establish increased ventricular stiffness or filling pressures, as performed in a recent study of physiological cardiac aging that also reported reduced E/A ratios without significant changes in E/e′^39^. These findings are particularly noteworthy given the growing recognition that endothelial dysfunction and microvascular abnormalities are central contributors to HFpEF pathogenesis^10,13^. HFpEF-like remodeling in *Lmna*KO^iEC^ mice is likely mediated by multiple mechanisms, including reduced NO bioavailability, hypertension, and chronic vascular inflammation, all of which are well-established contributors to myocardial remodeling and diastolic dysfunction^10,40^. Nevertheless, whether the observed changes arise primarily from increased systemic blood pressure and arterial stiffness or from local cardiac endothelial dysfunction remains unresolved.

Proteomic analyses of whole hearts provided strong molecular support for the HFpEF-like phenotype observed in *Lmna*KO*^iEC^*mice. Most notably, comparison with an independent proteomic dataset from patients with HFpEF^28^ revealed a significant enrichment of the human HFpEF molecular signature in *Lmna*KO^iEC^ hearts. Proteomic profiling revealed activation of inflammatory, profibrotic, and ECM remodeling pathways together with suppression of metabolic and mitochondrial programs, consistent with the molecular remodeling associated with HFpEF^28,40^. Increased ECM remodeling signatures and predicted activation of TGFβ signaling correlated well with the increased myocardial collagen deposition observed histologically, whereas enrichment of TNF, interferon, and inflammasome-related pathways was consistent with the growing recognition of chronic low-grade inflammation as a central contributor to both cardiovascular aging and HFpEF. Moreover, increased IGFBP7 expression in *Lmna*KO*^iEC^* mice was consistently detected in both whole-heart proteomic analyses and aortic ECs by scRNA-seq. Given the association of IGFBP7 with endothelial dysfunction, cardiac remodeling, and adverse outcomes in patients with HFpEF^27^, this cross-platform validation provides additional support for the relevance of endothelial lamin A/C deficiency to molecular processes underlying human HFpEF.

Mechanistically, our scRNA-seq analysis, NO production assays, and *in vitro* experiments strongly support impaired endothelial mechanotransduction as the central mechanism linking lamin A/C deficiency to cardiovascular dysfunction. ECs continuously sense hemodynamic forces, and LSS maintains endothelial homeostasis by activating transcriptional programs that preserve endothelial quiescence and vascular integrity^41^. Consistent with this concept, lamin A/C-deficient aortic ECs exhibited marked suppression of *Klf2*, *Klf4*, and *Nos3*, together with activation of inflammatory and cellular stress programs. KLF2 and KLF4 are master regulators of endothelial responses to LSS, orchestrating the expression of anti-inflammatory, anti-thrombotic, and vasoprotective genes, including *NOS3,* that are essential for vascular homeostasis^42–44^. Importantly, *LMNA* knockdown in HAECs impaired the induction of *KLF2* and *NOS3* under LSS, but not under static conditions, demonstrating that lamin A/C is required for the endothelial transcriptional response to blood flow rather than for basal gene expression. Previous studies showed that expression of mutant LMNA proteins impairs shear stress-mediated induction of NOS3^45,46^. Our findings extend these observations by demonstrating that, even in the absence of *LMNA* mutations, age-associated reduction of wild-type lamin A/C in ECs is sufficient to compromise flow-responsive endothelial programs, reduce NO bioavailability, and promote cardiovascular dysfunction. Interestingly, recent work in cardiomyocyte-specific *Lmna* knockout mice demonstrated that mechanically-induced nuclear envelope rupture contributes to inflammatory and fibrotic cardiac remodeling^47^, further highlighting the importance of lamin A/C in maintaining cellular responses to mechanical stress. Although the molecular events linking lamin A/C downregulation to defective mechanotransduction remain to be elucidated, our findings identify lamin A/C as a critical component of the machinery that enables ECs to translate physiological mechanical stimuli into vasoprotective transcriptional responses.

Recent studies have highlighted the important anti-atherogenic role of KLF2^48,49^. Moreover, statin-mediated KLF2 activation restores endothelial function in *LMNA*-mutant ECs *in vitro*^50^, raising the possibility that pharmacological restoration of endothelial mechanotransduction-related pathways may mitigate vascular dysfunction associated with lamin A/C downregulation during aging. In this context, it is noteworthy that statins have recently been associated with improved outcomes in patients with HFpEF, even in the absence of overt atherosclerotic disease^51^. In light of our findings, it will be important to determine whether these clinical benefits are mediated, at least in part, through restoration of flow-responsive endothelial programs impaired during vascular aging. More broadly, identifying the molecular mechanisms responsible for endothelial lamin A/C downregulation during physiological aging may uncover novel therapeutic strategies aimed at preserving endothelial mechanotransduction, maintaining vascular homeostasis, and delaying age-associated cardiovascular dysfunction driven by multiple mechanisms downstream of impaired NO bioavailability^8^.

In conclusion, this work identifies endothelial lamin A/C as a critical regulator of cardiovascular homeostasis. Our findings demonstrate that age-related endothelial lamin A/C decline impairs endothelial mechanotransduction, leading to endothelial dysfunction, vascular inflammation, arterial stiffening, atherosclerosis, and HFpEF-like cardiac remodeling. Together, these findings establish endothelial lamin A/C downregulation as a previously unrecognized molecular link between physiological vascular aging and CVD.

## Methods

### Ethical statement

All animal procedures complied with Directive 2010/63/EU and Recommendation 2007/526/EC of the European Union, implemented in Spanish legislation through Real Decreto 53/2013. All animal protocols were approved by the institutional ethics committees and the Animal Protection Area of the Comunidad de Madrid (PROEX 71.4/20 and PROEX 072.2/24) and are reported in accordance with the ARRIVE guidelines^52^.

Human coronary artery samples were obtained from patients undergoing heart transplantation at Hospital de la Santa Creu i Sant Pau (Barcelona, Spain) following approval by the hospital’s Ethics Committee (approval numbers: 19/267 and 23/111). Patients had end-stage dilated cardiomyopathy and underwent heart transplantation according to the recommendations of the European Society of Cardiology guidelines^26^. The use of discarded human tissue complied with the principles of the Declaration of Helsinki (1975, revised in 2013). Written informed consent was obtained from all participants or their legal representatives.

### Mice

Mice were maintained under specific pathogen-free conditions in individually ventilated cages under a 12-hour light/12-hour dark cycle at 22 ± 2 °C and 50% relative humidity (45-60%), with *ad libitum* access to water and standard chow diet (D184, SAFE). All mice were maintained on a C57BL/6J genetic background.

Experiments were performed using atheroprone *Ldlr*⁻^/^⁻ *Lmna^flox/flox^ Cdh5Cre^ERT2+/-^*mice (hereafter referred to as *Lmna*KO*^iEC^*), and their littermate controls (*Ldlr*⁻^/^⁻ *Lmna^flox/flox^ Cdh5Cre^ERT2^*⁻^/^⁻). The floxed *Lmna* allele (*Lmna^tm1Stw^*) contains loxP sites flanking exon 2^24^. The endothelial-specific *Cdh5Cre^ERT2^* transgene corresponds to *Tg(Cdh5-cre/ERT2)1Rha*^25^, and the *Ldlr* knockout allele (*Ldlr*^tm1Her^) was introduced from the JAX strain B6.129S7-*Ldlr*^tm1Her/J^ (stock #002207). Experimental and control animals were littermates.

At 4 weeks of age, mice received intraperitoneal tamoxifen (T5648, Sigma; 0.2 mg/g body weight) dissolved in corn oil (C8267, Sigma) once daily for 3 consecutive days to induce endothelial-specific *Lmna* recombination. Where indicated, mice were fed a HFD containing 10.7% fat and 0.75% cholesterol (S9167-E011, Ssniff) for 6 or 12 weeks.

Both male and female mice were included in all experiments. Unless otherwise indicated, mice were euthanized by CO_2_ inhalation. Independent cohorts of young and aged wild-type C57BL/6J mice (The Jackson Laboratory) maintained on standard chow were used to assess physiological aging. Mice were identified by numerical codes, and investigators were blinded to genotype during data acquisition and primary data analysis, except for experiments shown in Figures 3E-G, 4A and 7G. Genotypes were unblinded only after completion of the analyses.

### Quantification of lamin A/C expression in mouse aortic and cardiac cell populations by flow cytometry

Hearts and aortas were harvested from young and aged wild-type mice (ages are indicated in the figures and main text). Mice were euthanized and transcardially perfused with 10 mL of ice-cold phosphate-buffered saline (PBS) to remove circulating blood. Tissues were dissected and placed in ice-cold DMEM (Sigma) supplemented with 0.292 mg/mL L-glutamine (HyClone), 0.1 mM sodium pyruvate (SH30239.01, HyClone) and antibiotics (100 U/mL penicillin and 0.1 mg/mL streptomycin, Sigma). Aortas were cleared of perivascular adipose tissue, opened longitudinally, and digested in DMEM containing collagenase A (6.25 mg/mL; Roche), dispase II (6.25 mg/mL; Roche), DNase (62.5 μg/mL; Roche), and elastase (0.17 mg/mL; Sigma) at 37 °C for 15 min. Hearts were minced into ∼1 mm^3^ fragments, and digested under the same conditions, except that elastase was omitted. Cell suspensions were filtered through a 70-μm cell strainer, centrifuged (400 x g, 5 min, 4 °C), and resuspended in cold sorting buffer (PBS containing 1% FBS, 2 mM EDTA, 5 mM glucose, and 10 mM HEPES).

For immunostaining, Fc receptors were blocked with anti-CD16/CD32 (1:200, 101302), followed by staining with Zombie Yellow (1:100, 423103), anti-CD31-Alexa647 (1:100, 102416), and anti-CD45-Pacific Blue (1:100, 103126) (all from BioLegend) for 1 h on ice in the dark with gentle mixing after 30 minutes. Cells were washed in cold sorting buffer, fixed in 4% formalin (VWR), permeabilized with PBS containing 0.05% Triton X-100 (Sigma) for 5 min, washed in cold sorting buffer, and incubated with anti-lamin A/C Alexa488 antibody (1:50, 8671, Cell Signaling Technology) for 1 h on ice in the dark with gentle mixing after 30 minutes. Samples were washed in sorting buffer and analyzed on a BD FACSymphony flow cytometer using FACSDiva software (BD Biosciences). Unstained cells, Zombie Yellow-only controls, and single-color compensation controls prepared with UltraComp eBeads (Thermo Scientific) were included for gating and fluorescence compensation. Data were analyzed using FlowJo software (version 10.10.0, BD Biosciences). Gating strategy is shown in **Supplemental Figures** S**1 and S2**.

### Immunohistochemical analysis of human coronary arteries

Human coronary artery samples were collected from two groups of patients defined according to their age: young (range: 14-30 years [mean: 23.2 ± 2.4 years]; n=8) and old (range: 59-68 years [mean: 65.7 ± 0.98 years]; n=10). Coronary arteries from both groups showed atherosclerotic lesions ranging from early to intermediate stages. Patients with advanced, complicated, or inflamed coronary atherosclerotic lesions were excluded. Arterial specimens were fixed in 4% paraformaldehyde in 0.1 M PBS (pH 7.4), embedded in paraffin, and sectioned at 5 µm. Sections were deparaffinized, rehydrated through graded ethanol solutions, and subjected to heat-induced antigen retrieval using either 10 mM citrate buffer (pH 6.0) or 10 mM Tris/1 mM EDTA buffer (pH 9.0) in a water bath at 95-99 °C for 20 min. Endogenous peroxidase activity was blocked with 3% hydrogen peroxide in methanol for 30 min. Sections were permeabilized with 1% Tween-20 in PBS and blocked with 0.1% Tween-20/PBS containing 10% serum from the species in which the secondary antibody was raised.

Endogenous avidin and biotin were blocked using a commercial kit (Vector Laboratories). Sections were incubated overnight at 4 °C with primary antibodies against CD31 (ready to use, IR610, Agilent) or lamin A/C (1:100, sc-376248, Santa Cruz). After washing, sections were incubated with the appropriate biotinylated secondary antibody (Vector Laboratories) for 1 h at room temperature, followed by the VECTASTAIN ABC avidin-biotin peroxidase complex (Vector Laboratories) for 30 min. Immunoreactivity was visualized using 3,3′-diaminobenzidine (Roche). Sections were counterstained with hematoxylin, dehydrated, and mounted using DPX mounting medium (Casa Álvarez). Negative controls omitting the primary antibody were included to assess nonspecific staining. Lamin A/C-positive ECs were quantified in preserved arterial endothelium identified by CD31 staining in consecutive tissue sections, and results were expressed as lamin A/C-positive cells relative to the total number of CD31-positive ECs.

### Isolation of mouse liver ECs by fluorescence-activated cell sorting

Livers were harvested from control and *Lmna*KO*^iEC^*mice 12 weeks after tamoxifen administration. Mouse euthanasia, tissue collection, and digestion were performed as described above for flow cytometry, except that elastase was omitted from the digestion buffer. Cell suspensions were stained with propidium iodide (PI) (1 µM, Sigma), anti-CD31-Alexa647 (1:100, 102416, BioLegend), and anti-CD45-Pacific Blue (1:100, 103126, BioLegend). Viable ECs were isolated on a FACSAria II cell sorter (BD Biosciences). Cells were first gated based on forward and side scatter (FSC/SSC), followed by selection of viable ECs (PI⁻CD31⁺CD45⁻) and the non-EC fraction (PI⁻CD31⁻). Sorted cells were collected directly into 750 µL of QIAzol reagent (Qiagen) and stored at -80 °C until further processing.

### RNA isolation and RT-qPCR

Total RNA from sorted mouse ECs, mouse aortic arch samples, or HAECs was isolated using the RNeasy Mini Kit (Qiagen). Aortic arch samples were homogenized using a TissueLyser system (Qiagen). For sorted mouse EC, RNA extraction was performed according to the manufacturer’s instructions up to the phase separation step. The aqueous phase was then mixed 1:1 with 70% ethanol and applied to RNeasy Mini Kit columns (Qiagen) to complete the purification procedure. cDNA was synthesized using the High-Capacity cDNA Reverse Transcription Kit (Applied Biosystems). RT-qPCR was performed on a QuantStudio 5 Real-Time PCR System (384-well format, Applied Biosystems) using Power SYBR Green PCR Master Mix (Thermo Scientific). The thermal cycling protocol consisted of 50 °C for 2 min, followed by 95 °C for 10 min and 40 cycles of 95 °C for 15 s and 60 °C for 1 min. Primer sequences are listed in **Table S1**. Gene expression was normalized to *Gapdh* and relative expression was calculated using the 2^−(ΔΔCt)^ method. For HAEC experiments, results were expressed as fold change relative to the corresponding control condition in each independent experiment. All reactions were performed in technical triplicates.

### Histology and immunofluorescence on paraffin-embedded sections

Mouse tissues were fixed in 4% formalin (VWR), dehydrated through a graded ethanol series, embedded in paraffin, and sectioned at 4 μm using a microtome (RM2245, Leica). For immunofluorescence analysis, paraffin sections were deparaffinized, rehydrated, and rinsed in PBS. Antigen retrieval was performed by heating sections in 10 mM sodium citrate buffer (pH 6.0) for 20 min. Sections were then permeabilized and blocked for 1 h at room temperature in PBS containing 0.3% Triton X-100, 5% bovine serum albumin (BSA), and 5% normal goat serum (005-000-001; Jackson ImmunoResearch). Primary and secondary antibodies were diluted in PBS containing 0.3% Triton X-100, 5% BSA, and 2.5% normal goat serum.

For thoracic aorta (TA) immunofluorescence, sections were incubated overnight at 4 °C with anti-CD31 antibody (1:100, DIA-310, Dianova), followed by incubation for 2 h at room temperature with Hoechst 33342 (10 μM, B2261, Sigma), goat anti-rat Alexa Fluor 488 (1:400, 11006, Invitrogen), anti-smooth muscle α-actin-Cy3 (SMA-Cy3, 1:400, C6198, Sigma), and anti-lamin A/C Alexa Fluor 647 (1:100, sc-376248, Santa Cruz). Sections were mounted with Fluoromount-G (eBioscience) and imaged using a Zeiss LSM 700 confocal microscope.

### Whole-mount *en face* immunofluorescence of thoracic aorta

*En face* immunofluorescence of TA was performed as described^53^. In brief, mice were anesthetized by intraperitoneal injection of ketamine (225 mg/kg; Ketamidor, Richter Pharma) and xylazine (15 mg/kg; Nerfasin, Fatro). Animals were perfused transcardially through the left ventricle with 4% paraformaldehyde in PBS for 5 min. Aortas were excised, cleared of perivascular adipose tissue under a stereomicroscope (SZX3, Olympus Life Science), opened longitudinally along the outer curvature, pinned lumen-side up onto silicone-coated dishes, and fixed overnight in 4% paraformaldehyde at 4 °C. The following day, aortas were rinsed three times with PBS, cut into 4-mm segments, and processed as free-floating preparations. Tissues were blocked and permeabilized in PBS containing 0.3% Triton X-100, 5% BSA, and 5% normal goat serum (005-000-001; Jackson ImmunoResearch) (1 h at room temperature under gentle agitation). Primary and secondary antibodies were diluted in PBS containing 0.3% Triton X-100, 5% BSA, and 2.5% normal goat serum. Samples were incubated overnight at 4 °C with anti-ERG Alexa Fluor 488 (1:100, ab196374, Abcam) and anti-lamin A/C Alexa Fluor 647 (1:50, sc-376248, Santa Cruz), followed by goat anti-Armenian hamster IgG H&L Alexa Fluor 488 (1:400, ab173003, Abcam) for 2 h at room temperature under gentle agitation. Samples were mounted lumen side up in glycerol and imaged using a Zeiss LSM 700 confocal microscope. Images were quantified using ImageJ Fiji software^54^. Maximum-intensity projections of z-stacks were generated, and EC nuclei were identified by ERG immunostaining. Lamin A/C expression was quantified in ERG⁺ cells from comparable aortic regions using a fixed fluorescence intensity threshold applied uniformly to all samples. Results are expressed as the percentage of lamin A/C⁺ ECs relative to the total number of ERG⁺ ECs. Quantification was performed in three 25× fields per animal, and the mean value was used for statistical analysis.

### Longitudinal monitoring of survival and body weight

Mouse longitudinal survival studies began at 8 weeks of age, 4 weeks after tamoxifen administration, and body weight was monitored weekly. Diseased animals were examined and followed up by a specialized veterinarian blinded to genotype. Animals that met predefined humane endpoint criteria, in accordance with institutional ethical guidelines, were euthanized and their deaths recorded. Animals euthanized because of prolapse, malocclusion, or severe dermatitis were censored from the survival analysis.

### Analysis of atherosclerotic burden in mice

Hearts and aortas were harvested from mice after *in situ* perfusion with PBS through the left ventricle and fixed in 4% paraformaldehyde in PBS at 4 °C. For quantification of atherosclerotic burden, the aortic arch and TA were cleaned of surrounding adipose tissue and incubated in 78% methanol with gentle agitation for 1 h at room temperature, followed by staining with 0.2% Oil Red O (O0625, Sigma) as described^55^. The aorta was opened longitudinally along the inner curvature and pinned lumen-side up onto a silicone surface. Images were acquired using an Olympus UC30 digital camera mounted on a stereomicroscope (Olympus Life Science). Total aortic area and Oil Red O-positive plaque area were quantified by planimetric analysis using SigmaScan Pro 5 (Systat Software) and Fiji/ImageJ software^54^.

For histological analysis of atherosclerotic lesions in the aortic root, hearts were embedded in paraffin and sectioned at the level of the aortic valve leaflets. Serial 4-μm sections were stained with Masson’s trichrome and scanned using an Axio Scan Z1 (Zeiss). Images were exported using ZEN 3.4 blue software (Zeiss) and analyzed with Fiji/ImageJ^54^. Plaque area was manually delineated, and the mean plaque area was calculated from four sections separated by approximately 40 µm. To account for sex-dependent differences in atherosclerotic burden, plaque area measurements were normalized to the mean value of the corresponding sex-matched control group before pooling male and female data for statistical analysis.

### Wire myography analysis of vasoreactivity in mouse aortic rings *ex vivo*

Aortas were carefully dissected to remove surrounding fat and connective tissue, then immediately placed in cold Krebs-Henseleit solution (KHS: 115 mmol/L NaCl, 25 mmol/L NaHCO_3_, 4.7 mmol/L KCl, 1.2 mmol/L MgSO_4_·7H2O, 2.5 mmol/L CaCl_2_, 1.2 mmol/L KH_2_PO_4_, 11.1 mmol/L glucose, and 0.027 mmol/L EDTA). TAs were segmented in rings approximately 2 mm in length and mounted on a wire myograph system (Model 620M, DMT) for isometric tension measurements. Following a 30-minute equilibration period in KHS, continuously oxygenated with a gas mixture of 95% O₂ and 5% CO₂ at 37 °C (pH 7.4), diameter-tension relationships were assessed by progressively increasing the distance between the two wires inserted through the lumen, thereby increasing the vessel passive diameter. At each step, both the developed force and the internal circumference were recorded. Subsequently, segments were adjusted to their optimal lumen diameter for active tension generation. For this purpose and subsequent analysis, LabChart software (ADInstruments) was used. This optimal point was determined by calculating the ratio of internal circumference to wall tension, setting the internal circumference (L_0_) to 90% of the value corresponding to a passive tension equivalent to that produced by a transmural pressure of 100 mm Hg^56^.

To assess vascular contractility, segments were first exposed to a depolarizing high-potassium solution (K⁺-KHS, 120 mmol/L). After equilibration, aortic rings were precontracted with phenylephrine to achieve approximately 50% of the maximal contraction induced by K⁺-KHS, after which endothelial function was assessed by recording a vasodilation response curve to cumulative doses of acetylcholine (from 1 nmol/L to 10 µmol/L). Following washing and re-equilibration, response curves to cumulative doses of phenylephrine (from 1 nmol/L to 30 µmol/L) were recorded. Finally, endothelium-independent relaxation was evaluated by generating concentration-response curves to the nitric oxide donor diethylamine NONOate (DEA-NO; from 1 nmol/L to 10 µmol/L) in phenylephrine-precontracted segments.

### MRI-based assessment of endothelial function and permeability *in vivo*

MRI studies were performed on a 9.4 T BioSpec 94/20 USR scanner (Bruker). Mice were anesthetized with isoflurane (Aerrane, 1.7%) in an oxygen/air mixture (1:2) and positioned supine for imaging. Cardiac rhythm, electrocardiogram (ECG), respiration, and body temperature (maintained at 37 °C) were continuously monitored using a monitoring and gating system (SA Instruments).

Endothelial function was evaluated in both the TA and abdominal aorta following previously described procedures with minor modifications^57,58^. Endothelium-dependent and -independent vasodilatory responses were induced by intravenous administration of cevimeline (6 mg/kg) and sodium nitroprusside (1 mg/kg), respectively, via tail-vein cannulation. Two time-resolved 3D MRI datasets of the aorta were acquired, one before and one 30 minutes after drug administration, to assess vasomotor responses. Imaging was performed using a cine IntraGate FLASH 3D sequence, and datasets were reconstructed with the IntraGate 1.2.b.2 macro (Bruker). Image analysis was performed with ImageJ and MATLAB (MathWorks) scripts. For both the TA (10 diastolic slices from the celiac artery upward) and the abdominal aorta (10 diastolic slices from the renal arteries downward), vessel cross-sectional areas were segmented by thresholding and used to reconstruct and quantify total vessel volumes. Vasomotor responses were expressed as the percentage change in vessel volume relative to pre-administration measurements, with positive and negative values indicating vasodilation and vasoconstriction, respectively. Imaging parameters for the endothelial function study were: repetition time 6.4 ms; echo time 1.4 ms; field of view 30 × 30 × 14 mm³; matrix size 256 × 256 × 35; flip angle 30°; number of averages 15; reconstructed into 7 cardiac frames. Total scan duration was approximately 12 min.

Endothelial permeability in the brachiocephalic artery was assessed using the same cine IntraGate FLASH 3D sequence combined with a variable flip angle approach to obtain T₁ relaxation maps. T₁ maps acquired before and immediately after intravenous injection of gadolinium-rich liposome contrast agent (Gadobutrol in the liposome, formulation concentration, 287 mg/mL; dose, 4.5 mL/kg) were compared pixel by pixel using MATLAB scripts^58–60^. The number of pixels exhibiting a T₁ reduction greater than 50% after contrast administration (Npx50) was calculated as an index of increased vascular permeability. The threshold value (50%) was determined experimentally^57^. Acquisition parameters for the permeability protocol were: repetition time 10 ms, echo time 1.1 ms, field of view 30 × 30 × 4 mm³, matrix size 192 × 160 × 8; 12 repetitions; reconstructed into a single cardiac frame. Eight flip angles (2°, 4°, 6°, 8°, 14°, 20°, 30°, 50°) were applied by modulating radiofrequency pulse length at constant amplifier power. Total acquisition time was approximately 16 min.

### Assessment of aortic pulse wave velocity

Aortic PWV was evaluated using a Doppler flow velocity system (Indus Instruments) as previously described^61^. Mice were anesthetized with 1.5% isoflurane (Aerrane) delivered in an oxygen/air mixture (1:2). During measurements, animals were positioned supine on ECG electrodes integrated into a temperature-controlled platform. Core temperature was continuously monitored via a rectal probe and maintained at 37 °C. Prior to data acquisition, the thoracic area and right hind limb were shaved to optimize signal quality.

PWV was determined using two 20-MHz Doppler probes to simultaneously record flow velocity waveforms from two distinct aortic segments^62^. For TA measurements, one probe was placed over the aortic arch, while the second was positioned approximately 15-20 mm downstream along the descending aorta. To capture the descending aortic signal, the probe was placed just below the sternum and angled toward the heart at a depth of approximately 4 mm. The second probe for TA recordings was positioned at the base of the left forelimb, aligned nearly parallel to the platform and directed toward the upper mid-chest at a depth of approximately 6 mm. For abdominal aorta measurements, flow velocities were recorded from the descending aorta and the right femoral artery (≈25-30 mm apart). The second probe was positioned on the right hind limb and angled toward the groin region at a depth of 3-4 mm. PWV was calculated as the distance between the two recording sites divided by the difference in the arrival time of the flow velocity waveforms, each referenced to the ECG signal. Data acquisition and analysis were performed using the Doppler signal processing workstation (version 1.625, Indus Instruments).

### Systolic blood pressure measurements

SBP was measured in conscious mice using a BP2000 non-invasive tail-cuff plethysmography system (Visitech Systems). Animals were acclimatized to the procedure for one week prior to data collection, undergoing daily training sessions to minimize stress-induced variability. All measurements were performed at the same time of day in a quiet, temperature-controlled environment. During recordings, mice were placed in restrainers on a warming platform to ensure adequate tail blood flow. For each animal, at least seven stable readings were obtained and averaged to determine the final SBP value, excluding any irregular traces due to movement artifacts.

### Transthoracic echocardiography studies

Transthoracic echocardiography was performed by a blinded expert operator at CNIC’s Advanced Imaging Unit using a high-frequency ultrasound imaging system (Vevo 2100, Vevo 3100, and F2; VisualSonics Inc.) with a 30-MHz linear probe. A base-apex ECG was continuously monitored via four leads connected to the ultrasound system. Isoflurane anesthesia was carefully adjusted to maintain a stable heart rate of 450 ± 50 beats per minute. Body temperature was maintained within the physiological range by placing the mice on a heated platform.

Images were acquired and subsequently analyzed offline in a blinded manner using Vevo Workstation software (VisualSonics). Standard parasternal long-axis two-dimensional (2D) and M-mode images were acquired to assess left ventricular (LV) structure and function. From M-mode recordings, left ventricular internal diameters in diastole and systole (LVIDd and LVIDs), interventricular septal thickness in diastole (IVSd), and posterior wall thickness in diastole (LVPWd) were measured. Left ventricular mass (corrected) was calculated using the geometric formula provided by the Vevo software. Two-dimensional measurements of LV area in systole and diastole (LVESA and LVEDA) were used to derive corresponding LV volumes (LVESV and LVEDV), and to calculate LV ejection fraction (LVEF). Stroke volume (SV), fractional shortening (FS), and cardiac output (CO) were automatically computed from M-mode data.

Diastolic function was evaluated from the 2D apical view using:

a. Pulsed-wave (PW) Doppler, to assess mitral inflow patterns and isovolumetric relaxation time (IVRT). Early (E) and late (A) diastolic peak velocities were measured, and the E/A and E/IVRT ratios were calculated. Pulmonary venous flow profiles were also acquired to determine systolic (PVs) and diastolic (PVd) velocities, and the PVs/PVd ratio.
b. Tissue Doppler imaging (TDI), to measure early (E′) and late (A′) mitral annular velocities and calculate the E′/A′ and E/E′ ratios.
c. Left atrial diameter (LA), measured in the parasternal long-axis view as the anteroposterior diameter (mm).

### Cardiac cine MRI acquisition

Animals were anesthetized with isoflurane (2% in oxygen, 0.5 L/min) and monitored for core body temperature and respiration rate using a dedicated monitoring system (SA Instruments). *In vivo* cardiac images were acquired using a 7 T MRI scanner (Agilent Varian), operated via an Avance NEO Spectrometer (Bruker). An IntraGate FLASH sequence was used in combination with a 112/86 mm resonator and a surface heart array coil (Bruker). Cardiac four-chamber and left two-chamber views were obtained to plan the short-axis image acquisition. Imaging parameters were as follows: number of slices, 11-14; slice thickness, 0.8 mm; matrix size, 256 × 256; field of view, 25 × 25 mm²; frames per cardiac cycle, 12; oversampling, 100%; echo time (TE), 1.58 ms; repetition time (TR), 8.5 ms; and flip angle, 20°. Cine magnetic resonance images were exported in DICOM format and analyzed using Segment software (Medviso AB, version 4.1.0.1 R14284b). The short-axis image stack was analyzed by manual delineation of the left and right ventricular (LV and RV) endocardial borders at end-diastole and end-systole. Papillary muscles and trabeculae were included in the segmentation. From these analyses, left and right ventricular end-diastolic volumes (LVEDV, RVEDV), end-systolic volumes (LVESV, RVESV), and ventricular masses (LVM, RVM) were obtained. The LVM/LVEDV and RVM/RVEDV ratios were calculated as indices of concentricity.

### Cardiac histological analysis of fibrosis

Hearts were dissected after *in situ* perfusion with phosphate-buffered saline (PBS) injected through the left ventricle. Samples were fixed and embedded in paraffin as previously described. Serial 4-µm sections were obtained using a microtome (Leica) and stained with Masson’s trichrome to visualize collagen fibers. Stained sections were scanned using an Axio Scan Z1 slide scanner (Zeiss), and images were exported with ZEN 3.4 blue software (Zeiss). Quantification of collagen content was performed using ImageJ^54^ with user-customized macros. To isolate the collagen signal, color deconvolution was applied, and the proportion of the green-stained area (representing collagen) was quantified across the entire heart section. Results were expressed as the percentage of total tissue area occupied by collagen.

### Assessment of heart weight

Mice were euthanized by CO₂ inhalation and perfused through the left ventricle with PBS to remove circulating blood. The heart was then excised, blotted dry, and weighed. Heart weight was used as an index of cardiac hypertrophy and normalized to tibia length.

### Serum lipid profile and NT-proBNP quantification

Blood samples were collected from the right ventricle immediately after euthanasia and allowed to clot at room temperature for 2-3 h before serum isolation. Samples were then centrifuged at 2000 × g for 10 min at 4 °C. The resulting supernatant was carefully transferred to a new microcentrifuge tube and centrifuged again under the same conditions to ensure complete removal of cellular debris. The clarified supernatant was collected, aliquoted, and stored at −80 °C until further analysis. Hemolyzed samples were excluded from further analysis. Lipid profiles, including total cholesterol, free cholesterol, LDL-C, HDL-C, triglycerides, and non-esterified fatty acids were measured using a Dimension RxL Max Integrated Chemistry System (Siemens Healthineers) by specialized staff at the CNIC Animal Facility. Serum NT-proBNP concentrations were measured by ELISA (MBS2501591, MyBioSource) according to the manufacturer’s instructions and analyzed using an xMark™ Microplate Spectrophotometer (Bio-Rad).

## High-throughput cardiac proteomic analysis

### Preparation of protein extracts and protein digestion

Hearts from 9-month-old control and *Lmna*KO*^iEC^*mice fed HFD for 12 weeks were dissected after transcardial perfusion with 10 mL of ice-cold PBS and snap-frozen in liquid nitrogen. Samples were pulverized and boiled for 10 min in lysis buffer (100 mM Tris-HCl pH 6.8, 2% sodium dodecyl sulfate, 10 mM dithiothreitol). Protein extracts were quantified using the RC DC Protein Assay (Bio-Rad) and loaded onto FASP filters (Expedeon) for overnight digestion at 37 °C with sequencing grade trypsin (Promega) at a 1:40 (w/w) trypsin-to-protein ratio, following the manufacturer’s instructions. The resulting tryptic peptides were recovered by centrifugation, after which trifluoroacetic acid was added to a final concentration of 1%. Peptides were desalted on C18 Oasis HLB extraction cartridges (Waters Corporation), and vacuum-dried.

### Isobaric labeling of tryptic peptides

The peptide samples were resuspended in 1 M triethylammonium bicarbonate and quantified using a Direct Detect IR spectrometer (Millipore). Equal amounts of peptides were isobarically labeled with tandem mass tags (TMT) 16-plex reagents (Thermo Fisher Scientific) according to the manufacturer’s instructions. Labeled peptide samples were pooled and dried under vacuum.

### Peptide fractionation

Aliquots of the dried labeled peptide samples were resuspended in 300 µL of 0.1% trifluoroacetic acid and separated into five fractions using the High pH Reversed-Phase Peptide Fractionation Kit (Thermo Fisher Scientific). The spin column was equilibrated with 300 µL of acetonitrile twice, followed by two washes with 300 µL of 0.1% trifluoroacetic acid. Samples were loaded onto the column and centrifuged at 3,000 x g for 2 min, followed by a wash with 300 µL of water. Bound peptides were eluted into five fractions with 300 µL of freshly prepared solutions containing: (1) 12.5% acetonitrile, 87.5% triethylamine; (2) 15% acetonitrile/85% triethylamine; (3) 17.5% acetonitrile/82.5% triethylamine; (4) 20% acetonitrile/80% triethylamine; and (5) 50% acetonitrile/50% triethylamine. Fractions were vacuum-dried and stored at −20 °C until nano-flow liquid chromatography coupled to tandem mass spectrometry (LC-MS/MS) analysis.

### Liquid chromatography tandem mass spectrometry analysis

The labeled peptide samples were resuspended in 0.1% formic acid and analyzed by LC-MS/MS using an Easy-nLC 1000 HPLC system (Thermo Fisher Scientific) coupled online to a Fusion Tribrid mass spectrometer (Thermo Fisher Scientific). C18-based reversed phase separation was performed using a 2-cm trap column (Acclaim PepMap100, 75-µm internal diameter, 3-µm particle size; Thermo Fisher Scientific) and a 50-cm analytical column (EASY-Spray PepMap RSLC C18, 75-µm internal diameter, 3-µm particle size, Thermo Fisher Scientific). Peptides were loaded in buffer A, 0.1% formic acid (v/v) and eluted with a linear gradient (260-360 min) of buffer B, 90% acetonitrile/0.1% formic acid (v/v) at 200 nL/min flow. MS spectra were acquired in the Orbitrap analyzer over an m/z range of 390-1500 at a resolution of 70,000. Higher-energy collisional dissociation MS/MS spectra were acquired for the top 20 ions in each full MS scan using data-dependent acquisition, 30 normalized collision energy, 45 s dynamic exclusion, and Orbitrap resolution of 30,000.

### Peptide and protein identification

The SEQUEST HT algorithm integrated in Proteome Discoverer 2.5 (Thermo Fisher Scientific) was used for peptide identification from LC-MS/MS data. The MS/MS spectra were searched against a concatenated target-decoy database containing mouse protein sequences (UniProtKB/Swiss-Prot 2024_02 Release). Database search parameters included trypsin digestion with a maximum of two missed cleavage sites, precursor and fragment mass tolerance of 800 ppm and 0.02 Da, respectively, fixed TMT modification at peptide N-termini and lysine residues, fixed carbamidomethylation of cysteine, and variable oxidation of methionine. Peptide identification was performed using the probability ratio method^63^. False discovery rate was calculated using the refined method after 15-ppm precursor mass tolerance post-filtering, and a 1% false discovery rate threshold was applied for peptide identification^64,65^. Peptides were assigned to the most probable protein using an in-house developed algorithm available at ProteinAssigner _v4.py.

### Quantification at the peptide and protein levels

Peptide and protein abundance were quantified using iSanXoT, a publicly available implementation of the WSPP statistical model and the Generic Integration Algorithm^66–68^. Protein log2-ratios were expressed in units of standard deviation according to their estimated variance (Zq values). Quantified proteins were functionally annotated using the DAVID repository, including Gene Ontology, KEGG, and Panther databases, among others^69^. The Limma package was used to ascertain statistical significance by means of p-values^70^.

### Protein-protein interaction network analysis (STRING)

Protein-protein interaction network analysis was performed using STRING (version 12.0). Proteins significantly upregulated in *Lmna*KO*^iEC^*hearts compared with controls were analyzed using *Mus musculus* as the reference organism. Interaction networks were generated using a minimum interaction score of 0.4 (medium confidence). Markov clustering was performed with an inflation parameter of 2.5 to identify clusters of functionally associated proteins.

### Gene set enrichment analysis

To evaluate the similarity between the cardiac proteomic signature of *Lmna*KO*^iEC^* mice and human HFpEF, Gene Set Enrichment Analysis (GSEA) was performed using the GSEA Desktop application (Broad Institute, version 4.4.0). Using a published human HFpEF proteomic dataset^28^, a custom protein set was generated by selecting proteins significantly upregulated in hearts from HFpEF donors compared with controls (n=10, log_2_ fold change > 0, adjusted *P* < 0.05). This protein set was tested for enrichment against our preranked cardiac proteomic dataset, ranked according to the formula -log_10_P × sign (average Zq *Lmna*KO*^iEC^*– average Zq controls). Preranked GSEA was performed using the default parameters (1,000 permutations, weighted enrichment statistics, meandiv normalization mode, and permutation seed 149). Enrichment significance was assessed using the normalized enrichment score and false discovery rate.

## Single-cell RNA sequencing analysis of atherosclerotic mouse aorta

### Aorta collection and single-cell suspension preparation

Aortas were collected from 9-month-old control and *Lmna*KO*^iEC^* mice fed HFD for 12 weeks. Mice were euthanized and perfused transcardially through the LV with 10 mL of cold PBS to remove blood from the aorta. The aortic arch and TA were dissected, cleared of perivascular adipose tissue on a Petri dish kept on ice, and opened longitudinally in cold DMEM (Sigma) supplemented with penicillin (100 U/mL), streptomycin (0.1 mg/mL; Sigma), L-glutamine (0.292 mg/mL; HyClone), and sodium pyruvate (0.1 mM; HyClone). Each biological replicate per genotype consisted of a pool of four aortas (two males and two females to minimize potential sex-related bias). Two biological replicates were included for each genotype.

Viable single-cell suspensions were obtained as previously described^29^. Briefly, aortas were digested at 37 °C for 15 min in DMEM containing collagenase A (6.25 mg/mL; Roche), dispase II (6.25 mg/mL; Roche), DNase I (62.5 µg/mL; Roche), and elastase (0.17 mg/mL; Sigma), followed by an additional digestion step at 37 °C for 5 min in 0.25% trypsin-EDTA (Gibco). Following digestion, cells were pelleted by centrifugation and resuspended in sorting buffer. Hoechst 33342 (10 µg/mL; Sigma) was included in both digestion and sorting buffers to label cell nuclei. Cell suspensions were passed through a 70-µm cell strainer, stained with TO-PRO-3 (1 µM; Invitrogen), and purified using a FACSAria cell sorter (BD Biosciences). Cells were gated by forward and side scatter to remove debris, and only viable nucleated cells (Hoechst 33342⁺TO-PRO-3⁻) were collected. Sorted cells were pelleted and resuspended in sorting buffer at an approximate concentration of 1,400 cells/µL for downstream single-cell RNA sequencing.

### Sequencing

Cells were counted and their viability was assessed using a Countess III cell counter (Thermo Fisher Scientific). Sample viability ranged from 71% to 81%. Each cell suspension was loaded into one channel of a Chromium Next GEM Chip G (10x Genomics). Single cells were encapsulated into emulsion droplets using the Chromium Controller (10x Genomics). Sequencing libraries were prepared using the Chromium Next GEM Single Cell 3’ Kit v3.1 (10x Genomics) according to the manufacturer’s instructions. Library size was then calculated using a High Sensitivity DNA chip on a 2100 Bioanalyzer (Agilent Technologies), and concentration was determined using a Qubit fluorometer (Thermo Fisher Scientific). Individual libraries were diluted to 4 nM and pooled for sequencing. The library pool was loaded at 700 pM onto P3 and P2 flow cells (100 cycles) of a NextSeq 2000 instrument (Illumina) in paired-end configuration (28bp Read1, 10bp Index 1, 10 bp Index 2, and 90 bp Read 2). FASTQ files for each sample were generated using the Cell Ranger mkfastq pipeline (10x Genomics).

### Alignment and preprocessing

Raw sequencing data were processed using the Cell Ranger pipeline (v6.1.1; 10x Genomics)^71^. Reads were aligned to the mm10 mouse reference genome (refdata-gex-mm10-2020-A) to generate gene-cell count matrices for downstream analysis.

### Quality control and filtering

Initial quality control and filtering were performed using the Scater (v1.26.1) and Seurat (v4.3.0) R packages^71,72^. Single cells were retained based on the following criteria:

- Sequencing depth between 4,000 and 80,000 reads
- Detection of at least 2,500 genes per cell
- Mitochondrial content < 20%
- Gene expression complexity < 65% (percentage of reads from the top 50 genes)
- Hemoglobin gene set expression < 2%

Cells not meeting these thresholds were excluded, and mitochondrial and hemoglobin genes were removed from downstream analyses. After filtering, 16,936 cells were retained. Doublets were detected and excluded using the scDblFinder (v1.12.0) algorithm^73^, resulting in a final dataset of 15,756 cells.

### Normalization, dimensionality reduction, and clustering

Gene counts were normalized, and the 2,000 most variable genes were identified using variance-stabilizing transformation. Dimensionality reduction was performed using principal component analysis, retaining the first 20 principal components, excluding those strongly correlated with cell cycle genes. Cells were clustered using Seurat’s implementation of the Louvain algorithm on a nearest-neighbor graph (resolution = 0.1). Cluster markers and differential gene expression between conditions were determined using the Wilcoxon rank-sum test, testing only genes expressed in >30% of cells (for marker identification) or >10% (for differential expression).

### Functional annotation and subclustering

Cell type annotation was performed manually based on the expression of established marker genes characteristic of each population. After identification of the main cell types, subclustering analyses were conducted separately for populations of ECs, VSMCs, hematopoietic cells, and fibroblasts, following the same analytical workflow described above. For each subset, highly variable genes were re-selected, principal component analysis was recomputed, and clustering parameters were optimized to best resolve subpopulation structure. Shared and unique differentially expressed genes among subclusters were visualized using UpSet plots to identify overlapping and subcluster-specific transcriptional signatures.

### Trajectory analysis

Trajectory inference was performed using the Slingshot R package (v2.8.0)^74^. For each analysis, UMAP embeddings obtained from the final Seurat subclustering were used as input. Separate trajectory analyses were conducted for macrophages and for the combined vascular and endothelial cell populations. Lineages were inferred based on the clustering structure, and pseudotime values were calculated for each cell along the identified trajectories. The resulting trajectories were projected onto the UMAP embeddings to visualize the inferred differentiation or transition paths across subpopulations.

### Functional enrichment and upstream regulator analysis

To identify predominant biological processes associated with each cluster, Gene Ontology (GO) enrichment analysis was performed using significantly enriched genes with DAVID bioinformatics resources^69^. Significance of enrichment was evaluated using the Benjamini-Hochberg adjusted p-value, and GO terms with adjusted p < 0.05 were considered significantly enriched. The most significantly enriched GO categories were used to aid in biological interpretation and cluster annotation. Analysis of predicted activation or inhibition of canonical pathways and upstream regulators was performed using IPA (Qiagen). Significance was determined using the Benjamini-Hochberg adjusted p-value, with pathways and regulators considered significant at adjusted p < 0.05.

### Calculation of a KLF2/KLF4 activation score

To visualize KLF2/KLF4 transcriptional activity in ECs, gene signature scoring was performed based on the expression of known downstream target genes, namely *Nos3*, *Thbd*, *Entpd1*, *Plpp3*, *Ucp2*, *Bmper*, and *Bmpr2*^30^. This score was obtained by aggregating the expression levels of the genes in the set using the AddModuleScore function from the Seurat package. Subsequently, differences in the score between *Lmna*KO*^iEC^* and control cells were analyzed using the Wilcoxon Rank Sum test.

### Cell culture

Immortalized human aortic endothelial cells (HAECs) were obtained from ATCC (CRL-4052) and cultured on 0.1% gelatin (Sigma) in Endothelial Cell Growth Basal Medium-2 (EBM-2, 00190860, Lonza) supplemented with the EGM-2 SingleQuots kit (CC-4176, Lonza). Cells were used between passages 2 and 6. Cultures were maintained at 37 °C in a humidified atmosphere containing 5% CO_2_.

### siRNA transfection and laminar shear stress stimulation in HAECs

HAECs were seeded on 60-mm dishes. When cells reached 70-80% confluence (16-24 h after seeding), they were transfected with 30 nM *LMNA* siRNA SMARTpool (Dharmacon). Control cells were transfected with an equal concentration of non-targeting siRNA (CT siRNA). All transfections were performed in Opti-MEM medium (Gibco) using Oligofectamine transfection reagent (Life Technologies). After 12 h, EGM-2 supplemented with 2x FBS and growth factors was added, and cells were cultured for an additional 36 h.

The *LMNA* siRNA SMARTpool (Dharmacon) included siRNAs targeting the following mRNA sequences:

- J-004978-05: GAAGGAGGGUGACCUGAUA
- J-004978-06: UCACAGCACGCACGCACUA
- J-004978-07: UGAAAGCGCGCAAUACCAA
- J-004978-08: CGUGUGCGCUCGCUGGAAA

The non-targeting control pool (CT siRNA) contained siRNAs targeting the following sequences:

- UGGUUUACAUGUCGACUAA
- UGGUUUACAUGUUGUGUGA
- UGGUUUACAUGUUUUCUGA
- UGGUUUACAUGUUUUCCUA

Following transfection, both control and *LMNA* siRNA-transfected HAECs were subjected to laminar shear stress using a custom-made cone-plate viscometer system, as previously described^75^. Shear stress was applied at a rate of 3 dyn/cm² for 28 h. Parallel cultures of control and *LMNA*-silenced HAECs were maintained under static conditions.

### Protein extraction and Western blot analysis

HAECs were lysed in radioimmunoprecipitation assay buffer (RIPA: 50 mM Tris-HCl pH 7.5, 100 mM NaCl, 1% Triton X-100, 0.1% SDS, 2 mM EDTA, and 0.5% sodium deoxycholate) supplemented with Complete protease inhibitor cocktail (Roche) and PhosSTOP Easypack phosphatase inhibitor (Roche). Samples were incubated on ice for 30 min with frequent vortexing and then sonicated at 4 °C for 5 min using a Bioruptor UCD-200 (Diagenode). Lysates were cleared by centrifugation at 10,000 x g for 30 min, and protein concentration was determined using the Pierce BCA Protein Assay Kit (Thermo Fisher Scientific).

Protein lysates (15 µg) were mixed with loading buffer containing 11 mg/mL 2-mercaptoethanol and incubated at 95 °C for 5 min. Proteins were separated on 7.5% polyacrylamide gels and transferred to PVDF membranes for 110 min at 4 °C and 300 mA. Membranes were blocked for 1 h at room temperature with 5% skimmed milk in TBST (20 mM Tris-HCl pH 7.4, 150 mM NaCl, 0.05% Tween 20), incubated overnight at 4 °C with primary antibodies against Lamin A/C (1:1000, sc-376248, Santa Cruz) and GAPDH (1:15000, MAB374, Sigma), and then incubated for 1 h at room temperature with m-IgGκ BP-HRP (1:5000, sc-516102, Santa Cruz). After three washes with TBST, membranes were incubated with Immobilon Classico Western HRP substrate (Merck Millipore). Protein bands were visualized using an iBright CL 1500 Imaging System (Thermo Fisher Scientific) and quantified with Fiji/ImageJ^54^. Protein abundance was normalized to GAPDH. Protein size was determined using Precision Plus Protein Dual Color Standards (Bio-Rad).

### Quantification of NO production by mouse thoracic aorta *ex vivo*

For *ex vivo* assessment of NOS3-dependent NO production, electron paramagnetic resonance spin- trapping with diethyldithiocarbamic acid sodium salt (DETC) was performed as previously described, with slight modifications^76^. The TA was isolated, cleared of surrounding adipose and connective tissue, opened longitudinally, and preincubated for 30 min at 37 °C in Krebs-HEPES buffer (99 mmol/L NaCl, 25 mmol/L NaHCO₃, 4.7 mmol/L KCl, 1.2 mmol/L MgSO₄·7H₂O, 2.5 mmol/L CaCl₂·H₂O, 1 mmol/L KH₂PO₄, 5.6 mmol/L glucose, and 20 mmol/L Na-HEPES) containing 10 μmol/L N^6^-(1-Iminoethyl)- lysine hydrochloride. This step enabled selective detection of NOS3-derived NO by inhibiting inducible NO synthase 2 activity.

For spin-trap preparation, DETC (3.6 mg) and FeSO₄·7H₂O (2.25 mg) were each dissolved under continuous argon bubbling in separate 10-mL aliquots of ice-cold Krebs-HEPES buffer and maintained on ice under argon flow until use. Following preincubation, spin-trapping colloid (125 μL FeSO₄·7H₂O + 125 μL DETC; final concentration, 285 μmol/L) and calcium ionophore A23187 (1 μmol/L) were added to the aortic preparations and incubated for 90 min at 37 °C. After incubation, the aortas were briefly blotted dry, suspended in fresh Krebs-HEPES buffer in a 400-μL column, rapidly frozen in liquid nitrogen, and stored at −80 °C until analysis. Electron paramagnetic resonance spectra were recorded using an X-band spectrometer (EMX Plus) equipped with a rectangular resonator cavity (H102). The NO-Fe(DETC)₂ signal was quantified by measuring the total peak amplitude after baseline correction. NO production was expressed as arbitrary units and normalized to the total protein content of each sample.

### Statistical analysis

Data are presented as mean ± standard error of the mean unless otherwise indicated. Statistical analyses were performed using GraphPad Prism 10 (GraphPad Software). Data normality was assessed using the Shapiro-Wilk test. For comparisons between two groups with a single variable, an unpaired two-tailed Student’s t-test was applied to normally distributed data, while the Mann-Whitney U test was used for non-parametric data. For experiments involving two independent variables, differences were evaluated using two-way ANOVA followed by Tukey’s post hoc test for multiple comparisons, unless stated otherwise. A p-value < 0.05 was considered statistically significant. The Wilcoxon rank-sum test was used for differential gene expression analysis in scRNA-seq datasets. Outliers identified with the ROUT test (Q = 1%) were excluded from further analysis.

## Data availability

Detailed information on the materials and methods used in this study is provided in the main article and the Supplemental Material. The data supporting the findings of this study are available within the article and its Supplemental Material or from the corresponding authors upon reasonable request. The mass spectrometry proteomics data have been deposited in the ProteomeXchange Consortium via the PRIDE partner repository under dataset identifier PXD081774. scRNA-seq data have been deposited in the BioStudies/ArrayExpress repository under accession number E-MTAB-17547, and the analysis code is available at https://github.com/LAB-VA-CNIC/17547.

## Acknowledgments

We thank Cristina González for assistance with mouse colony management and the CNIC Flow Cytometry Unit, Histology Service, Advanced Imaging Unit, Microscopy and Dynamic Imaging Unit (especially Verónica Labrador), Genomics Unit (especially Alberto Benguría), Proteomics Unit (especially Emilio Camafeita), and Animal Facility (especially Eva Santos) for their excellent technical support. Illustrations were created using BioRender.com (licensed to V.A.).

## Author contributions

**Conceptualization:** I.R.P., I.B., S.C., and V.A. **Methodology:** I.R.P., A. Bar, P.G., A.d.M.-M., M.G.-A., S.P., C.T., J.P., C.R., J.M.-G., J.M., S.C., I.B. and V.A. **Data generation and analysis:** I.R.P., A. Bar, P.G., A.d.M.-M., M.G.-A., A. Barettino, S.P., C.T., J.A., M.J.A.-M., J.P., G.d.R.H., Z.B., M.T., C.R. and J.M.-G. **Writing–review andediting:** I.R.P., B.D., I.B., S.H and V.A. **Supervision:** C.T., B.D., C.R., J.M.-G., J.M., S.C., I.B. and V.A.

## Non-standard Abbreviations and Acronyms

CVD: cardiovascular disease
DEA-NO: diethylamine NONOate
DETC: diethyldithiocarbamic acid sodium salt
EC: endothelial cell
ECM: extracellular matrix
GO: Gene Ontology
HAEC: human aortic endothelial cell
HFpEF: heart failure with preserved ejection fraction
HFD: high-fat diet
LC-MS/MS: nano-flow liquid chromatography coupled to tandem mass spectrometry
LINC: linker of nucleoskeleton and cytoskeleton
LSS: laminar shear stress
LV: left ventricle
MRI: magnetic resonance imaging
NO: nitric oxide
NOS3: endothelial nitric oxide synthase
PBS: phosphate-buffered saline
PI: propidium iodide
PWV: pulse wave velocity
RT-qPCR: reverse transcription quantitative polymerase chain reaction
SBP: systolic blood pressure
scRNA-seq: single-cell RNA sequencing
TA: thoracic aorta
UMAP: uniform manifold approximation and projection
VSMC: vascular smooth muscle cell

