## Supplemental Material for "Age-dependent downregulation of endothelial lamin A/C impairs mechanotransduction and promotes cardiovascular disease"

**INDEX**

Supplemental Results
Supplemental Discussion
Legends for Supplemental Tables
Supplemental Figures
Supplemental References

### Supplemental Results

#### Profibrotic fibroblast remodeling in *Lmna*KO$^{iEC}$ aortas

Subclustering of *Pdgfra*-enriched aortic fibroblasts (clusters C0, C5, and C7 in **Figure 6B**) identified seven transcriptionally distinct populations (FB0-FB6), present in both control and *Lmna*KO$^{iEC}$ mice (**Figure S11A-B**). To further characterize these subclusters, we identified specifically enriched genes and assessed differences in their relative abundance between genotypes (**Figure S11C-D and Table S3)**.

FB0 was enriched for genes associated with ECM remodeling, including *Fmod*, *Fbln7*, *Thbs2*, and *Col12a1*, and was less abundant in *Lmna*KO$^{iEC}$ aortas (**Figure S11C-D and Table S3)**. GO analysis confirmed enrichment for processes related to ECM and collagen fibril organization, suggesting that FB0 represents a classical fibroblast population involved in ECM remodeling (**Figure S12 and Table S4**).

FB1 was enriched in signaling-related genes, including *Flt1*, *Vegfb*, *Ntrk3*, *Ntf3*, and *Pdgfrb*, which are associated with vascular interactions and pro-survival pathways (**Figure S11C and Table S3)**. GO analysis revealed enrichment for MAPK signaling-related processes and response to hypoxia, consistent with a stress-responsive fibroblast population potentially involved in vascular remodeling (**Figure S12 and Table S4**).

FB2 was characterized by the expression of homeostatic ECM regulators (*Sparcl1*, *Apod*) together with *Meox2,* a transcription factor implicated in fibroblast quiescence, and was also reduced in *Lmna*KO$^{iEC}$ aortas (**Figure S11C-D and Table S3)**. These features suggest that FB2 represents a supportive fibroblast population contributing to ECM stability and vascular homeostasis.

FB3 was enriched in stemness- and migration-associated genes, including *Ly6a*, *Ly6c1*, *Igfbp6*, and *Cd248*, consistent with a population of activated, migratory fibroblasts with mesenchymal progenitor-like characteristics (**Figure S11C and Table S3)**.

FB4 displayed a robust inflammatory signature, with high expression of *Il6*, *Cxcl14*, *Lcn2*, and *Saa3*, consistent with previously described inflammatory fibroblasts[1], and was relatively enriched in *Lmna*KO$^{iEC}$ aortas (**Figure S11C-D and Table S3)**. GO analysis further confirmed enrichment for inflammatory biological processes (**Figure S12 and Table S4**).

FB5 uniquely expressed high levels of *Myh11*, *Lmod1*, and *Sox9*, suggesting that it represents a population of VSMC-like fibroblasts. This subcluster was less abundant in *Lmna*KO$^{iEC}$ aortas (**Figure S11C-D and Table S3)**.

FB6 co-expressed endothelial markers (*Pecam1*, *Icam2*) together with mesenchymal or developmental genes (*Dlx5*, *Lgr6*), consistent with an EC-like transcriptional signature suggestive of endothelial-to-mesenchymal transition. This subcluster was relatively enriched in *Lmna*KO$^{iEC}$

aortas (**Figure S11C-D and Table S3)**.

We also analyzed transcriptional differences between control and *Lmna*KO$^{iEC}$ mice within each fibroblast subcluster to identify both shared and subcluster-specific alterations (**Table S5)**. Analysis of shared differentially expressed genes (DEGs) revealed no significant changes common to all fibroblast subclusters, indicating the absence of a uniform pan-fibroblast transcriptional response to endothelial *Lmna* deletion (**Figure S13A and Table S5)**. Nevertheless, several fibroblast subclusters from *Lmna*KO$^{iEC}$ mice exhibited transcriptional changes consistent with enhanced matrix remodeling. Specifically, the ECM-associated FB0 subcluster in *Lmna*KO$^{iEC}$ mice showed enrichment for GO terms related to ossification **(Figure S13B)**, accompanied by upregulated expression of *Spp1,* which was also significantly increased in the stress-responsive FB1 subcluster (**Figure S13C and Table S5)**. Moreover, ECM organization was enriched in the inflammatory FB4 subcluster, consistent with marked upregulation of the ECM remodeling-associated genes *Postn* and *Fn1* (**Figure S13B-C and Table S5).** Collectively, these findings indicate that endothelial *Lmna* deletion induces distinct fibroblast subcluster-specific transcriptional responses that converge on enhanced ECM remodeling.

### Enhanced phenotypic VSMC modulation in *Lmna*KO$^{iEC}$ aortas

As described in the main text, we identified a non-canonical cell cluster (C4 in **Figure 6B**; 7.73% of total cells) expressing ECM-remodeling genes together with moderate levels of contractile VSMC markers, consistent with a population of phenotypically modulated VSMCs. To assess whether these cells originate from VSMCs, we conducted a pseudotime trajectory analysis using Slingshot on a refined subclustered dataset including the modulated VSMC cluster (C4) along with canonical VSMCs (C2) and ECs (C3), the two most closely related clusters based on UMAP proximity **(Figure S14A-B)**. The inferred trajectory indicated that canonical VSMCs (C2) were the most likely precursor population of C4, supporting its classification as a modulated VSMC population **(Figure S14C)**.

Subclustering of *Myh11*-enriched aortic VSMC clusters (C2 and C4 in **Figure 6B**) identified six transcriptionally distinct populations (VC0-VC5), present in both control and *Lmna*KO$^{iEC}$ aortas (**Figure S15A-B**). To further characterize these subclusters, we identified specifically enriched genes and assessed differences in their relative abundance between genotypes (**Figure S15C-D**).

VC0 and VC1 were enriched for canonical contractile VSMC markers, including *Myh11*, *Acta2*, *Lmod1*, and *Smtn* (**Figure S15C and Table S6**). GO analysis confirmed enrichment for biological processes related to actin filament assembly and cytoskeleton organization, supporting their identity as contractile VSMCs (**Figure S16 and Table S7**).

VC2 and VC3 exhibited expression profiles characteristic of phenotypically modulated VSMCs, with upregulation of *Lum*, *Dcn*, *Mmp2*, and *Lgals3*, together with downregulation of contractile and VSMC identity genes (*Myh11*, *Acta2*, *Smtn*, *Lmod1*) (**Figure S15C and Table S6**). Notably,

VC2 was enriched for *Fbln2*, *Sdc1*, and *Vcam1*, suggesting a fibromyocyte-like population involved in ECM remodeling, consistent with GO enrichment for collagen fibril and ECM organization (**Figure S15C, Figure S16, Table S6, and Table S7**). VC3 was characterized by high expression of *Spp1*, *Acan*, and *Chad*, indicative of a chondrocyte-like phenotype (**Figure S15C and Table S6**). This was corroborated by GO enrichment for pathways related to ECM remodeling, chondrocyte differentiation, and osteoblast differentiation, consistent with an osteogenic VSMC state associated with advanced atherosclerotic lesion remodeling (**Figure S16 and Table S7**). VC2 and VC3 together accounted for the majority of cells within the original modulated VSMC cluster C4 **(Figure 6B)**, indicating that this population is primarily composed of fibromyocyte- and chondrocyte-like VSMCs.

VC4 occupied an intermediate positioning in the UMAP representation, partially overlapping with both canonical VSMCs and fibromyocyte-like populations, suggesting that it represents a transitional or heterogeneous state (**Figure S15B**). The expression of genes such as *Taco1*, *Gpr39*, and *Fgd6* supports the possibility that these cells are undergoing metabolic reprogramming (**Figure S15C and Table S6**).

VC5 retained VSMC identity markers but was enriched in genes linked to mineralization, suggesting that this population represents an early transitional state toward an osteoblast-like phenotype (**Figure S15C and Table S6**).

Interestingly, the relative abundance of VSMC populations was altered in *Lmna*KO$^{iEC}$ mice, with enrichment of fibromyocyte- and chondrocyte-like populations (VC2 and VC3) and a reduction in contractile populations (VC0 and VC1) (**Figure S15D**).

We also analyzed transcriptional changes between control and *Lmna*KO$^{iEC}$ mice within each VSMC subcluster to identify both shared and subcluster-specific alterations **(Figure S17A and Table S8)**. VSMCs from *Lmna*KO$^{iEC}$ mice exhibited transcriptional signatures consistent with cellular stress and ribosomal dysregulation, as evidenced by upregulation of stress-related proteins (*Hspa1a*, *Hspa1b*) and the ribosomal component *Rpl35* **(Figure S17B and Table S8)**. Contractile VSMC clusters (VC0 and VC1) showed reduced expression of contractile genes, including *Cald1*, *Tpm1*, and *Lmod1*, together with upregulation of phenotypic modulation markers, including *S100a4*, *Lgals3*, and *Spp1,* in *Lmna*KO$^{iEC}$ aortas **(Figure S17C and Table S8)**. These findings, together with the observed reduction in contractile populations and a relative expansion of fibromyocyte- and chondrocyte-like populations, indicate enhanced phenotypic modulation of canonical VSMCs toward fibromyocyte- and chondrocyte-like states in the atherosclerotic aortas of *Lmna*KO$^{iEC}$ mice.

#### Expansion of inflammatory myeloid populations in *Lmna*KO$^{iEC}$ aortas

Next, we performed subclustering of *Ptprc* (CD45)-enriched aortic immune populations (clusters C1, C6, C8, C9, and C10 in **Figure 6B**), which identified thirteen transcriptionally distinct populations (IC0-IC12), present in both control and *Lmna*KO$^{iEC}$ mice (**Figure S18A-B**). To further characterize these subclusters, we identified specifically enriched genes and assessed differences in their relative abundance between genotypes (**Figure S18C-D and Table S9**).

As expected in an atherosclerotic environment, myeloid cells were the predominant immune population, with macrophage clusters expressing canonical markers such as *Cd68*, *Adgre1*, *Cd14,* and *Fcgr1* representing the largest cell populations (clusters IC0, IC1, IC2, and IC8) (**Figure S18B-C and Table S9**).

IC12 also co-expressed macrophage markers, including *Cd68,* together with canonical contractile VSMC genes (*Acta2*, *Tagln,* and *Myh11*), suggesting a population of phenotypically modulated VSMCs with macrophage-like features (**Figure S18C and Table S9**).

Cells expressing canonical dendritic cell (DC) markers, including *Flt3* and *Zbtb46,* were also prominently represented (IC3, IC5, and IC9) (**Figure S18B-C and Table S9**).

Distinct lymphoid populations were also identified, including T cells (IC4 and IC7), characterized by expression of *Cd3d*, *Cd3e,* and *Cd3g*; natural killer cells (IC6), expressing *Nkg7* and *Klrk1*; B cells (IC10), expressing *Cd79a*, *Cd79b,* and *Mzb1;* and innate lymphoid cells (ILCs) (IC11), characterized by *Gata3*, *Dach2,* and *Il1rl1* expression (**Figure S18C and Table S9**). Interestingly, the relative abundance of immune subpopulations differed between genotypes, with *Lmna*KO$^{iEC}$ aortas exhibiting expansion of myeloid populations, including macrophage clusters IC0, IC1, and IC8, together with DC clusters IC3, IC5 and IC9 (**Figure S18D**).

To further characterize macrophage heterogeneity, we subclustered *Cd68*$^{+}$ macrophages, identifying seven transcriptionally distinct populations (MC0-MC6) (**Figure S19A-B**). To study these subclusters, we identified specifically enriched genes and assessed differences in their relative abundance between genotypes (**Figure S19C-D and Table S10**).

MC0 was enriched for genes associated with foam cell formation and lipid handling, including *Lgals3*, *Trem2*, *Lpl*, *Ctsd*, and *Cd9,*[2] and was markedly expanded in *Lmna*KO$^{iEC}$ aortas (**Figure S19C-D and Table S10**). GO analysis revealed enrichment for cholesterol and lipid metabolism, supporting its identity as a foam cell-like macrophage population **(Figure S20 and Table S11).**

MC2 expressed canonical tissue-resident macrophage markers, including *Lyve1*, *Timd4*, *F13a1*, *Cd163*, and *Mrc1,* consistent with a resident macrophage phenotype[3] **(Figure S19C and Table S10)**.

In contrast, MC1, MC3, and MC4 were enriched for *Ccr2*, a marker commonly associated with macrophages derived from recruited monocytes[3], and all three populations were expanded in *Lmna*KO$^{iEC}$ aortas (**Figure S19C-D and Table S10**).

MC1 displayed a robust inflammatory profile, with high expression of cytokines including *Ccl3*, *Ccl4*, *Ccl12*, *Tnf*, and *Fcgr1*, together with enrichment for GO terms related to immune responses **(Figure S19C, Figure S20, Table S10, and Table S11)**.

MC3 expressed markers associated with recently recruited monocytes, including *Ly6c2*, *Plac8*, and *Trem1*, together with proliferation-associated genes like *Mki67* and *Top2a* **(Figure S19C and Table S10)**, suggesting a mixed population of recently recruited monocytes and proliferating inflammatory macrophages. Pseudotime trajectory analysis further identified MC3 as the most likely precursor of the recruited ($Ccr2^+$) macrophage populations MC1 and MC4, as well as the foam cell-like population MC0, supporting its role as the source of differentiated monocyte-derived macrophages **(Figure S21)**.

MC4 expressed DC markers, including *Cd209a*, while retaining classical recruited macrophage markers **(Figure S19C and Table S10).** GO analysis revealed enrichment for antigen presentation pathways, supporting its identity as a monocyte-derived DC population **(Figure S20 and Table S11)**.

MC5 expressed contractile VSMC markers (*Acta2*, *Tagln*, and *Myh11*) together with macrophage markers such as *Cd68* and the VSMC secretory phenotype-associated gene *Lum*, suggesting a population of phenotypically modulated VSMCs with macrophage-like features **(Figure S19C and Table S10)**.

MC6 exhibited strong expression of classical DC markers (*Flt3*, *Zbtb46*, and *Batf3*) together with apoptosis-related genes (*Apaf1* and *Casp3*), consistent with a terminally differentiated apoptotic monocyte-derived DC population. MC6 was also relatively enriched in *Lmna*KO$^{iEC}$ aortas (**Figure S19C-D and Table S10**).

Overall, *Lmna*KO$^{iEC}$ aortas exhibited expansion of recruited *Ccr2*$^+$ macrophages (MC1, MC3, MC4), foam cell-like macrophages (MC0), and antigen-presenting DC-like populations (MC6), consistent with increased monocyte recruitment and an enhanced pro-atherogenic inflammatory environment **(Figure S19D)**.

We next analyzed transcriptional differences between control and *Lmna*KO$^{iEC}$ mice within each macrophage subcluster to identify both shared and subcluster-specific alterations **(Figure S22A and Table S12)**. Macrophages from *Lmna*KO$^{iEC}$ mice exhibited widespread downregulation of *Mrc1*, a marker of anti-inflammatory macrophages, together with upregulation of *Hexb*, a lysosomal enzyme involved in lipid metabolism and immune activation, consistent with a shift toward a more inflammatory and metabolically stressed state (**Figure S22B and Table S12**). This shift was further supported by increased expression of inflammatory genes, including *Spp1*, *Socs3*, and *Cd52*, together with reduced expression of *Maf*, a transcription factor associated with anti-inflammatory macrophage programming, particularly within the inflammatory (MC1) and resident (MC2) macrophage populations **(Figure S22C and Table S12)**. Consistent with these findings, inflammatory MC1 macrophages from *Lmna*KO$^{iEC}$ mice also exhibited elevated expression of inflammatory cytokines and chemokines, including *Il1b*, *Ccrl2*, and *Cxcl16*, as well as *Pim1*, a kinase associated with macrophage activation, together with reduced expression of *Pf4* and *Gas6*, two genes associated with anti-inflammatory and pro-resolving macrophage functions[4,5] (**Figure S22C and Table S12**). A similar pattern was observed in resident macrophages (MC2), characterized by reduced *Cd163* expression and increased *Lgals3* expression (**Figure S22C and Table S12).** Overall, these findings indicate both expansion of pro-atherogenic myeloid populations and a shift toward a more inflammatory and less reparative macrophage phenotype in *Lmna*KO$^{iEC}$ mice with endothelial-specific lamin A/C deletion.

### Supplemental Discussion

The high-resolution subclustering performed on our scRNA-seq analysis of the aorta provides additional insight into how endothelial lamin A/C deficiency reshapes the multicellular compartments of the atherosclerotic vessel wall beyond the onset of endothelial dysfunction. Endothelial-specific *Lmna* deletion promoted distinct transcriptional adaptations across fibroblasts, VSMCs, immune cells, and ECs. Despite this cellular heterogeneity, these responses converged on common biological processes, including ECM remodeling, inflammatory activation, and disruption of vascular homeostasis, all of which are recognized hallmarks of vascular aging[6] and atherosclerosis[7].

Although fibroblasts and VSMCs were not direct targets in our endothelial-specific model, their transcriptional adaptations further support their contribution to the cardiovascular alterations observed in *Lmna*KO$^{iEC}$ mice. Fibroblast subpopulations enriched in inflammatory mediators and matrix-remodeling genes (e.g; *Il6*, *Postn,* and *Fn1*) were expanded, whereas quiescent fibroblast populations were reduced. Likewise, VSMCs exhibited increased representation of phenotypically modulated, osteogenic, and fibromyocyte-like states, together with reduced expression of contractile markers, changes that are characteristic of vascular aging and atherosclerotic remodeling.

The immune compartment also underwent extensive remodeling in *Lmna*KO$^{iEC}$ aortas, which exhibited expansion of recruited *Ccr2*$^{+}$ macrophages, foam cell-like macrophages, and monocyte-derived DCs, together with a relative reduction in resident macrophage populations and widespread activation of inflammatory transcriptional programs. The accompanying loss of homeostatic macrophage markers and increased expression of inflammatory mediators further support a shift toward a pro-inflammatory myeloid landscape. These alterations are consistent with enhanced monocyte recruitment and local differentiation within the plaque and likely contribute to the increased atherosclerotic burden observed in *Lmna*KO$^{iEC}$ mice[8].

Interestingly, these findings complement our recent demonstration that physiological aging is also associated with lamin A/C downregulation in circulating leukocytes, where reduced lamin A/C expression directly promotes atherosclerosis[9]. Together, these studies raise the possibility that age-associated lamin A/C decline in both endothelial and hematopoietic compartments cooperatively amplifies vascular inflammation and disease progression during aging. Although the present study demonstrates profound remodeling of the vascular immune landscape secondary to endothelial lamin A/C deficiency, future studies should determine whether endothelial lamin A/C loss directly promotes leukocyte adhesion, recruitment, and transendothelial migration, and whether simultaneous lamin A/C decline in endothelial and immune cells exerts synergistic effects on vascular inflammation and atherosclerosis.

### Legends for Supplemental Tables

**Table S1.** Primers used for quantitative real-time PCR.

**Table S2.** Differentially expressed genes in each cluster compared with all other clusters from the global clustering analysis.

**Table S3.** Differentially expressed genes in each fibroblast subcluster compared with all other fibroblast subclusters.

**Table S4.** Gene Ontology (GO) biological processes enriched in each fibroblast subcluster.

**Table S5.** Differential gene expression analysis between *Lmna*KO$^{iEC}$ mice and control mice within each fibroblast subcluster.

**Table S6.** Differentially expressed genes in each VSMC subcluster compared with all other VSMC subclusters.

**Table S7.** Gene Ontology (GO) biological processes enriched in each VSMC subcluster.

**Table S8.** Differential gene expression analysis between *Lmna*KO$^{iEC}$ mice and control mice within each VSMC subcluster.

**Table S9.** Differentially expressed genes in each immune cell subcluster compared with all other immune cell subclusters.

**Table S10.** Differentially expressed genes in each macrophage subcluster compared with all other macrophage subclusters.

**Table S11.** Gene Ontology (GO) biological processes enriched in each macrophage subcluster.

**Table S12.** Differential gene expression analysis between *Lmna*KO$^{iEC}$ mice and control mice within each macrophage subcluster.

**Table S13.** Differentially expressed genes in each EC subcluster compared with all other subclusters.

**Table S14.** Gene Ontology (GO) biological processes enriched in each EC subcluster.

**Table S15.** Differential gene expression analysis between *LmnaKO$^{iEC}$* and control mice within each EC subcluster.

**Table S16.** Ingenuity Pathway Analysis of predicted canonical pathway alterations in *LmnaKO$^{iEC}$* ECs compared with control ECs.

**Table S17.** Ingenuity Pathway Analysis of predicted upstream regulator activity in *LmnaKO$^{iEC}$* ECs compared with control ECs.

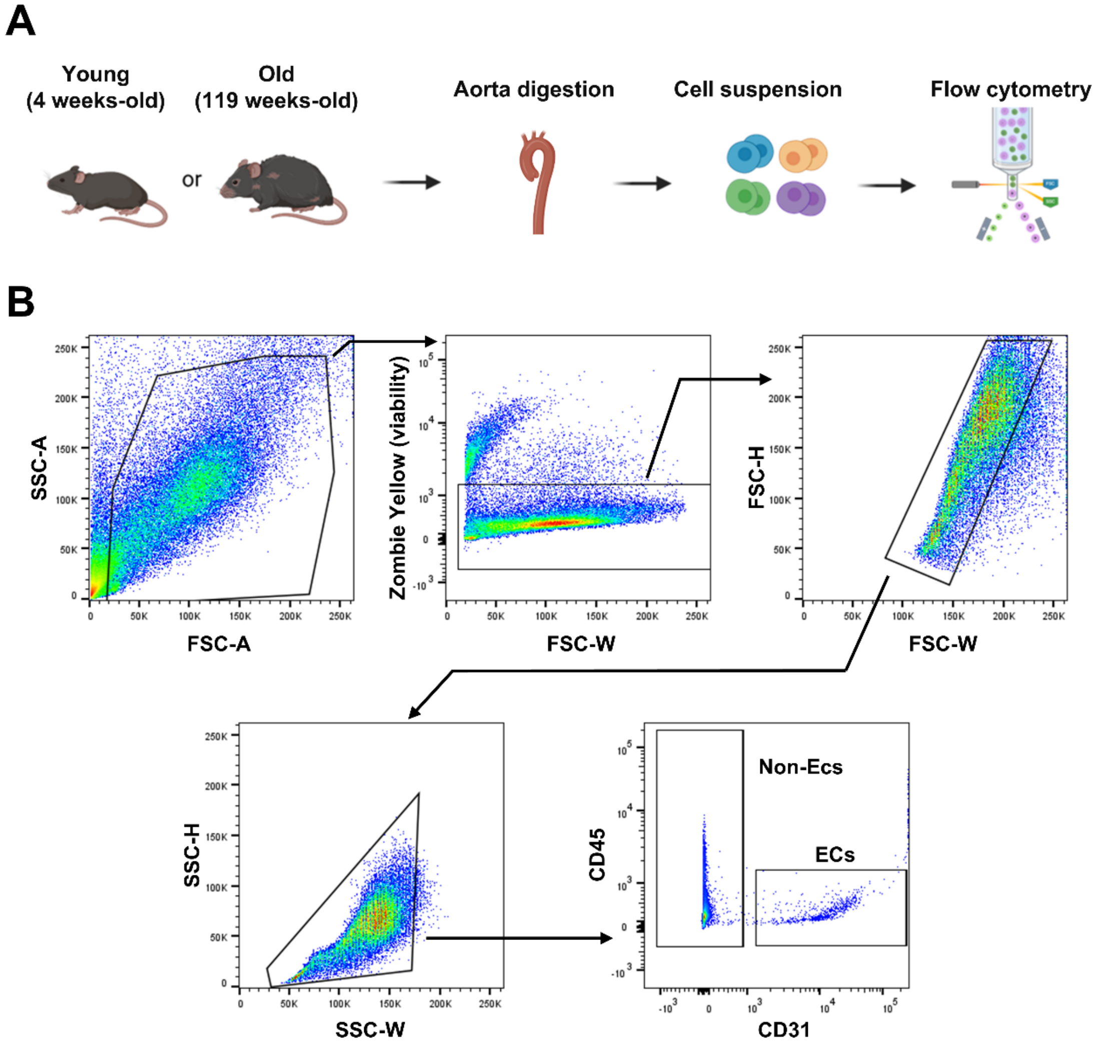


**Supplemental Figure S1. Flow cytometry gating strategy for analysis of lamin A/C expression in aortic cell populations during mouse aging. (A)** Experimental design. Young (4-week-old) and aged (119-week-old) mice were euthanized, and aortas were collected and enzymatically digested to obtain single-cell suspensions for flow cytometry. **(B)** Gating strategy for analysis of lamin A/C expression in endothelial cells (ECs) ($CD31^{+}CD45^{-}$) and non-EC populations ($CD31^{-}$).

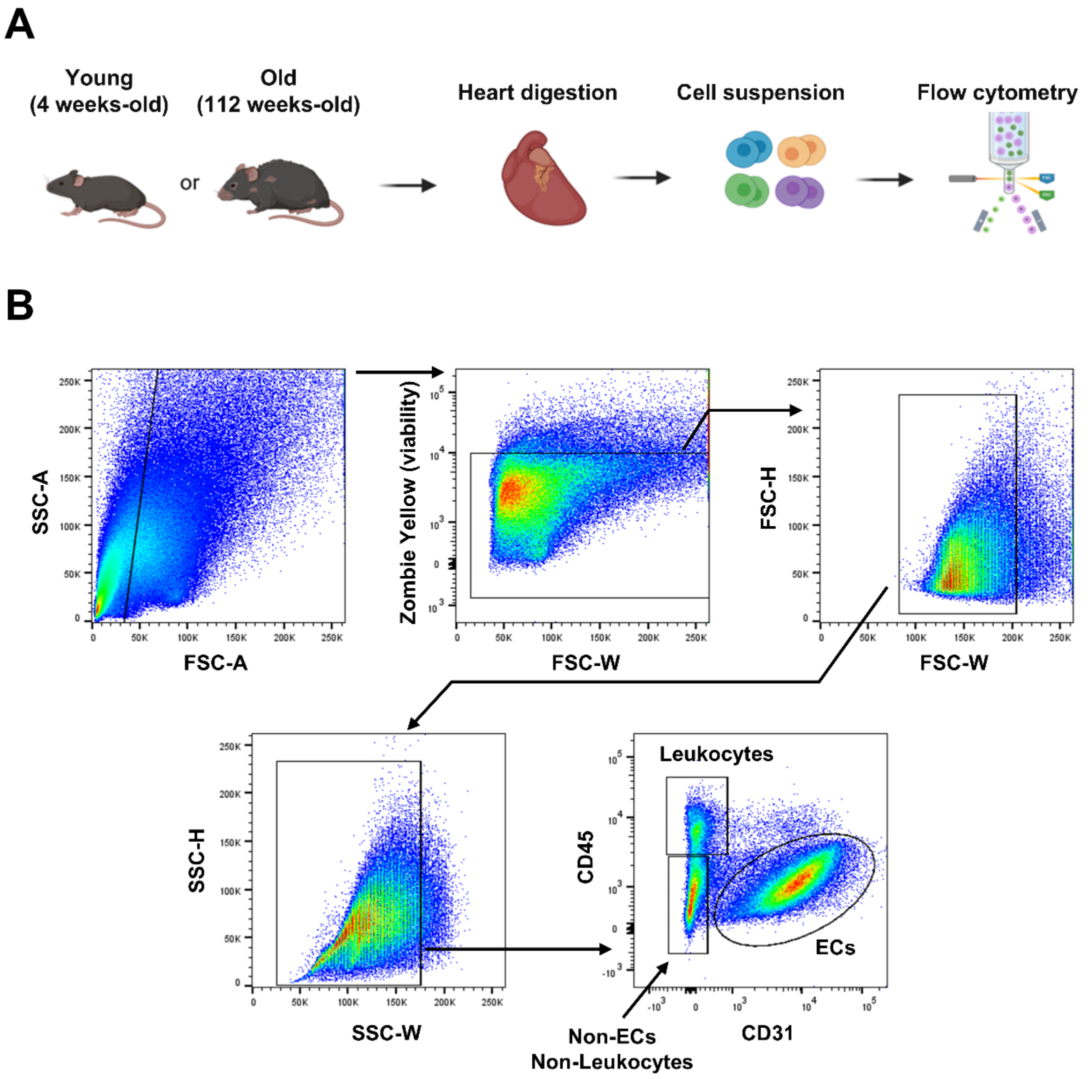


**Supplemental Figure S2. Flow cytometry gating strategy for analysis of lamin A/C expression in cardiac cell populations during mouse aging. (A)** Experimental design. Young (4-week-old) and aged (112-week-old) mice were euthanized, and hearts were collected and enzymatically digested to obtain single-cell suspensions for flow cytometry. **(B)** Gating strategy for analysis of lamin A/C expression in endothelial cells (ECs) ($CD31^{+}CD45^{-}$), leukocytes ($CD31^{-}CD45^{+}$), and non-ECs/non-leukocytes populations ($CD31^{-}CD45^{-}$).

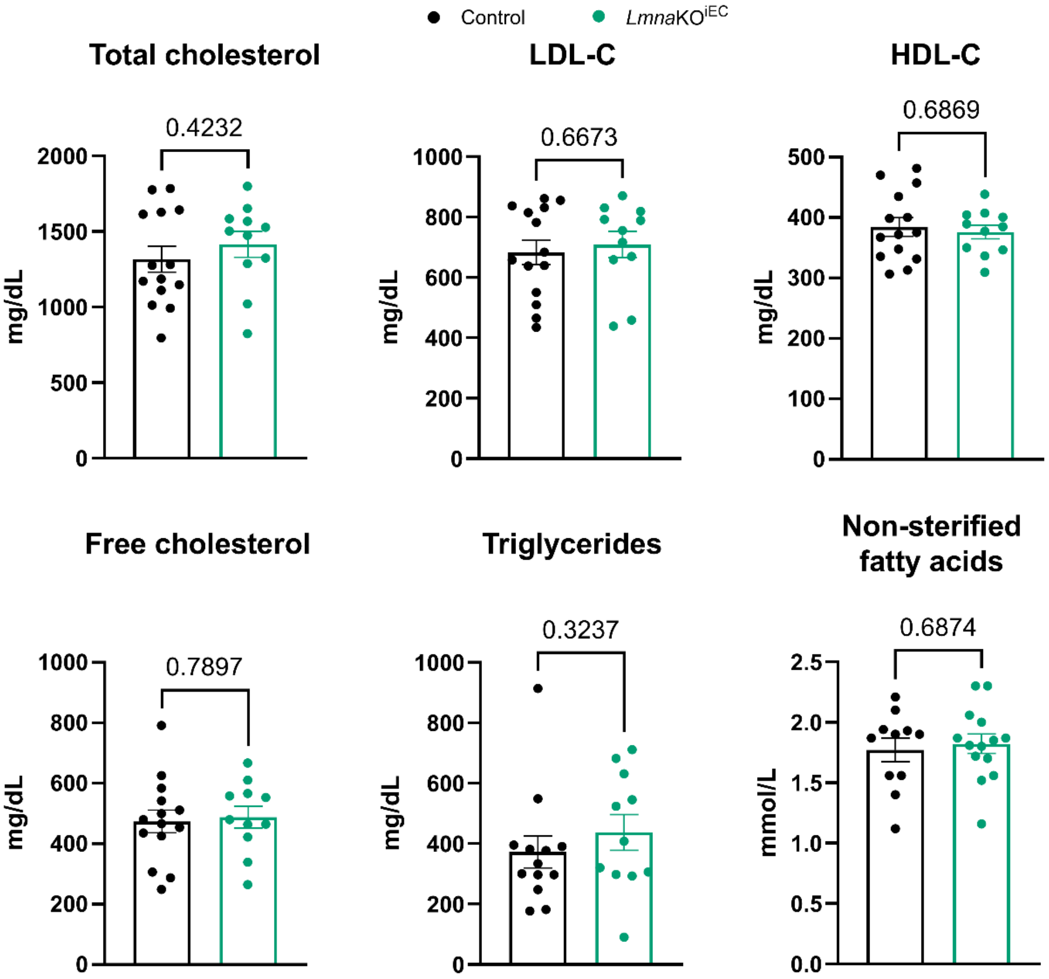


**Supplemental Figure S3. Circulating lipid profile in control and *Lmna*KO$^{iEC}$ mice.** Mice were fed a high-fat diet for 12 weeks and euthanized at 9 months of age. Graphs show serum concentrations of total cholesterol, low-density lipoprotein cholesterol (LDL-C), high-density lipoprotein cholesterol (HDL-C), free cholesterol, triglycerides, and non-esterified fatty acids (control, n=14; *Lmna*KO$^{iEC}$, n=11). Statistical significance was assessed by unpaired Student's t-test.

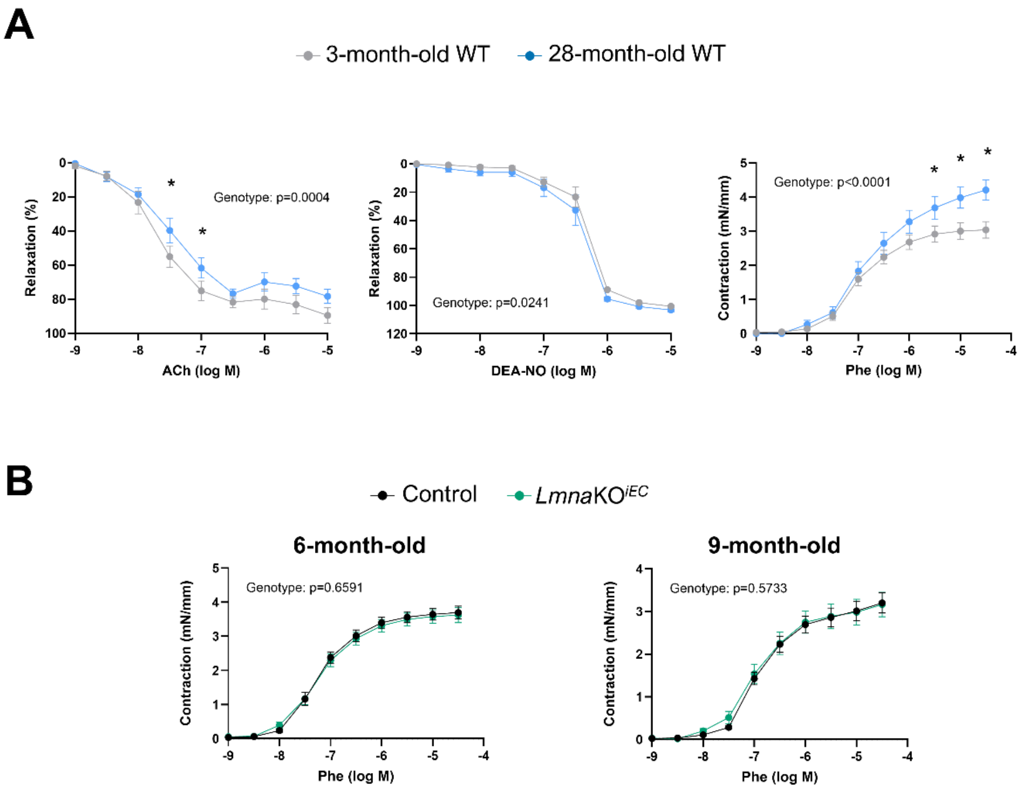


**Supplemental Figure S4. *Ex vivo* analysis of thoracic aorta vasoreactivity in wild-type, *Lmna*KO$^{iEC}$, and control mice by wire myography. (A)** Endothelium-dependent vasorelaxation in response to increasing doses of acetylcholine (ACh), endothelium-independent vasorelaxation in response to the NO donor DEA-NO, and vasoconstriction in response to increasing doses of phenylephrine (Phe) in young (3-month-old; n=9) and aged (28-month-old; n=9) wild-type mice fed normal chow. **(B)** Vasoconstriction in response to increasing doses of Phe in 6- and 9-month-old *Lmna*KO$^{iEC}$ mice and age-matched controls (6-month-old cohort: n=20; 9-month-old cohort: control, n=11; *Lmna*KO$^{iEC}$, n=9). Mice were fed high-fat diet for 12 weeks. Statistical significance was assessed by two-way ANOVA. Asterisks indicate $P$<0.05 in the multiple-comparison analysis at each dose.

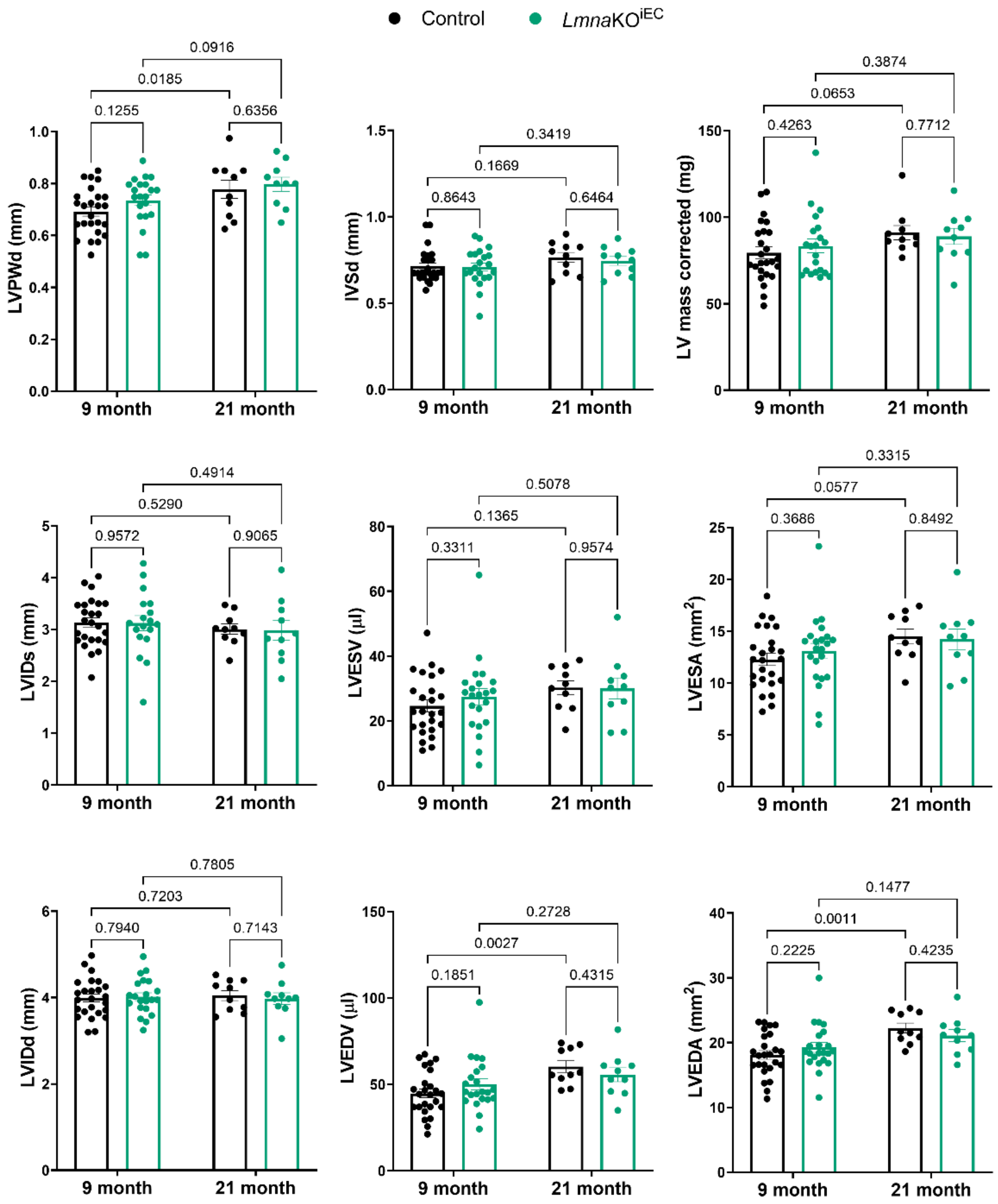


**Supplemental Figure S5. Echocardiographic assessment of left ventricular morphology in control and *Lmna*KO$^{iEC}$ mice.** Echocardiography was performed in 9- and 21-month-old mice fed a high-fat diet for 12 and 6 weeks, respectively (9-month-old cohort: control, n=24; *Lmna*KO$^{iEC}$, n=22; 21-month-old cohort: n=10). Statistical significance was assessed by two-way ANOVA. **Abbreviations: LVPWd,** left ventricular posterior wall thickness in diastole; **IVSd, I**nterventricular septal thickness in diastole; **LV mass (corrected),** left ventricular mass with geometric correction; **LVIDs,** left ventricular internal diameter in systole; **LVESV,** left ventricular end-systolic volume; **LVESA,** left ventricular end-systolic area; **LVIDd,** left ventricular internal diameter in diastole; **LVEDV,** left ventricular end-diastolic volume; **LVEDA,** left ventricular end-diastolic area.

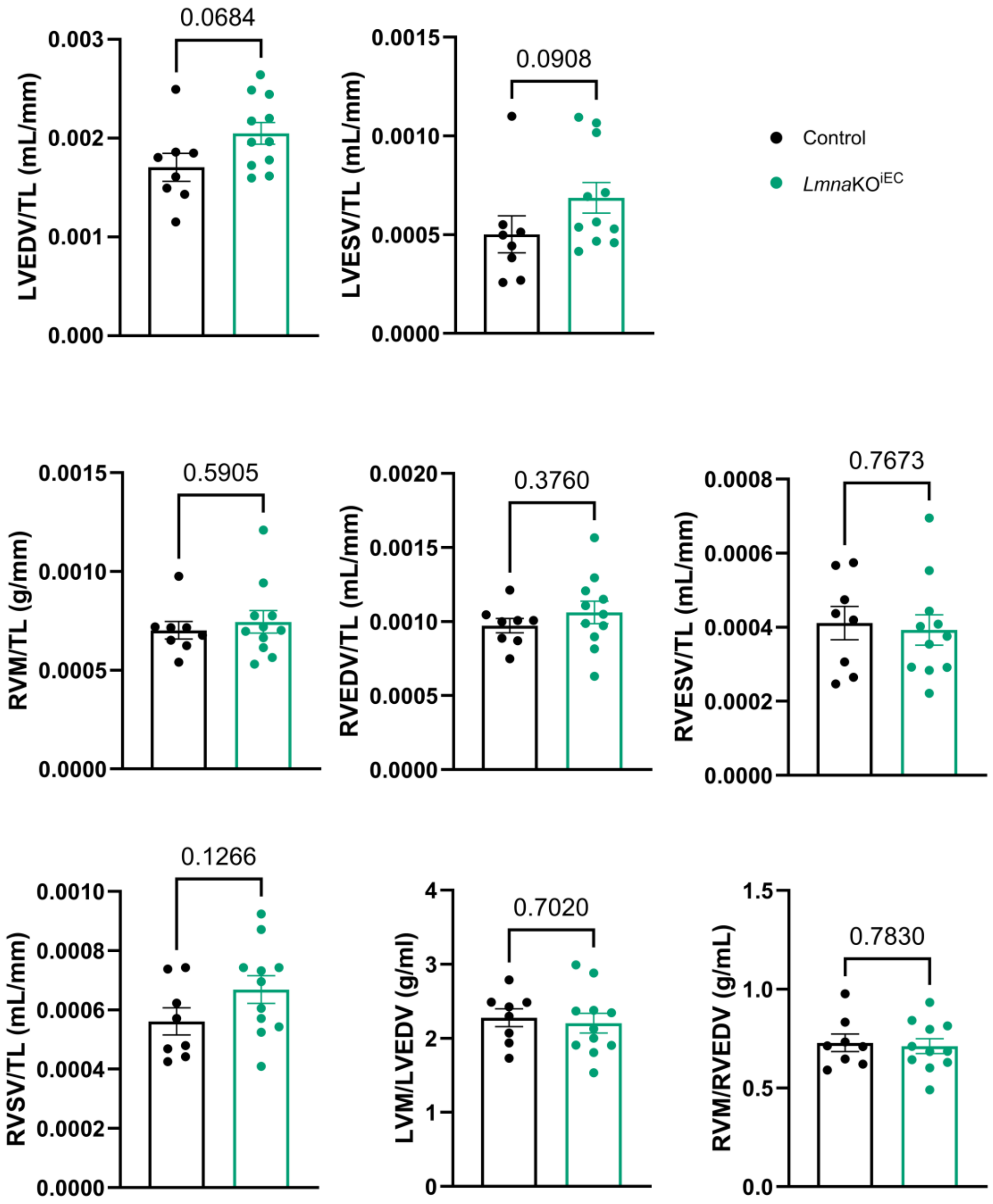


**Supplemental Figure S6. Cardiac cine magnetic resonance imaging (MRI)-derived morphological parameters in control and *Lmna*KO$^{iEC}$ mice.** Cardiac cine MRI was performed in 9-month-old mice fed a high-fat diet for 12 weeks (control, n=8; *Lmna*KO$^{iEC}$, n=11). All parameters were normalized to tibia length, except the LVM/LVEDV and RVM/RVEDV ratios. Statistical significance was assessed by unpaired Student's t-test or Mann-Whitney U test. **Abbreviations: LVEDV,** left ventricular end-diastolic volume; **LVESV,** left ventricular end-systolic volume; **RVM,** right ventricular mass; **RVEDV,** right ventricular end-diastolic volume; **RVESV,** right ventricular end-systolic volume; **RVSV,** right ventricular stroke volume; **LVM/LVEDV,** ratio of left ventricular mass to left ventricular end-diastolic volume; **RVM/RVEDV,** ratio of right ventricular mass to right ventricular end-diastolic volume.

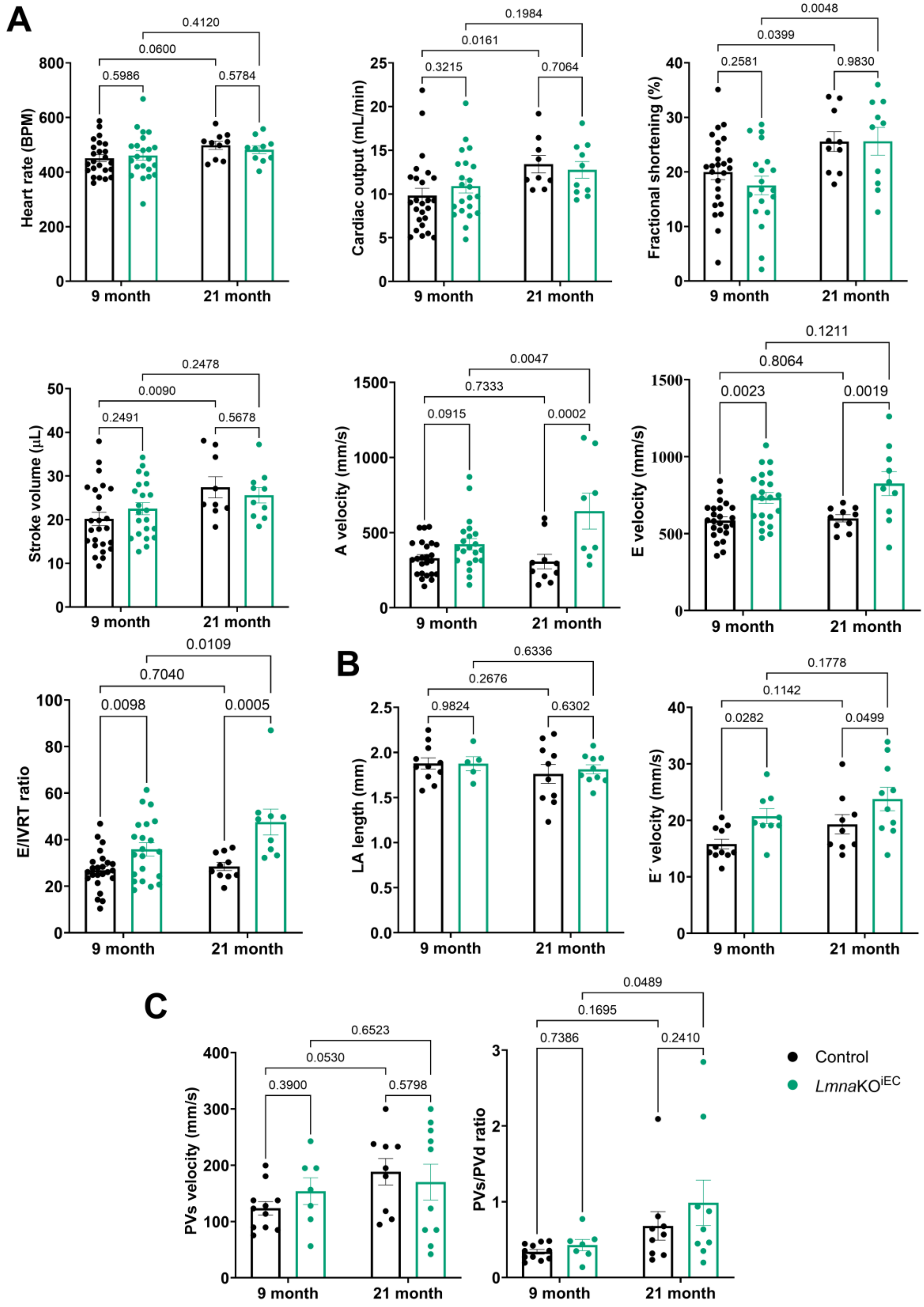


**Supplemental Figure S7. Echocardiographic assessment of systolic and diastolic function in control and *Lmna*KO$^{iEC}$ mice. (A-C)** Echocardiography was performed in 9- and 21-month-old mice fed a high-fat diet for 12 and 6 weeks, respectively. Panel **A**: 9-month-old cohort: control, n=24; *Lmna*KO$^{iEC}$, n=22; 21-month-old cohort: n=10. Panel **B and C**: 9-month-old cohort: control, n=11; *Lmna*KO$^{iEC}$, n=9; 21-month-old cohort: n=10. Statistical significance was assessed by two-way ANOVA. **Abbreviations: BPM,** beats per minute; **A velocity,** peak late (atrial) transmitral flow velocity; ; **E velocity,** peak early transmitral flow velocity; **IVRT,** isovolumetric relaxation time; **LA length,** left atrial length; **E′ velocity,** peak early diastolic myocardial velocity at the mitral annulus; **E/E′,** index of left ventricular filling pressure; **PVs velocity,** peak systolic pulmonary venous flow velocity; **PVs/PVd,** ratio of systolic to diastolic pulmonary venous flow.

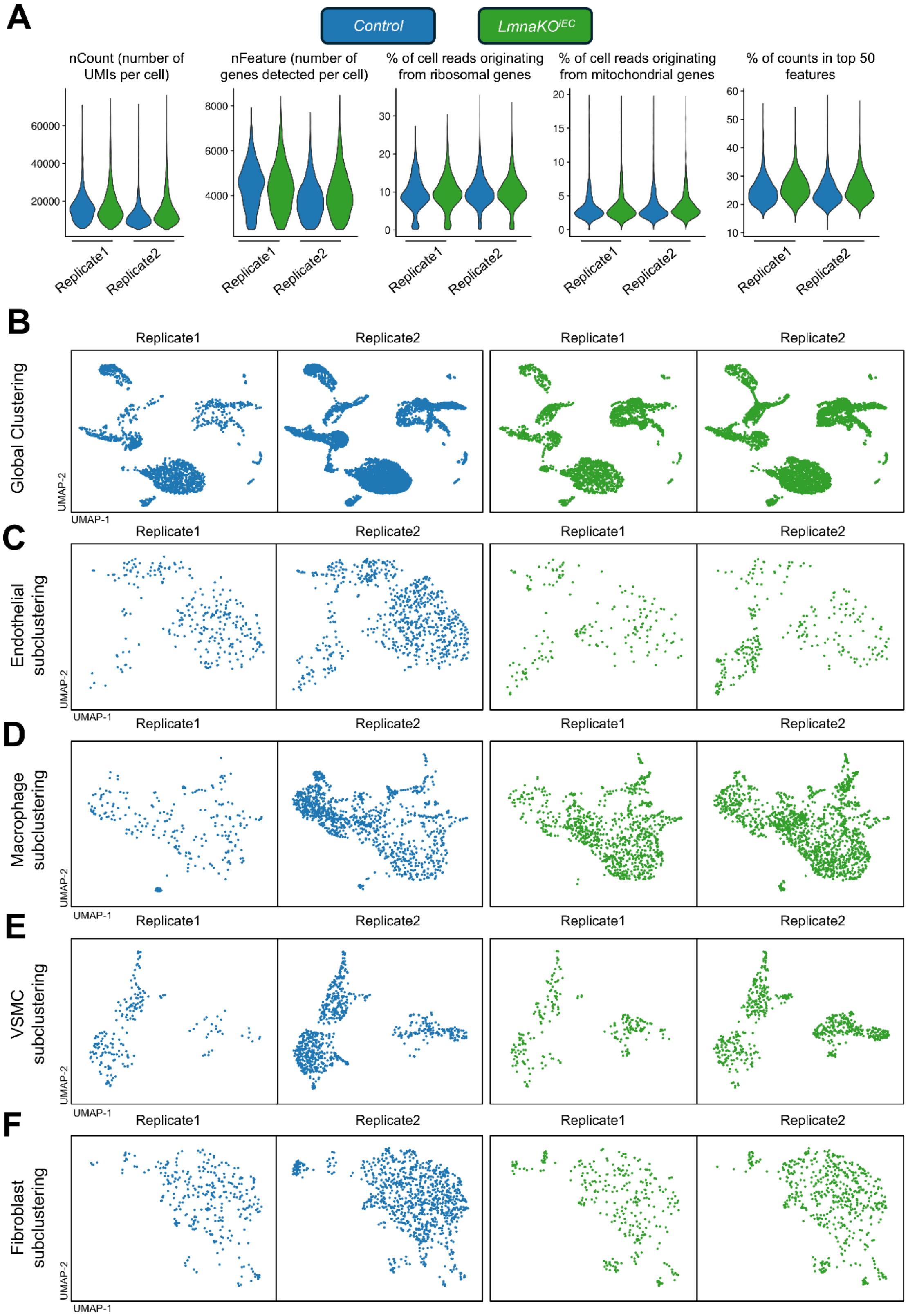


**Supplemental Figure S8. Quality control metrics and sample representation in single-cell RNA sequencing analysis. (A)** Violin plots showing quality control metrics for each sample, grouped by genotype and biological replicate. **(B)** Uniform manifold approximation and projection (UMAP) visualization of the global clustering for each replicate, colored by genotype. **(C-F)** UMAP projections of subclustered cell populations, colored by genotype and replicate, showing endothelial cells in **(C)**, macrophages in **(D)**, vascular smooth muscle cells (VSMCs) in **(E)**, and fibroblasts in **(F)**.

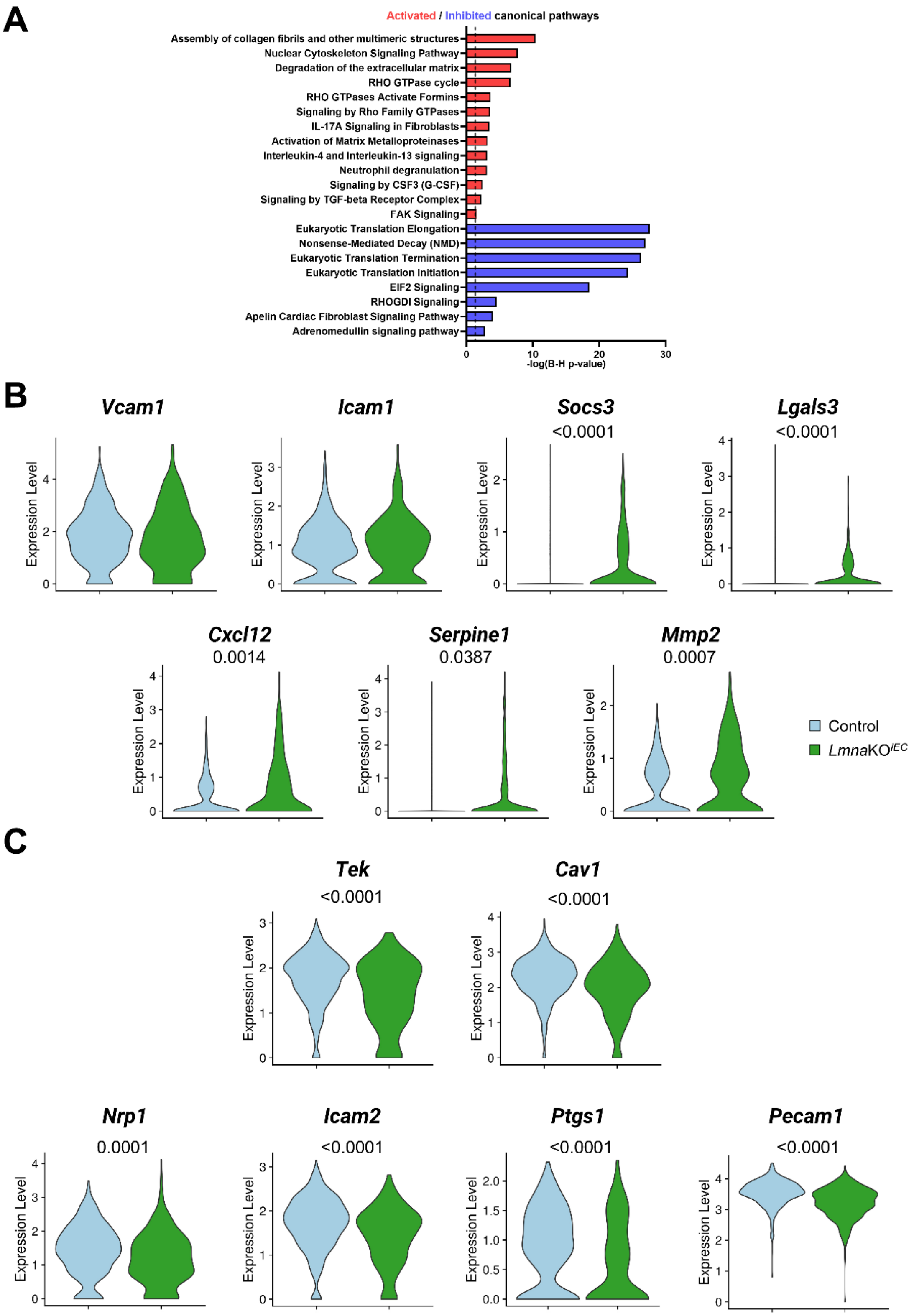


**Supplemental Figure S9. Gene expression changes in aortic endothelial cells (ECs) from *Lmna*KO$^{iEC}$ mice. (A)** Ingenuity pathway analysis activation *z*-scores of selected upstream regulators significantly altered after Benjamini-Hochberg correction (P<0.05). Activated and inhibited regulators are shown in red and blue, respectively. **(B)** Violin plots showing the expression of selected genes related to inflammation and endothelial activation by genotype in ECs. **(C)** Violin plots showing the expression of selected genes related to endothelial homeostasis and mechanosensing by genotype in ECs. Statistical analyses were performed using the Wilcoxon rank-sum test. Only significant p values (p<0.05) are shown.

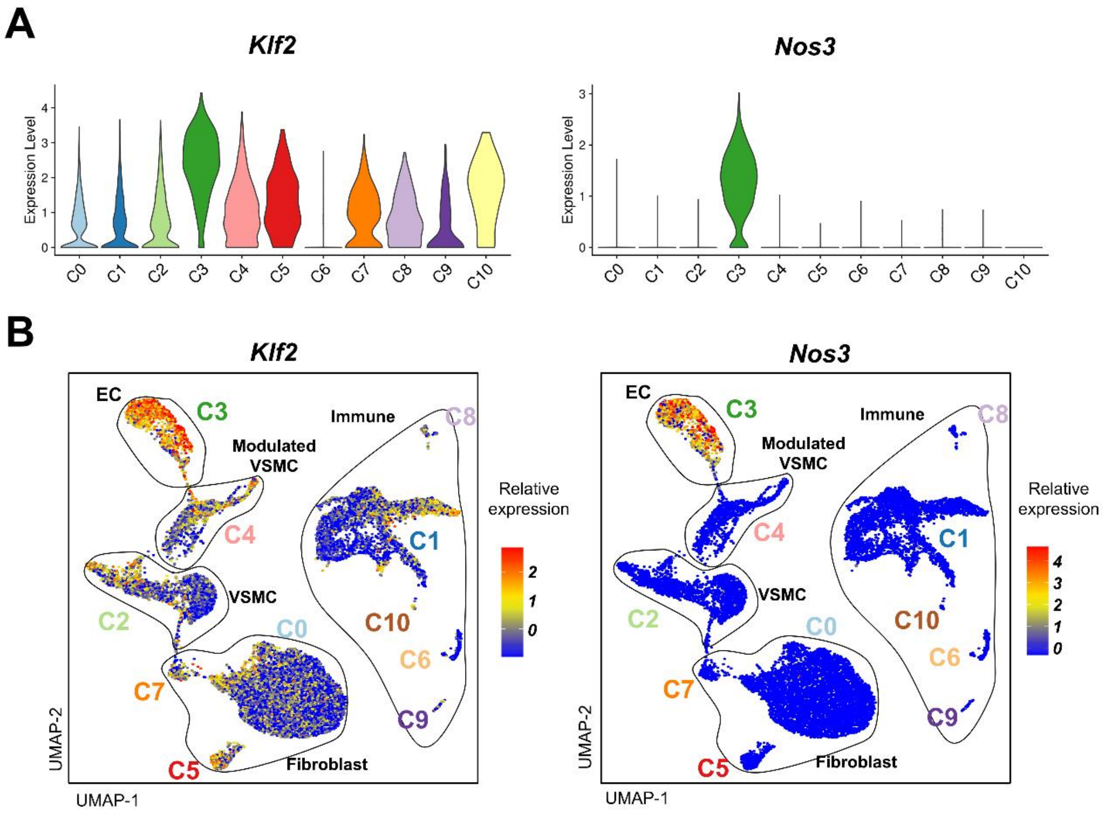


**Supplemental Figure S10. Endothelial cell (EC)-enriched expression of *Klf2* and *Nos3* in the aortic scRNA-seq dataset.** Violin plots **(A)** and Uniform manifold approximation and projection (UMAP) visualization **(B)** showing *Klf2* and *Nos3* expression across the different cell populations in the global scRNA-seq clustering.

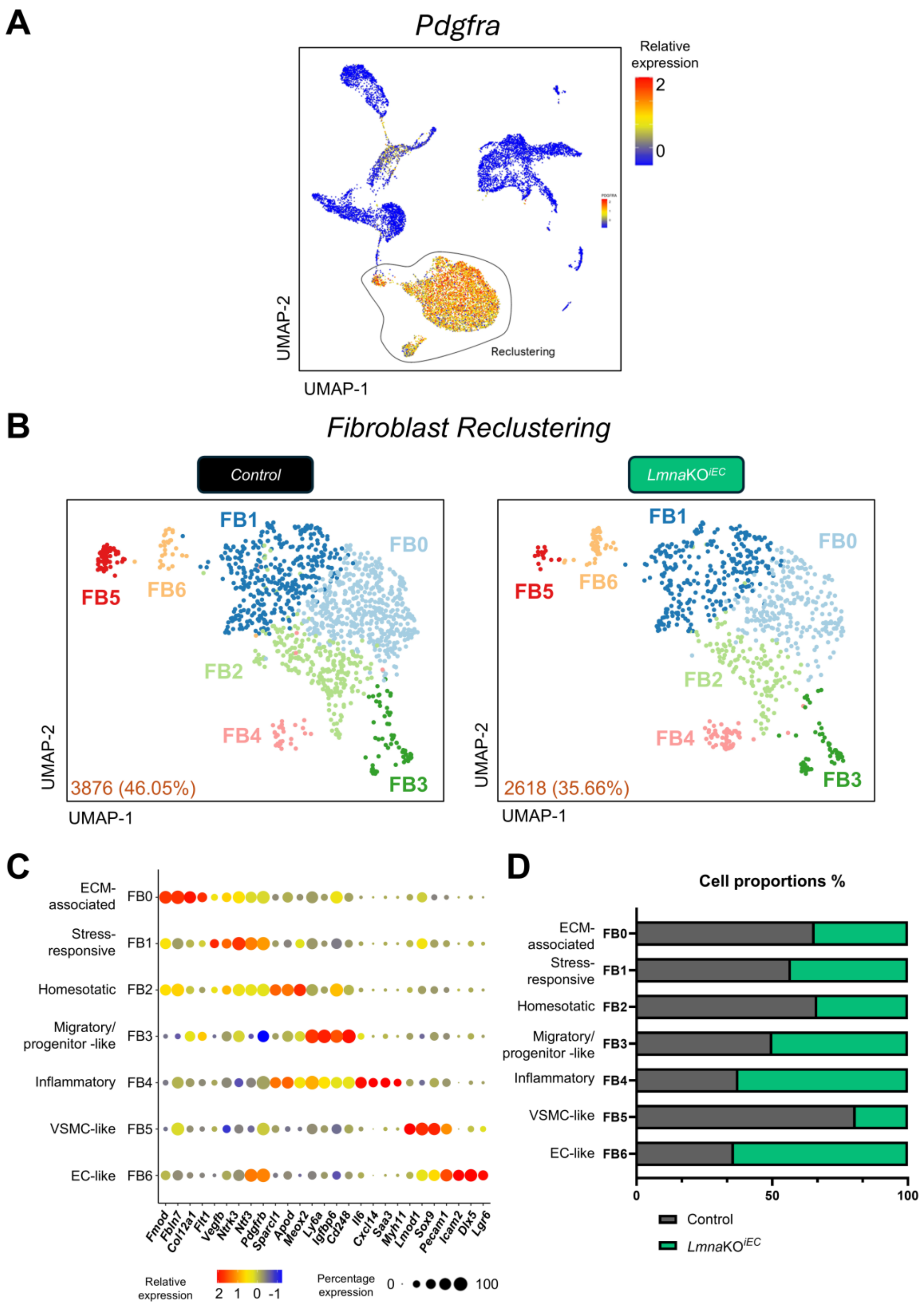


**Supplemental Figure S11. Reclustering and characterization of fibroblast populations from control and *Lmna*KO$^{iEC}$ aortas. (A)** Uniform manifold approximation and projection (UMAP) visualization of the expression of the fibroblast marker *Pdgfra* across all unsupervised clusters. The lasso indicates the cells included in the reclustering. **(B)** UMAP visualization of fibroblast reclustering in control and *Lmna*KO$^{iEC}$ mice. **(C)** Dot plot showing the average expression and the percentage of cells expressing selected genes across fibroblast subclusters. **(D)** Proportions of cells in each fibroblast subcluster by genotype. **Abbreviations: ECM**, extracellular matrix; **VSMC,** vascular smooth muscle cell; **EC,** endothelial cell.

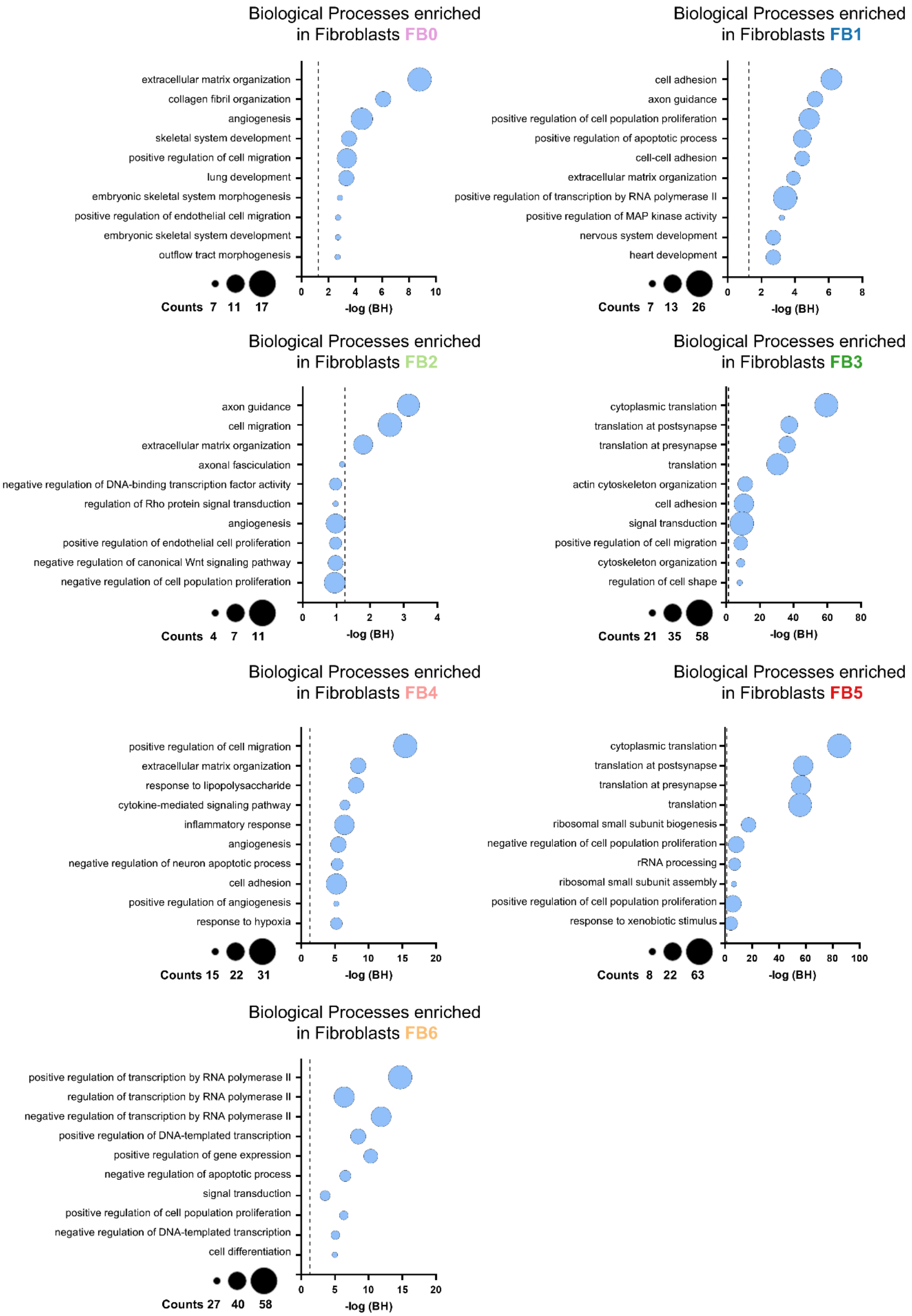


**Supplemental Figure S12. Top 10 enriched Gene Ontology (GO) biological processes for each fibroblast subcluster.** Bubble plots showing the significance and gene count for the top 10 enriched GO biological processes in each fibroblast subcluster. Bubble size indicates the number of genes associated with each GO term. The dashed line indicates the Benjamini-Hochberg adjusted P value threshold of 0.05. **Abbreviations: MAP**, mitogen-activated protein.

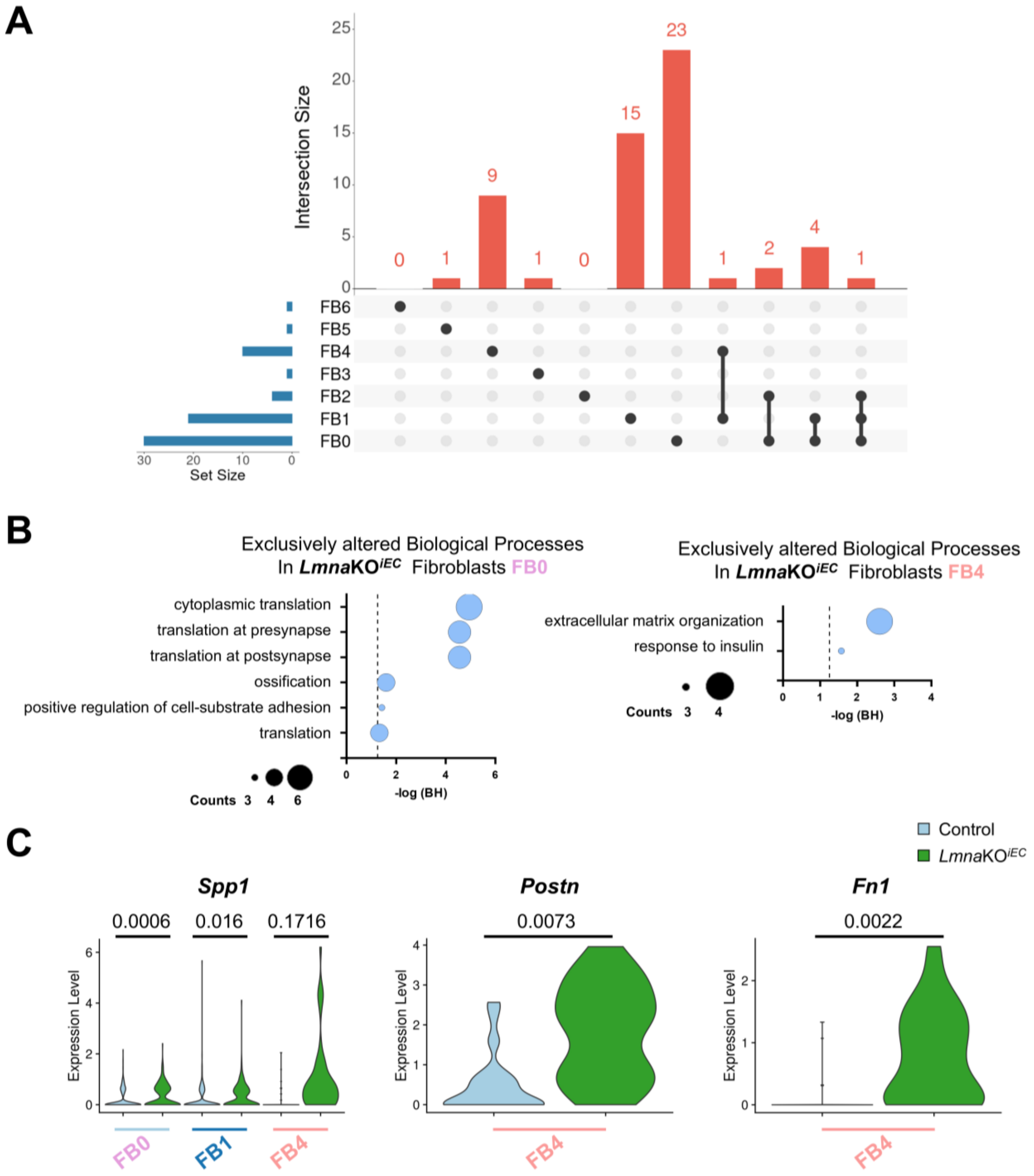


**Supplemental Figure S13. Gene expression changes in aortic fibroblast subclusters from *Lmna*KO$^{iEC}$ mice. (A)** UpSet plot showing the number of differentially expressed genes (DEGs) specific to each subcluster or shared among subclusters. Only intersections involving two or more subclusters with at least one shared DEG are shown. Set size indicates the total number of DEGs identified in each fibroblast subcluster. **(B)** Gene Ontology (GO) biological process enrichment analysis of DEGs in *Lmna*KO$^{iEC}$ fibroblast subclusters compared with control mice. **(C)** Violin plots showing the expression of selected genes by genotype across fibroblast subclusters. Statistical analyses were performed by Wilcoxon rank-sum test.

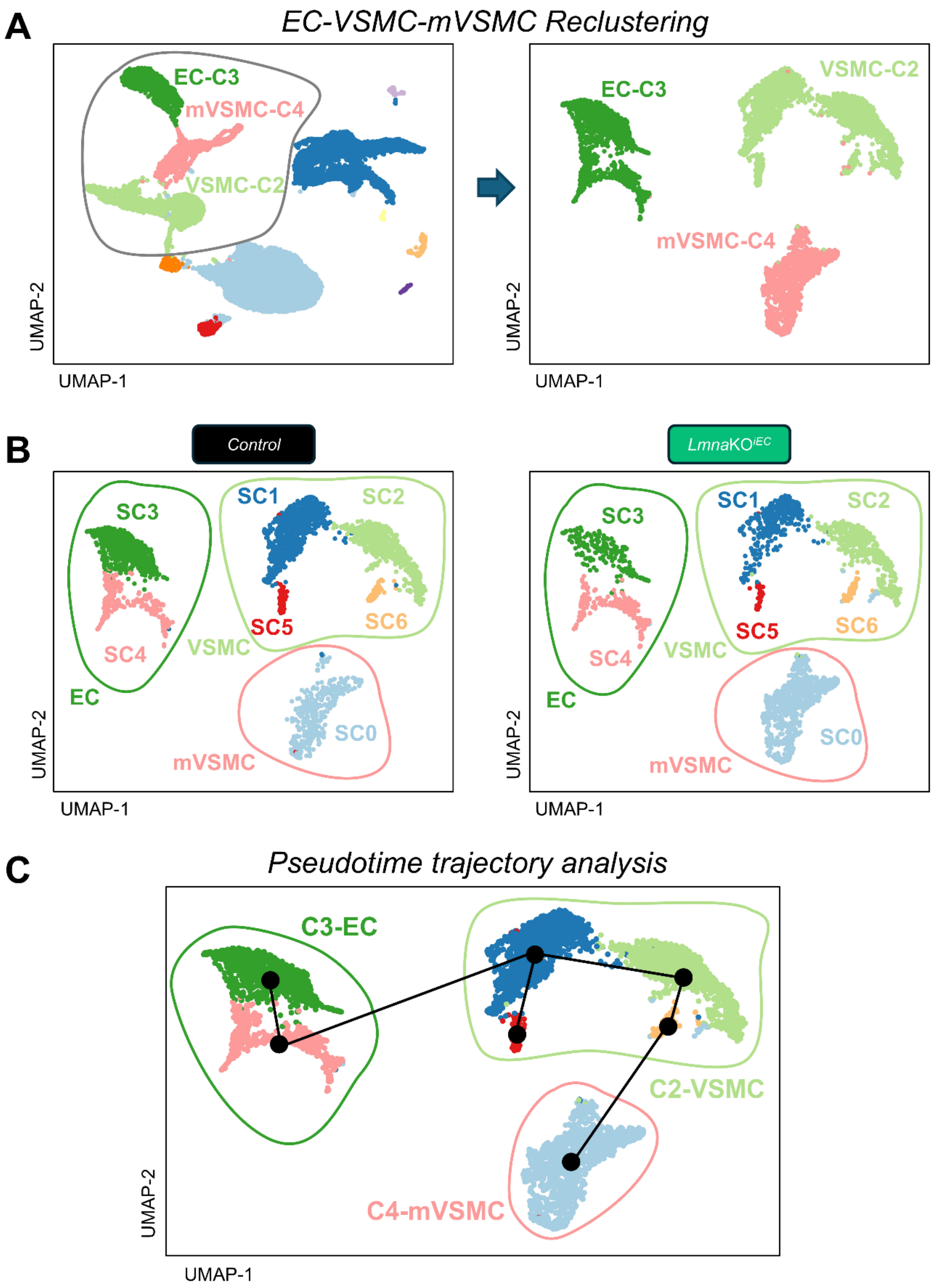


**Supplemental Figure S14. Pseudotime trajectory analysis after reclustering endothelial cells (EC), vascular smooth muscle cells (VSMC), and modulated VSMCs (mVSMC). (A)** Uniform manifold approximation and projection (UMAP) visualization of all unsupervised clusters and the reclustering of ECs, VSMCs, and mVSMCs, with the original subcluster identity indicated by color. The lasso indicates the cells included in the reclustering. **(B)** UMAP visualization of the reclustered cells in control and *Lmna*KO$^{iEC}$ mice. The color of the lasso indicates the cluster of origin for each subclusters. **(C)** Slingshot-based pseudotime trajectory analysis of EC, VSMC, and mVSMC subclusters.

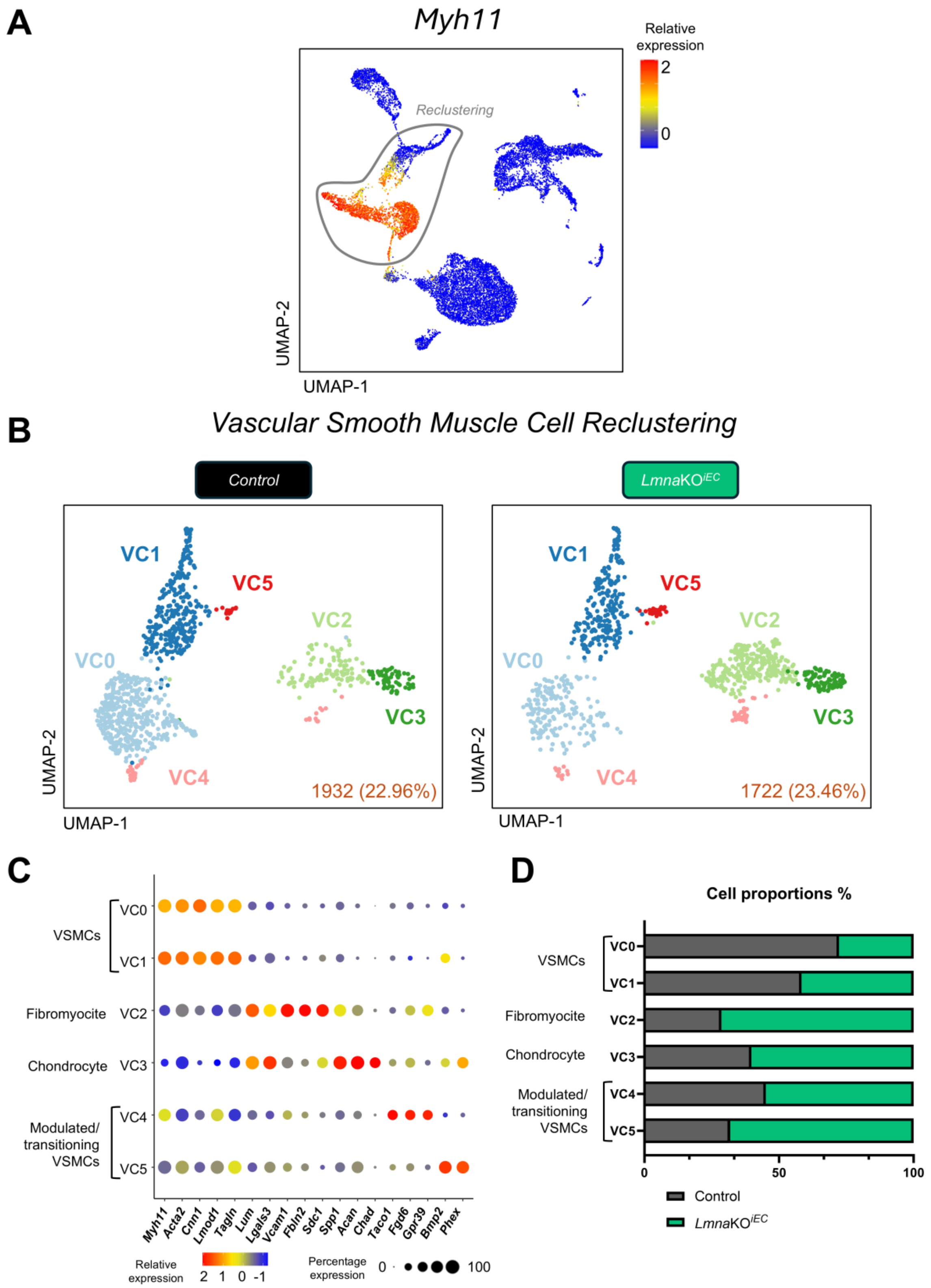


**Supplemental Figure S15. Reclustering and characterization of vascular smooth muscle cell (VSMC) populations from control and *Lmna*KO$^{iEC}$ aortas. (A)** Uniform manifold approximation and projection (UMAP) visualization of the expression of the VSMC marker *Myh11* across all unsupervised clusters. The lasso indicates the cells included in the reclustering. **(B)** UMAP visualization of VSMC reclustering in control and *Lmna*KO$^{iEC}$ mice. **(C)** Dot plot showing the average expression and the percentage of cells expressing selected genes across VSMC subclusters. **(D)** Proportions of cells in each VSMC subcluster by genotype.

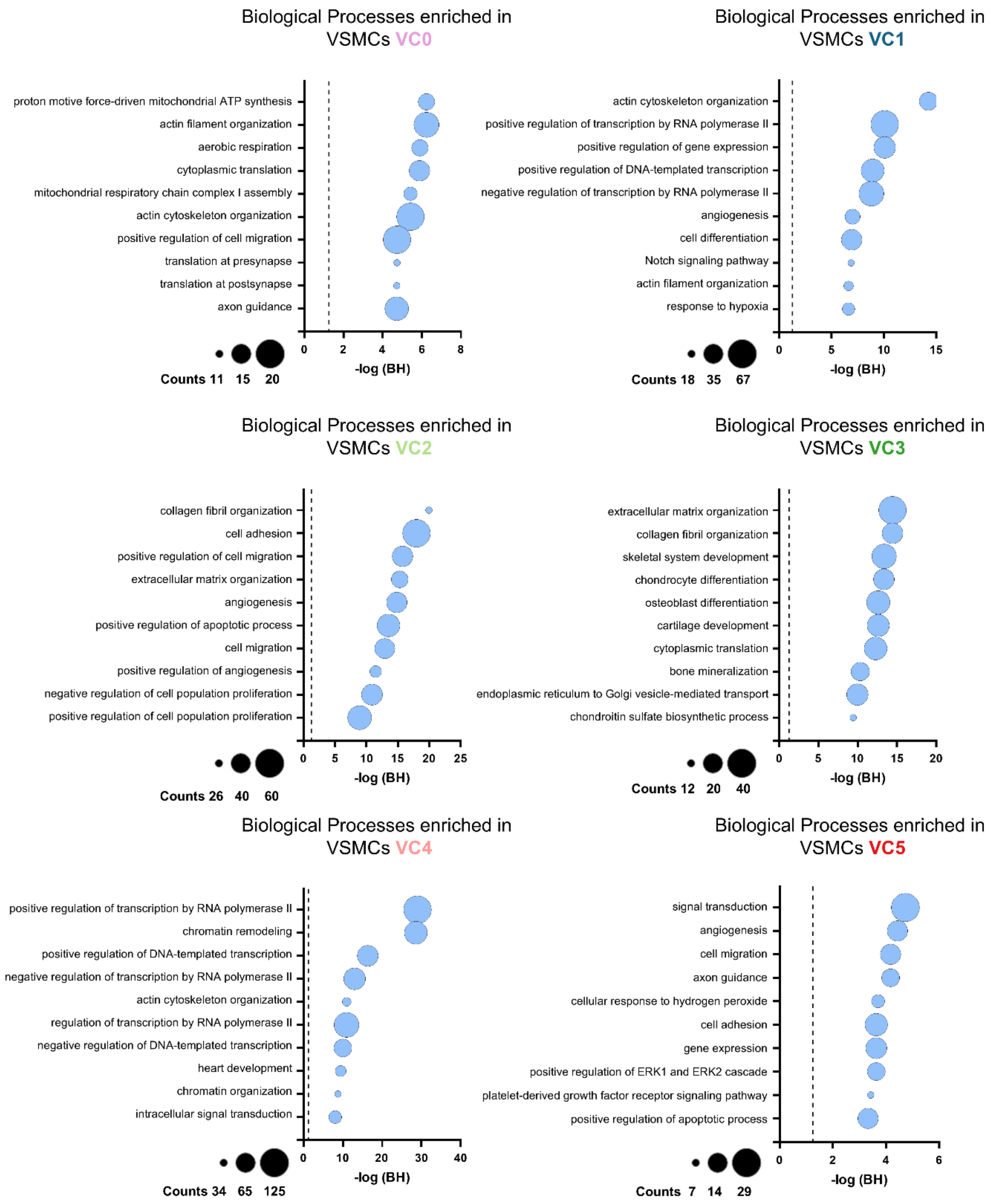


**Supplemental Figure S16. Top 10 enriched Gene Ontology (GO) biological processes for each vascular smooth muscle cell (VSMC) subcluster.** Bubble plots showing the significance and gene count for the top 10 enriched GO biological processes in each VSMC subcluster. Bubble size indicates the number of genes associated with each GO term. The dashed line indicates the Benjamini-Hochberg adjusted P-value threshold of 0.05.

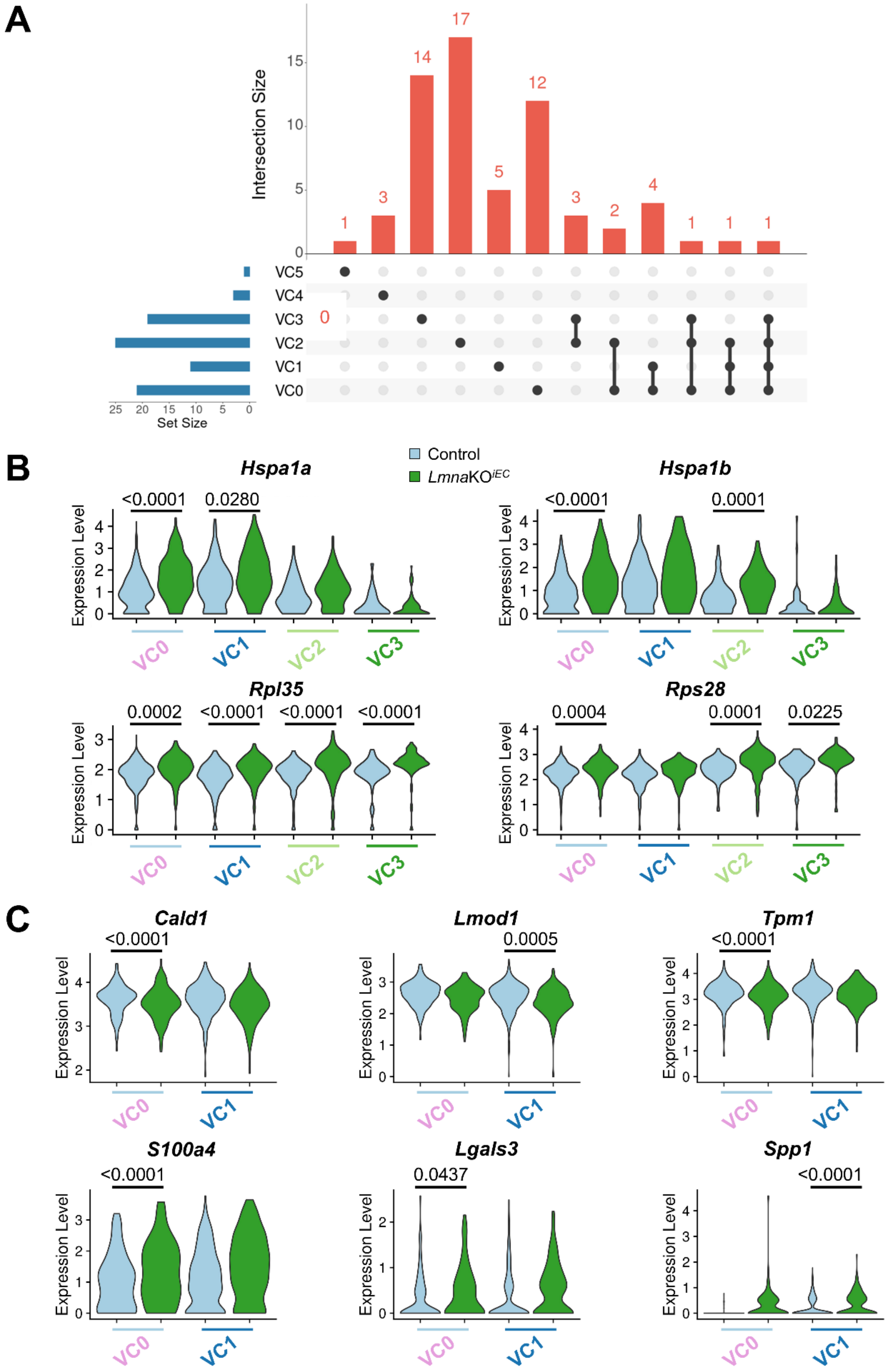


**Supplemental Figure S17. Gene expression changes in aortic vascular smooth muscle cell (VSMC) subclusters from *Lmna*KO$^{iEC}$ mice. (A)** UpSet plot showing the number of differentially expressed genes (DEGs) shared among VSMC subclusters. Only intersections involving two or more subclusters with at least one shared DEG are shown. Set size indicates the total number of DEGs identified in each VSMC subcluster. **(B)** Violin plots showing the expression of selected genes related to stress response and ribosome by genotype across VSMC subclusters. **(C)** Violin plots showing the expression of selected genes related to contractility and the secretory phenotype by genotype in the contractile VSMC subclusters (VC0 and VC1). Statistical analyses were performed using the Wilcoxon rank-sum test. Only significant p values ($p<0.05$) are shown.

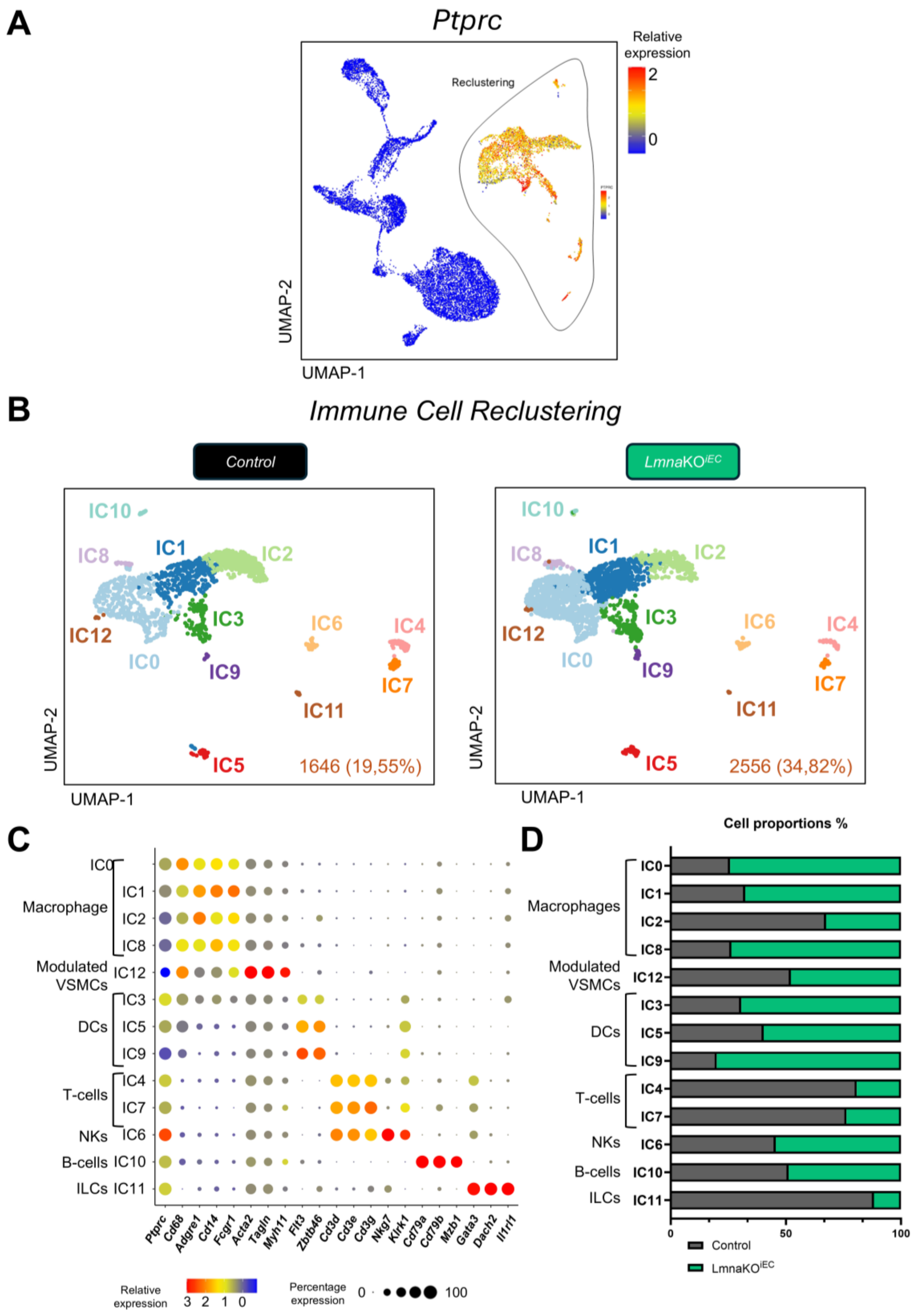


**Supplemental Figure S18. Reclustering and characterization of immune cell populations from control and *Lmna*KO$^{iEC}$ aortas. (A)** Uniform manifold approximation and projection (UMAP) visualization of *Ptprc* (CD45) expression across all unsupervised clusters. The lasso indicates the cells included in the reclustering. **(B)** UMAP visualization of immune cell reclustering in control and *Lmna*KO$^{iEC}$ mice. **(C)** Dot plot showing the average expression and the percentage of cells expressing selected genes across immune cell subclusters. **(D)** Proportions of cells in each immune cell subcluster by genotype. **Abbreviations: VSMC,** vascular smooth muscle cell; **NKs,** natural killer cells, **ILCs,** innate lymphoid cells, **DCs,** dendritic cells.

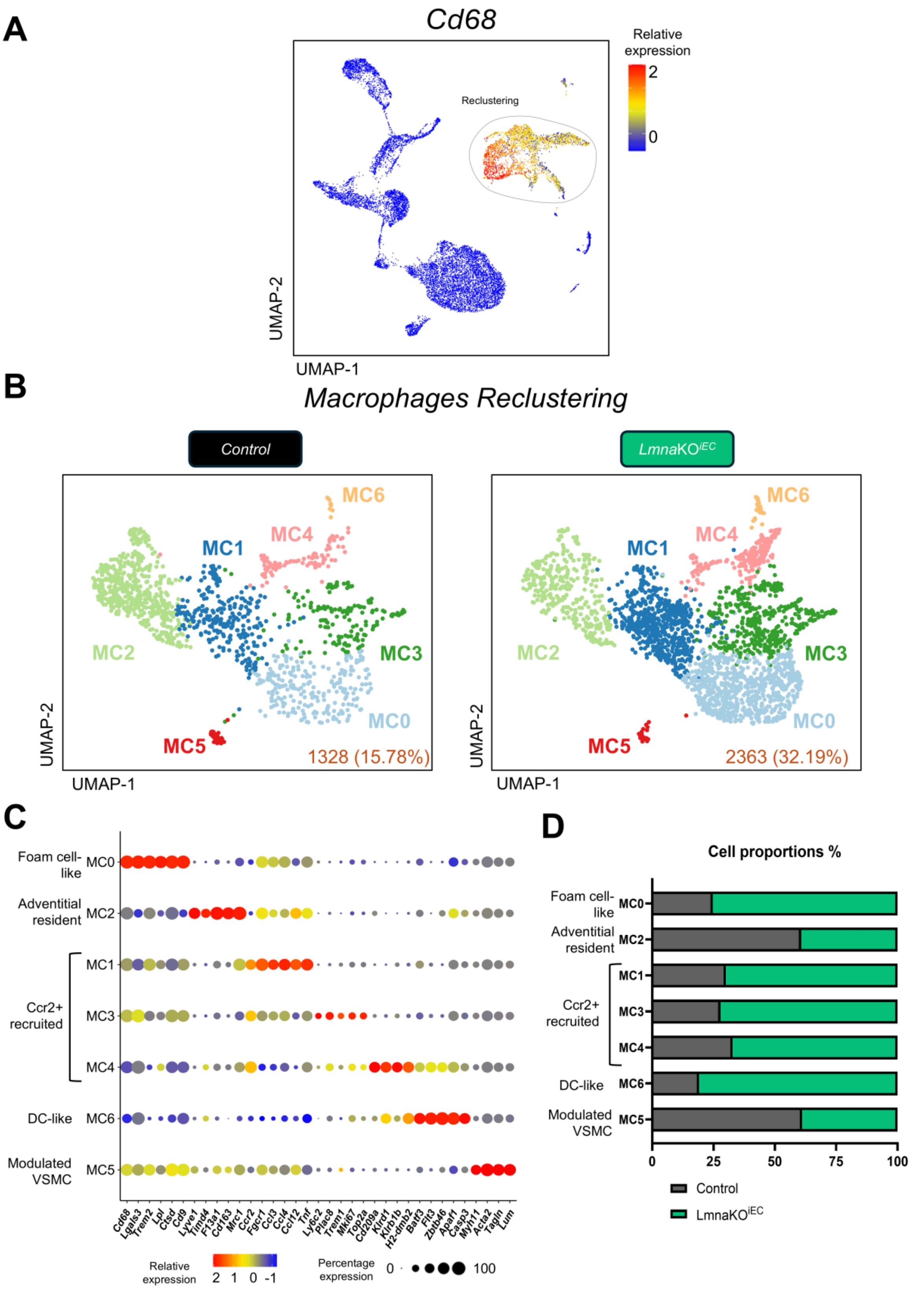


**Supplemental Figure S19. Reclustering and characterization of macrophage populations from control and *Lmna*KO$^{iEC}$ aortas. (A)** Uniform manifold approximation and projection (UMAP) visualization of the expression of the macrophage marker *Cd68* across all unsupervised clusters. The lasso indicates the cells included in the reclustering. **(B)** UMAP visualization of macrophage reclustering in control and *Lmna*KO$^{iEC}$ mice. **(C)** Dot plot showing the average expression and the percentage of cells expressing selected genes across macrophage subclusters. **(D)** Proportions of cells in each macrophage subcluster by genotype. **Abbreviations: VSMC,** vascular smooth muscle cell; **DC,** dendritic cell.

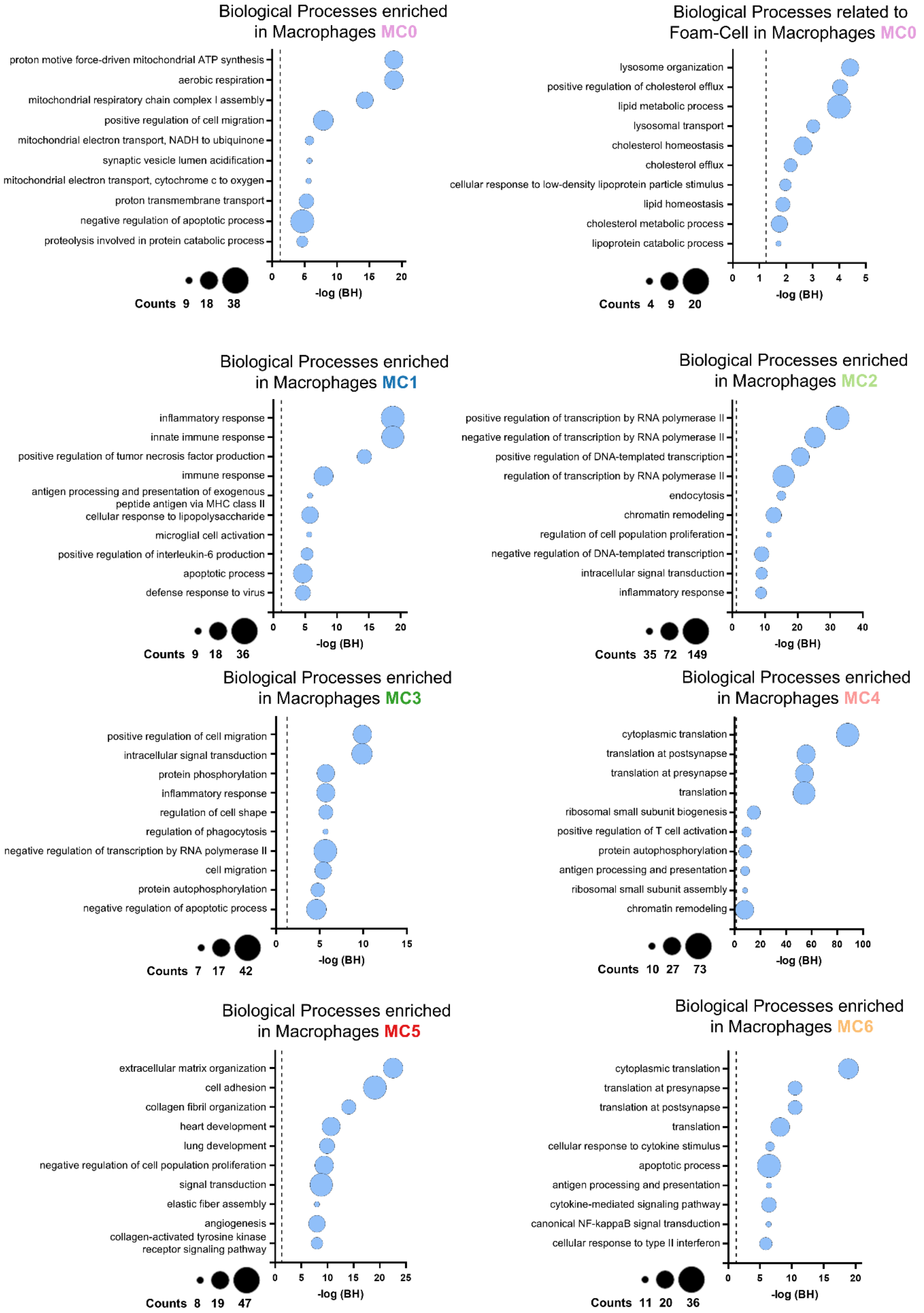


**Supplemental Figure S20. Top 10 enriched Gene Ontology (GO) biological processes for each macrophage subcluster.** Bubble plots showing the significance and gene count for the top 10 enriched GO biological processes in each macrophage subcluster, together with selected GO terms related to foam-cell formation in MC0. Bubble size indicates the number of genes associated with each GO term. The dashed line indicates the Benjamini-Hochberg adjusted P-value threshold of 0.05.

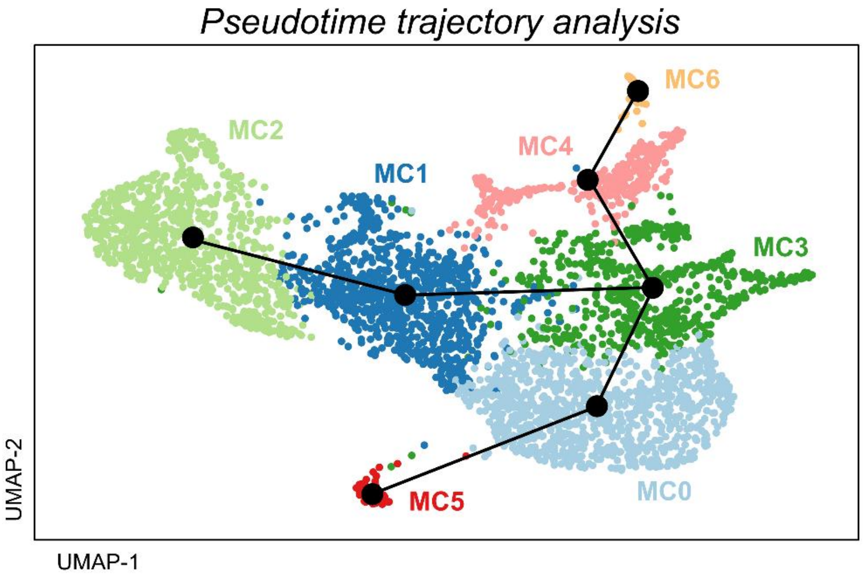


**Supplemental Figure S21. Pseudotime trajectory analysis of macrophage reclustering**. Slingshot-based pseudotime trajectory analysis of macrophage subclusters.

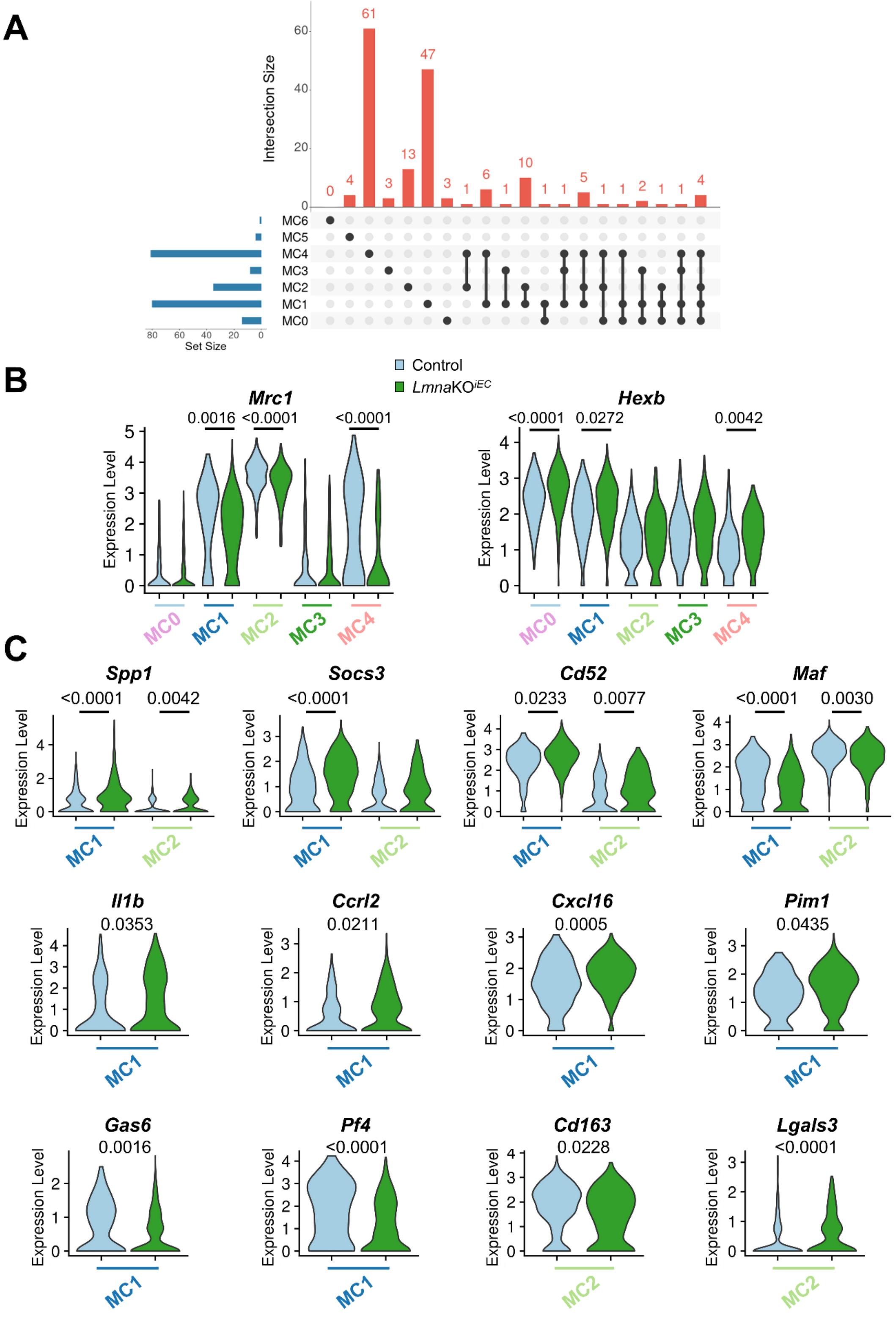


**Supplemental Figure S22. Gene expression changes in aortic macrophage subclusters from *Lmna*KO$^{iEC}$ mice. (A)** UpSet plot showing the number of differentially expressed genes (DEGs) shared among macrophage subclusters. Only intersections involving two or more subclusters with at least one shared DEG are shown. Set size indicates the total number of DEGs identified in each macrophage subcluster. **(B)** Violin plots showing the expression of selected genes across the major macrophage subclusters. **(C)** Violin plots showing the expression of selected genes related to inflammation and macrophage activation by genotype in the inflammatory (MC1) and resident (MC2) macrophage subclusters. Statistical analyses were performed using the Wilcoxon rank-sum test. Only significant p values (p<0.05) are shown.
